# Mathematical Modeling Reveals Quantitative Properties of KEAP1-NRF2 Signaling

**DOI:** 10.1101/2021.08.08.455554

**Authors:** Shengnan Liu, Jingbo Pi, Qiang Zhang

**Affiliations:** Program of Environmental Toxicology, School of Public Health, China Medical University, Shenyang, 110122, China; Gangarosa Department of Environmental Health, Rollins School of Public Health, Emory University, Atlanta, GA, 30322, USA

**Keywords:** Oxidative Stress, KEAP1, NRF2, Ultrasensitivity, Protein Sequestration, Zero-order degradation

## Abstract

In response to oxidative and electrophilic stresses, cells launch an NRF2-mediated transcriptional antioxidant program. The activation of NRF2 depends on a redox sensor, KEAP1, which acts as an E3-ligase adaptor to promote the ubiquitination and degradation of NRF2. While a great deal has been learned about this redox duo, its quantitative signaling properties are still largely unexplored. In the present study, we examined these properties including response time, half-life, maximal activation, and response steepness (ultrasensitivity) of NRF2, through mathematical modeling. The models describe, with increasing complexity, the reversible binding of KEAP1 dimer and NRF2 via the ETGE and DLG motifs, NRF2 production, KEAP1-dependent and independent NRF2 degradation, and perturbations by different classes of NRF2 activators. Our simulations revealed that at the basal condition, NRF2 molecules are largely sequestered by KEAP1, with the KEAP1-NRF2 complex distributed comparably in either an ETGE-bound only (open) state or an ETGE and DLG dual-bound (closed) state, corresponding to the unlatched and latched configurations of the conceptual hinge-latch model. With two-step ETGE binding, the open and closed states operate in cycle mode at the basal condition and transition to equilibrium mode at stressed conditions. Class I-V, electrophilic NRF2 activators, which modify redox-sensing cysteine residues of KEAP1, shift the balance to a closed state that is unable to degrade NRF2 effectively. When total NRF2 accumulates to a level that nearly saturates existing KEAP1, ultrasensitive NRF2 activation, i.e., a steep rise in the free NRF2 level, can occur. The ultrasensitivity results from two simultaneous mechanisms, zero-order degradation mediated by DLG binding and protein sequestration (molecular titration) mediated by ETGE binding. These response characteristics of class I-V activators do not require disruption of DLG binding to unlatch the KEAP1-NRF2 complex. In comparison, class VI NRF2 activators, which directly compete with NRF2 for KEAP1 binding, cause a shift to the unlatched, open state of KEAP1-NRF2 complex and ultimately its complete dissociation (unhinged), resulting in a fast release of free NRF2 followed by stabilization. Although class VI activators may induce free NRF2 to higher levels, ultrasensitivity is lost due to lower free KEAP1 and thus its NRF2-sequestering effect. Stress-induced nuclear NRF2 accumulation is enhanced when basal nuclear NRF2 turnover constitutes a small load of NRF2 production. Our simulation further demonstrated that optimal abundances of cytosolic and nuclear KEAP1 exist to maximize ultrasensitivity. In summary, by simulating the dual role of KEAP1 in repressing NRF2, i.e., sequestration and promoting degradation, our mathematical modeling provides key novel quantitative insights into the signaling properties of the KEAP1-NRF2 system, which may help with the design of novel classes of NRF2 activators and inhibitors and understanding of the toxic actions of environmental oxidative stressors.

## INTRODUCTION

Under oxidative stress, the antioxidant capacity of cells is upregulated to meet the increasing demand for reactive species removal to maintain cellular redox homeostasis and limit cellular damage (Nguyen et al. 2003). Similar to many other cytoprotective responses, this adaptive antioxidant response is underpinned by a complex molecular circuitry of primarily negative feedback and incoherent feedforward nature, involving both posttranslational and transcriptional regulations (Zhang et al. 2010, Zhang et al. 2015). In mammalian cells, the main circuit mediating the transcriptional part of the antioxidant response is the KEAP1-NRF2-ARE pathway (Kobayashi et al. 2009). KEAP1 (Kelch ECH associating protein 1) is the molecular thiol-based sensor of ROS and other reactive species, which detects the redox status inside the cell and relays it to NRF2 (nuclear factor erythroid 2-related factor 2) (Dinkova-Kostova et al. 2002, Suzuki and Yamamoto 2017). As the master transcription factor, NRF2 partners with small Maf (sMaf) proteins to recognize promoter consensus sequences containing AREs (antioxidant response element) and induce a suite of target genes participating in antioxidant and detoxification reactions (Katsuoka et al. 2005, Kobayashi and Yamamoto 2005, Malhotra et al. 2010, Bellezza et al. 2018, Tonelli et al. 2018).

As the essential components for the transcriptional induction of antioxidant genes, the KEAP1 and NRF2 proteins and their interactions have been learned in great details in the past two decades (Itoh et al. 2010, Yamamoto et al. 2018, Paunkov et al. 2019, Baird and Yamamoto 2020). Tethered to the perinuclear actin cytoskeleton in the cytosol, KEAP1 functions as a homodimer (McMahon et al. 2006, Watai et al. 2007, Ogura et al. 2010). The KEAP1 peptide is composed of 624 amino acid residues forming five functional domains: NTR (N-terminal region), BTB (Broad complex, Tramtrack, and Bric-a-Brac), IVR (intervening region), DGR (double glycine repeat) or Kelch-repeat, and CTR (C-terminal region) (Canning et al. 2015, Dayalan Naidu and Dinkova-Kostova 2020). The BTB domain at the N-terminal is responsible for the formation of KEAP1 homodimer (Zipper and Mulcahy 2002). The neighboring Kelch and CTR domains (collectively termed as DC region) are responsible for the interaction of KEAP1 with NRF2 (Li et al. 2004, Lo et al. 2006). As a redox sensor, KEAP1 contains 27 cysteine residues distributed across the five domains, many of which can be modified or conjugated on the thiol group by oxidants and electrophiles (Dinkova-Kostova et al. 2002, Yamamoto et al. 2008, Sekhar et al. 2010).

The NRF2 protein is composed of 589 amino acids forming six functional domains, Neh1 through Neh6 (Moi et al. 1994, Tonelli et al. 2018). The Neh2 domain on the N-terminal is responsible for the binding with the KEAP1 dimer (Itoh et al. 1999). Within Neh2, there exist two conserved motifs in the N-to-C direction: DLG and ETGE, with an intervening sequence containing 7 lysine residues that can be ubiquitinated (Tong et al. 2006, Tong et al. 2007). Both motifs are involved in mediating the association between NRF2 and KEAP1 dimer. The ETGE motif can bind to the DC region of one of the monomeric subunits of KEAP1 dimer, and the DLG motif of the same NRF2 molecule binds to the DC region of the other subunit (McMahon et al. 2006). Therefore, the KEAP1-NRF2 complex exists at an internal molar ratio of 2:1 (Tong et al. 2006, Horie et al. 2021). The binding affinities between ETGE and KEAP1 and between DLG and KEAP1 are substantially different, with ETGE nearly 100-fold higher than DLG (Lo et al. 2006, Tong et al. 2006, Chen et al. 2011, Ichimura et al. 2013, Fukutomi et al. 2014). It is therefore expected that the binding between KEAP1 and NRF2 occurs primarily in two sequential events: an initial ETGE-mediated association forming an “open” KEAP1-NRF2 complex, and a subsequent DLG-mediated intra-complex association forming a “closed” KEAP1-NRF2 complex (Tong et al. 2006).

By interacting with CUL3 (Cullin 3) via its BTB and IVR domains, KEAP1 is an adaptor of the KEAP1-CUL3-RBX1 E3 ubiquitin ligase complex (Kobayashi et al. 2004). When KEAP1 is associated with NRF2 in the closed state, KEAP1 is able to enable the transfer of ubiquitin molecules from the E2-ubiquitin conjugating enzyme bound to RBX1 (RING-box protein 1) to the 7 lysine residues in the intervening region between the DLG and ETGE motifs of NRF2 (Katoh et al. 2005, Tong et al. 2006, Tong et al. 2007). Once ubiquitinated, NRF2 is rapidly degraded by the proteasomal pathway (Kobayashi et al. 2006). Therefore, at basal conditions, NRF2 in the cytosol has a very short half-life, mostly ranging between 6-20 min (Kwak et al. 2002, Alam et al. 2003, Itoh et al. 2003, Stewart et al. 2003, Kobayashi et al. 2004, He et al. 2006, Khalil et al. 2015, Crinelli et al. 2021). Under oxidative stress, certain sensor cysteine residues on KEAP1 are modified, which disables KEAP1’s capability of mediating NRF2 ubiquitination (Yamamoto et al. 2008, Sekhar et al. 2010, Suzuki and Yamamoto 2017). As a result, NRF2 is stabilized and accumulates via *de novo* synthesis in the cytosol. Rising NRF2 then translocates into the nuclei where it induces antioxidant and detoxification genes (Kobayashi and Yamamoto 2005, Itoh et al. 2010, Tonelli et al. 2018).

Despite the molecular details of KEAP1 and NRF2 interactions have been revealed to a great extent, the quantitative signaling properties of the duo, culminating in NRF2 accumulation and nuclear translocation, are still poorly understood. It has been demonstrated that the binding between KEAP1 and NRF2 is not altered by oxidative stress, such that NRF2 does not dissociate from KEAP1 (Eggler et al. 2005, He et al. 2006, Kobayashi et al. 2006). Since the discovery of the two-site sequential binding scheme for KEAP1-NRF2 interaction, i.e., first through ETGE and then through DLG, a hinge-latch model has been proposed (Tong et al. 2006, Tong et al. 2007, Fukutomi et al. 2014). The model considers that the ETGE-mediated association (the hinge) between KEAP1 and NRF2 is always engaged regardless of the presence of oxidative stressors. However, oxidative stressors may disrupt the weaker DLG-mediated association (the latch), rendering the closed KEAP1-NRF2 complex to revert to the open configuration (McMahon et al. 2006, Ogura et al. 2010). In the open state, KEAP1 can no longer mediate the ubiquitination of NRF2, resulting in NRF2 stabilization. However, the validity of the hinge-latch model for KEAP1 cysteine-modifying, electrophilic oxidants (i.e., class I-V NRF2 inducers) becomes questionable as emerging evidence suggests that these classes of compounds do not disrupt DLG binding (Horie et al. 2021). Studies using Förster resonance energy transfer (FRET) revealed that the association between KEAP1 and NRF2 may become even stronger when cells are exposed to KEAP1 cysteine-modifying compounds (Baird et al. 2013).

The ETGE-mediated binding affinity between KEAP1 and NRF2 is high relative to their cellular abundances, with the dissociation constant (*K_d_*) ranging between 5-26 nM, as summarized in Table S1 footnote (Lo et al. 2006, Tong et al. 2006, Chen et al. 2011, Ichimura et al. 2013, Fukutomi et al. 2014), and the intracellular concentrations of KEAP1 dimer and NRF2 in the order of hundreds of nM as observed in a variety of cell types (Iso et al. 2016). This suggests that when KEAP1 is in excess relative to NRF2, as often the case at the basal condition, NRF2 molecules are largely sequestered by KEAP1, leaving free NRF2 only a very small fraction of its total abundance. Such binding kinetics suggests that under oxidative stress, newly synthesized NRF2 molecules will be still first sequestered by the remaining free KEAP1 reserve, and only when it is nearly all filled by NRF2, will NRF2 becomes more available for nuclear translocation. Therefore, the degree of NRF2 activation is in part regulated by the KEAP1 reserve capacity of NRF2 sequestration. This mode of NRF2 activation is recently suggested in the floodgate hypothesis (Iso et al. 2016, Suzuki and Yamamoto 2017, Yamamoto et al. 2018). If total NRF2 never accumulates to a level that can saturate existing KEAP1 molecules, nuclear NRF2 translocation and gene induction will remain muted. However, if total NRF2 can rise to a higher level that nearly saturates KEAP1, from a quantitative signaling prospective, KEAP1-dependent NRF2 degradation will operate near zero order and simultaneously NRF2 begins to escape KEAP1 sequestration, both of which are robust ultrasensitive mechanisms that can produce a steep rise in free NRF2 levels (Buchler and Louis 2008, Zhang et al. 2013, Ferrell and Ha 2014). This amplified, nonlinear NRF2 activation can in turn induce antioxidant genes strongly. Therefore, the kinetic parameters governing the interactions between KEAP1 dimer and NRF2 seem to be critical to the quantitative behaviors of KEAP1-NRF2-ARE-mediated redox signal transduction.

From the perspective of effectively restoring redox homeostasis, the induction of these antioxidant genes needs to be launched timely and to levels that are sufficient to counteract the oxidative impacts exerted by the stressors (Zhang et al. 2010). Strong antioxidant induction would require signal amplification, i.e., ultrasensitivity, by which a small percentage change in the redox status can be transduced to induce a larger percentage change in the expression of antioxidant genes (Zhang et al. 2013, Ferrell and Ha 2014). a number of ultrasensitive mechanisms, including multistep signaling, homomultimerization, and autoregulation, have been revealed in the KEAP1-NRF2-ARE mediated transcriptional pathway (Zhang and Andersen 2007, Zhang et al. 2010). They operate collectively to ensure that the cellular antioxidant capacity can be adequately induced to levels matching the intensity of the oxidant insult.

Mathematical modeling plays a crucial role in understanding and predicting the quantitative behavior of redox pathways (Adimora et al. 2010, Selvaggio et al. 2018). Earlier modeling work including our own has included the KEAP1-NRF2 module in the larger context of the NRF2-mediated antioxidant response pathways (Zhang and Andersen 2007, Zhang et al. 2009, Hamon et al. 2014, Leclerc et al. 2014, Khalil et al. 2015, Xue et al. 2015, Kolodkin et al. 2020). However, in most of these studies the KEAP1-NRF2 module was treated as simplified degradation network motifs, yet the details of KEAP1-NRF2 interactions and especially the likely nonlinearity in signaling have not been explicitly and fully explored. In the present study, we developed a suite of mathematical models of detailed KEAP1-NRF2 interactions to explore the quantitative properties of NRF2 activation. With these models we examined the roles of open and closed states of the KEAP1-NRF2 complex for the hinge-latch and floodgate hypotheses. Our simulation predicts that ultrasensitive NRF2 activation may occur via zero-order protein degradation and protein sequestration by KEAP1 under certain circumstances. Our mathematical models provide key quantitative insights into the signaling properties of the KEAP1-NRF2 module of the adaptive, cellular antioxidant response pathway.

## METHODS

### Model structure

In keeping with the principle of parsimony and exploring the importance of molecular details, we started with a minimal model capturing the basic interactions between KEAP1 and NRF2, and we then progressively built more complexity into the model based on more recent quantitative knowledge about the interactions. As a result of this evolution, a total of 6 models were explored with increasing complexities, as summarized in Table 1. For all models, the following assumptions were made.

i. KEAP1 is treated as a single molecule of homodimer with two binding sites for NRF2 as the dimer structure is required for NRF2 binding (Zipper and Mulcahy 2002).
ii. Total KEAP1 abundance is a constant which is not altered by oxidative stress as extensively demonstrated in experimental studies (Iso et al. 2016) and KEAP1 turnover (synthesis and degradation) is not considered.
iii. Since the binding affinity between KEAP1 and the ETGE motif of NRF2 is much higher than the binding affinity between KEAP1 and the DLG motif of NRF2 (> 100-fold), as summarized in Table S1 (Lo et al. 2006, Tong et al. 2006, Chen et al. 2011, Ichimura et al. 2013, Fukutomi et al. 2014), for simplicity and following the concept of hinge-latch hypothesis (Yamamoto et al. 2018), the initial interaction between KEAP1 and NRF2 is assumed to always start with the binding between KEAP1 and ETGE while the binding between KEAP1 and DLG occurs subsequently, as an intramolecular event.
iv. Oxidation or conjugation of one monomeric subunit of the KEAP1 dimer by a class I-V NRF2 activator is sufficient to cause KEAP1 to lose its ability to mediate NRF2 degradation. The oxidation or conjugation can occur to either free KEAP1 dimer or KEAP1 complexed with NRF2 equally.
v. For the Models (4a and 4b) with nuclear NRF2 translocation, cytosolic KEAP1 and nuclear KEAP1 are kept as separate pools.

Model 1 is the most basic model, which captures the known essence of interactions between KEAP1 and NRF2 in the cytosol as shown in Fig. 1. In model 1, NRF2 is synthesized at a constant rate of *k*_0_. Free NRF2 (*NRF2*_free_) is degraded with a first-order rate constant of *k*_5_, reflecting KEAP1-independent degradation such as the one mediated by the Neh6 domain involving the GSK-3, β-TrCP and Cul1 system (McMahon et al. 2004, Rada et al. 2011, Chowdhry et al. 2013, Hayes et al. 2015). *NRF2*_free_ first binds to one of the monomeric subunits of the KEAP1 dimer through the ETGE domain with a second-order association rate constant *k*_1_ and a first-order dissociation rate constant *k*_2_, forming an intermediate complex *KEAP1_NRF2*_open_ (termed open state here). Since KEAP1 in the open state of the complex cannot execute its E3 ligase adaptor function (Katoh et al. 2005, Tong et al. 2006, Tong et al. 2007), NRF2 in *KEAP1_NRF2*_open_ is assumed to be degraded with a first-order rate constant of *k*_9_ that is equal to *k_5_*. As NRF2 is degraded, KEAP1 is recycled joining the free KEAP1 dimer pool. The NRF2 molecule in *KEAP1_NRF2*_open_ then further associates with the other unoccupied monomeric subunit of KEAP1 dimer through the DLG motif with a first-order association rate constant *k*_3_ and a first-order dissociation rate constant *k*_4_, forming the final complex *KEAP1_NRF2*_closed_ (termed closed state here). NRF2 in *KEAP1_NRF2*_closed_ is degraded with a first-order rate constant of *k*_6_ which is much higher than *k*_5_ and *k*_9_, reflecting KEAP1-mediated ubiquitination and accelerated degradation of NRF2, and KEAP1 dimer is recycled. Class I-V oxidants and electrophiles can oxidize or conjugate KEAP1 (Yamamoto et al. 2008, Sekhar et al. 2010, Suzuki and Yamamoto 2017). In the model the oxidant converts *KEAP1* to an oxidized form, *KEAP1*_o_, with a second-order rate constant *k*_7_. The same oxidation reaction is assumed to occur on the KEAP1 molecule in *KEAP1_NRF2*_open_ and *KEAP1_NRF2*_closed_ as well, forming *KEAP1*_o_*_NRF2*_open_ and *KEAP1_o__NRF2*_closed_ respectively. *KEAP1*_o_, *KEAP1*_o_*_NRF2*_open_, and *KEAP1*_o_*_NRF2*_closed_ can be reduced back to the respective original states with a first-order rate constant *k*_8_. Since there is no evidence that the association of KEAP1 with NRF2 alters the kinetics of oxidation or conjugation of KEAP1 by oxidants, the same values of *k*_7_ and *k*_8_ are used across all three oxidation/reduction reaction pairs. Since the alteration of NRF2 stability only occurs in the closed state, NRF2 in *KEAP1*_o_*_NRF2*_open_ is degraded with a first-order rate constant of *k’*_9_ that is equal to *k*_9_. NRF2 in *KEAP1*_o_*_NRF2*_closed_ is degraded with a first-order rate constant of *k’*_6_ that is much lower than *k*_6_, reflecting the well-established fact that oxidant-modified KEAP1 in the closed state loses its capability to mediate the ubiquitination and degradation of NRF2 (Katoh et al. 2005, Tong et al. 2006, Tong et al. 2007). In both the *k’*_9_ and *k’*_6_ steps, *KEAP1*_o_ is recycled joining the free *KEAP1*_o_ pool. The binding between *NRF2*_free_ and *KEAP1*_o_ through the ETGE domain is described by the second-order association rate constant *k’*_1_ and first-order dissociation rate constant *k’*_2_, which are kept the same as *k*_1_ and *k*_2_ respectively since class I-V oxidants do not alter the binding affinity between KEAP1 and NRF2 (Eggler et al. 2005, He et al. 2006, Kobayashi et al. 2006). The association and dissociation rate constants *k’*_3_ and *k’*_4_ for the intramolecular DLG binding between *KEAP1*_o_*_NRF2*_open_ and *KEAP1*_o_*_NRF2*_closed_ are also kept the same as *k*_3_ and *k*_4_ respectively, however, their values are varied to explore the behavior of the hinge-latch hypothesis.

**Table 1.** KEAP1-NRF2 Model Features.

| Model # | Model structure | Cycle mode of operation | Two-step ETGE binding | NRF2 nuclear translocation | Class of NRF2 activator |
| --- | --- | --- | --- | --- | --- |
| 1 | Fig. 1 |  |  |  | I-V |
| 2 | Fig. 1 | X |  |  | I-V |
| 3a | Fig. 2A | X | X |  | I-V |
| 3b | Fig. 6A | X | X |  | VI |
| 4a | Fig. 7A | X | X | X | I-V |
| 4b | Fig. 8A | X | X | X | VI |

Note: X denotes that a model has the corresponding feature. The structure of each model is illustrated in the Figures indicated.

**Figure 1.**
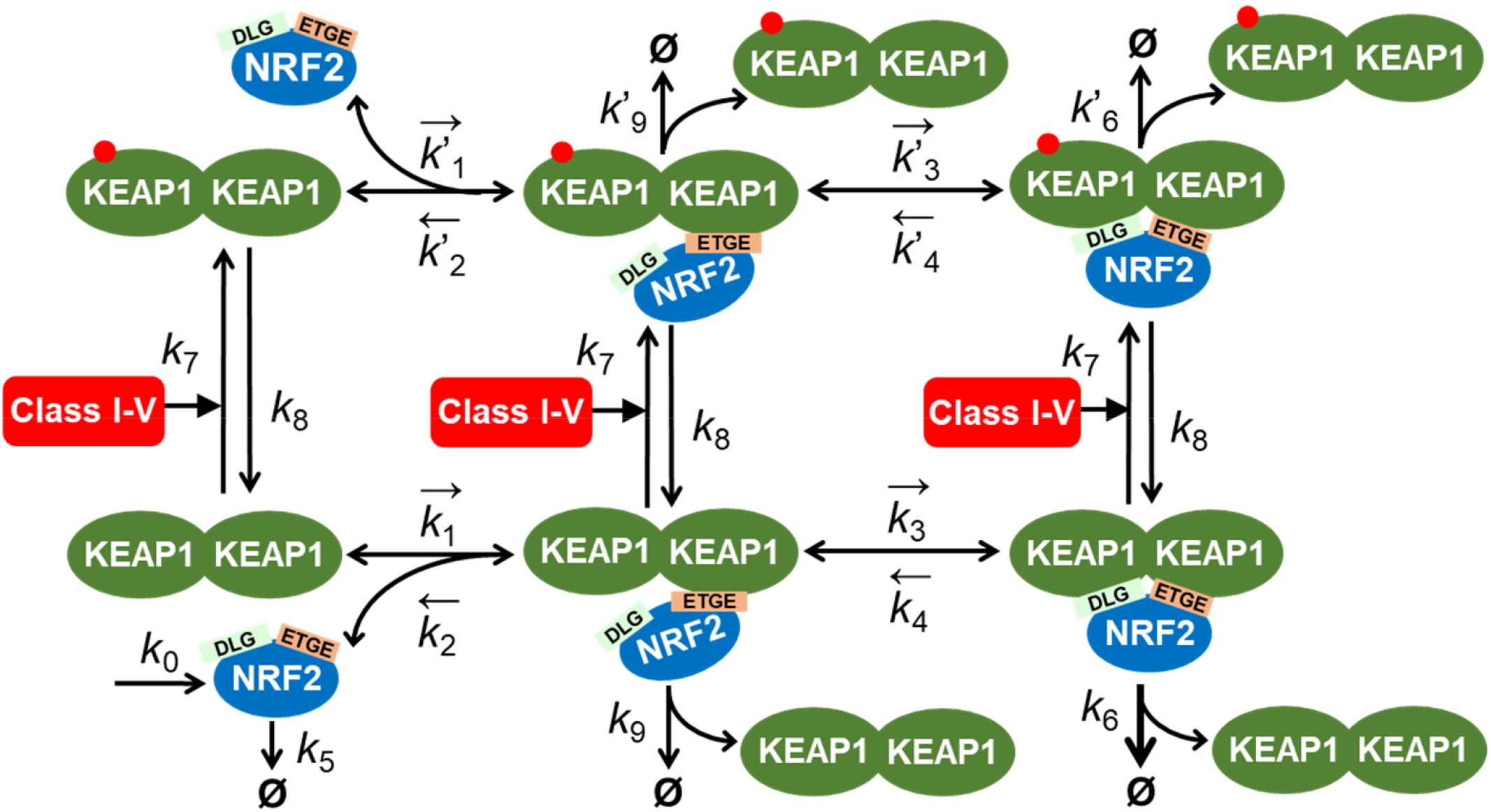
Structure of KEAP1-NRF2 Models 1 and 2. The models feature one-step ETGE binding and interaction with class I-V activator. Short arrow bars next to a parameter symbol denote the direction of the reversible binding described by the parameter. Φ denotes degradation. These denotations apply to all other model structures.

The detailed structure of Models 2, 3a, 3b, 4a, and 4b are presented in Figs. 1, 2A, 6A, 7A, and 8A, respectively. Briefly, in Model 2, the DLG-mediated internal binding kinetics (*k*_3_ and *k*_4_) between *KEAP1_NRF2*_open_ and *KEAP1_NRF2*_closed_ is modified from Model 1 to simulate the situation that the transitioning between the two states occurs in a cycle mode rather than an equilibrium mode, as observed experimentally (Baird et al. 2013). In Models 3a and 3b, the ETGE-mediated binding between KEAP1 and NRF2 is modified from the one-step mode as in Models 1 and 2 to a two-step mode to simulate the situation that ETGE-mediated binding involves an initial fast binding event followed by a subsequent slow binding event observed experimentally (Fukutomi et al. 2014). This modification allows us to achieve the cycle mode of operation without altering the DLG-mediated binding kinetics dramatically as done in Model 2. Models 3a and 3b consider class I-V and VI NRF2 inducers as separate cases respectively. Lastly, in Models 4a and 4b, translocation of NRF2 to the nucleus and its interaction with KEAP1 in the nucleus are considered, and the two models consider class I-V and VI NRF2 activators as separate cases respectively.

### Model parameters and ordinary differentiation equations (ODEs)

The values of most of the model parameters, including binding rate constants, degradation rate constants, and abundances (concentrations) of KEAP1 and NRF2, were obtained or derived from the literature. For those unknown parameter values, they were estimated based on other constraints of the modeled system. References and details of the determination and calculation of all parameter values are presented in Table S1 and its footnote. The unit of concentration of the state variables is nM and time is second (S). The ODEs are presented in Tables S2-S6 and algebraic equations calculating the concentrations of state variables in various combinations are presented in Table S7. The steady-state concentrations of state variables at the basal and maximally induced conditions are in Tables S8 and S9 respectively, and the steady-state turnover fluxes of reactions at the basal and maximally induced conditions are in Tables S10 and S11 respectively.

### Modeling tools

The models were constructed and simulated in Berkeley Madonna (version 8.3.18, University of California, Berkeley, CA) using the “Rosenbrock (stiff)” ODE solver. All model codes in Berkeley Madonna format as well as in *R* format are available as additional Supplemental files.

### Metrics of ultrasensitivity

In the present study, all oxidant-NRF2 dose-response (DR) curves were obtained once the simulation has achieved steady state. The degree of ultrasensitivity of a steady-state DR curve can be evaluated with two related metrics. First, the Hill coefficient, *n_H_*, is approximated from the equation

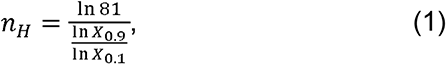

where *X*_0.9_ and *X*_0.1_ are the concentrations of an oxidant that produce 90% and 10% respectively of the maximal NRF2 response (after subtracting the basal NRF2 levels) (Zhang et al. 2013). *n_H_* represents the overall steepness or global degree of ultrasensitivity of the DR curve. Second, we evaluate the local response coefficient (*LRC*) of a DR curve by calculating all slopes of the curve on dual-log scales, which are equivalent to the ratios of the fractional change in response (*R*) to the fractional change in dose (*D*) (Goldbeter and Koshland 1982):

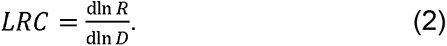

The maximal |*LRC*| of a DR curve, *LRC_max_*, represents the maximal amplification capacity of KEAP1-NRF2-mediated signaling. Typical ultrasensitive responses have *LRC_max_* values substantially above 1. The comparison between *n_H_* and *LRC* is important as these quantities are not necessarily equivalent and depend on the basal response level and the shape of the DR curve; thus, *n_H_* alone can misrepresent the actual degree of signal amplification (Legewie et al. 2005, Zhang et al. 2013, Altszyler et al. 2017).

## Results

### Models 1 and 2

Following the principle of parsimony, we started with the minimal model structure for Models 1 and 2, which is depicted in Fig. 1 and detailed in Methods. In-depth analyses of the basic properties and response behaviors of the two models are provided in the Supplemental Material. Simulations of these two models provided the initial findings that the KEAP1-NRF2 interaction is capable of producing ultrasensitive activation of free NRF2 by class I-V compounds (Figs. S1C, S1D, and S3F), and the ultrasensitivity results from saturation of KEAP1 by NRF2 where both zero-order degradation and protein sequestration of NRF2 occur (Fig. S3G-S3H).

In Model 1, however, the abundance ratio of the open and closed KEAP1-NRF2 complexes (*KEAP1_NRF2*_open_:*KEAP1_NRF2*_closed_) remains constant at 1:1 at all times in all conditions (Figs. S1A-S1C and S2), indicating these two states are always at equilibrium to each other. Using FRET to track the open and closed states of KEAP1-NRF2 complex, Baird et al. observed that the two states diverge and do not follow an equilibrium mode of operation in a variety of chemically perturbed conditions (Baird et al. 2013). But rather, a “cyclic sequential attachment and regeneration” (abbreviated as “cycle”) mode of operation was suggested. In this mode, because of the rapid degradation of NRF2 in the closed KEAP1-NRF2 complex, KEAP1 is quickly released (or regenerated) to join the free KEAP1 dimer pool and able to sequester newly synthesized NRF2 again, thus completing a global cycle for KEAP1. Under oxidative stress, this cycle is blocked as the NRF2 degradation-coupled release of KEAP1 from the closed KEAP1-NRF2 complex is inhibited, leading to accumulation of the closed state and depletion of free KEAP1 dimer. Flux analysis indicated that Model 1 operates in equilibrium mode because of the much higher association and dissociation fluxes through the DLG-binding step than the connected NRF2 turnover fluxes (Tables S10-S11). We thus evolved Model 1 into Model 2 by dramatically reducing the parameter values of *k*_3_ and *k*_4_ (the rate constants for DLG binding), which indeed successfully made Model 2 behave in a cycle mode with divergent responses of the open and closed KEAP1-NRF2 complexes (Figs. S3A-S3B, and S4). However, this “success” was achieved by setting *k*_4_, the dissociation rate constants for DLG binding, to a value that is hundreds-fold lower than experimentally measured (Fukutomi et al. 2014). Importantly, in the same study, it was also demonstrated that the first binding event, i.e., between KEAP1 and the ETGE motif of NRF2, is a thermodynamically two-step process, involving an initial fast binding step to form a transient, intermediate complex (termed *KEAP1_NRF2*_open1_ here) first, followed by a much slower second step that leads to a more stable configuration of the open complex (termed *KEAP1_NRF2*_open*2*_ here). As a minimal model, Models 1 and 2 only considered single-step ETGE binding. We hypothesized that this second, slow ETGE-binding step may account for the experimentally observed cycle mode of operation. We next set out to test this hypothesis with Model 3a.

### Model 3a (Two-Step ETGE-Binding Cycle Mode for Class I-V activator)

In Model 3a we added an extra, reversible step, *k*_1.1_ and *k*_2.1_, to account for the intramolecular state transition between *KEAP1_NRF2*_open1_ and *KEAP1_NRF2*_open*2*_ (Fig. 2A), with *k*_4_ restored to the high value measured in (Fukutomi et al. 2014). As detailed in Table S1 footnote, we then iteratively adjusted the values of *k*_0_, *k*_3_ and *k_6_* such that the basal *NRF2*_tot_ level is still at 150 nM and half-life at 10 min, and the basal open (*KEAP1_NRF2*_open_ *= KEAP1_NRF2*_open1_ *+ KEAP1_NRF2*_open2_):closed (*KEAP1_NRF2*_closed_) state ratio remains at 1:1 (Fig. 2B). NRF2 in *KEAP1_NRF2*_open1_ and *KEAP1_NRF2*_open2_ were assumed to degrade with the same rate constants (*k*_9_=*k*_9.1_=*k*_5_).

**Figure 2.**
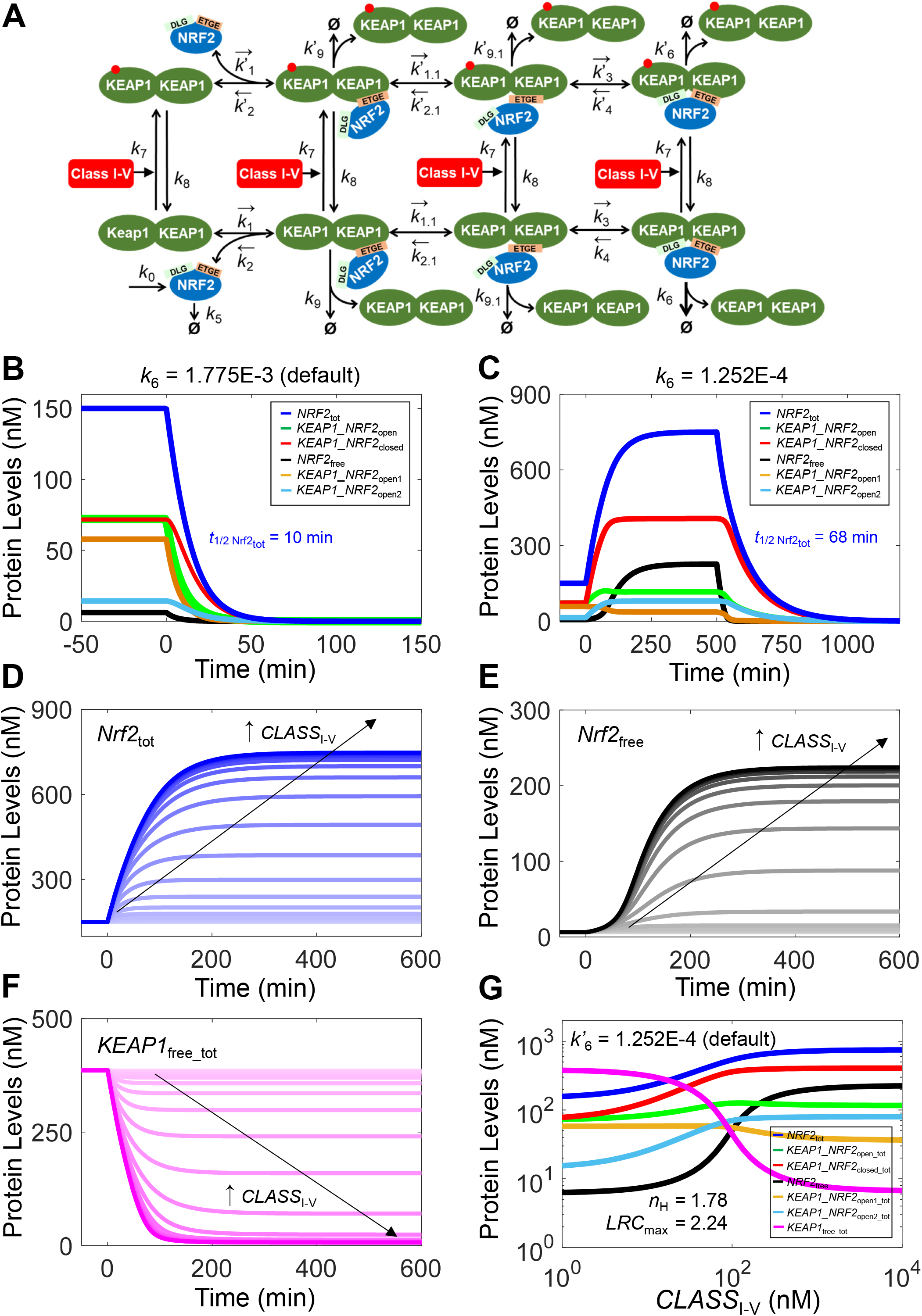
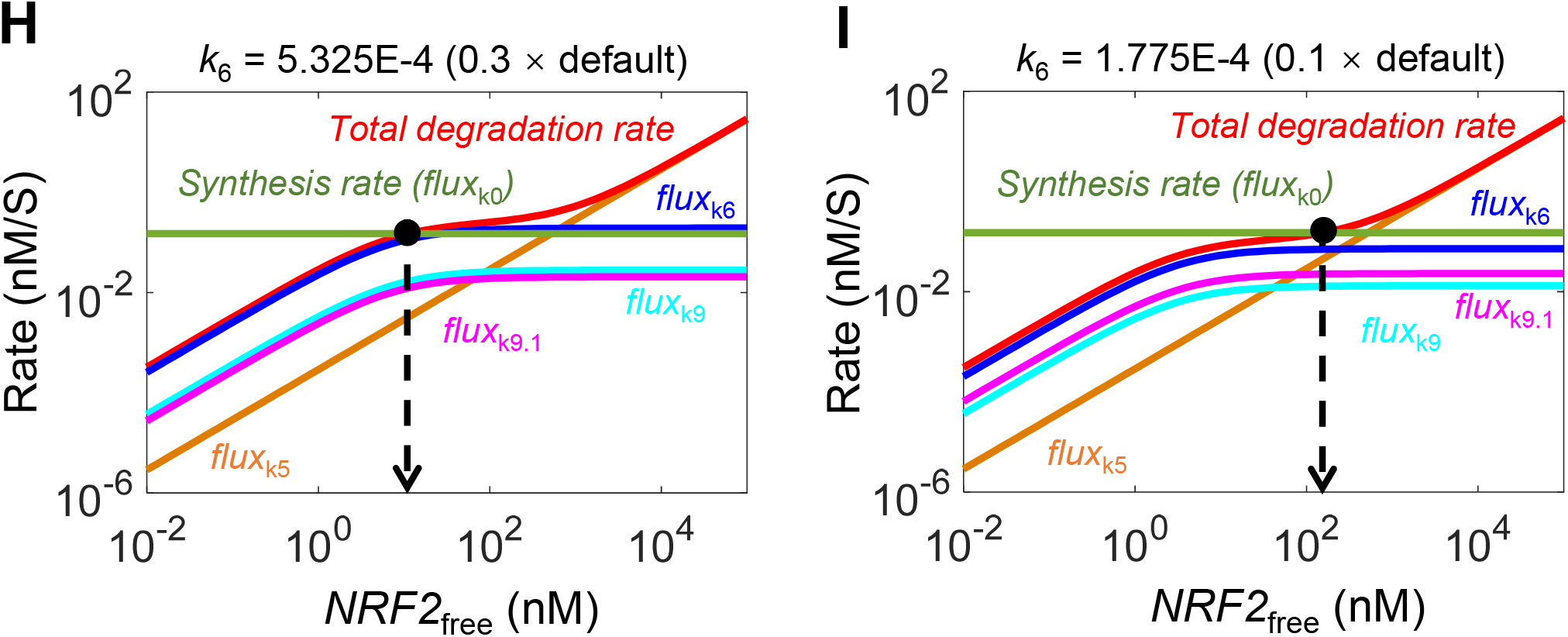
Structure, and dynamical and steady-state behaviors of Model 3a. **(A)** Structure of Model 3a featuring two-step ETGE binding and interaction with class I-V activator. **(B)** Dynamical changes of basal *NRF2*_free_, *KEAP1_NRF2*_open1_, *KEAP1_NRF2*_open2_, *KEAP1_NRF2*_closed_, and *NRF2*_tot_ in response to termination of NRF2 synthesis (by setting *k*_0_=0) starting at 0 min with *k*_6_ at default value. **(C)** Dynamical changes of various NRF2 species in response to stabilization of NRF2 in *KEAP1_NRF2*_closed_ by setting *k*_6_=1.252E-4 starting at 0 min and in response to termination of NRF2 synthesis (by setting *k*_0_=0) starting at 500 min. For simulations in (B) and (C), *CLASS*_I-V_ level is kept at zero. Dynamical changes of **(D)** *NRF2*_tot_, **(E)** *NRF2*_free_, and **(F)** *KEAP1*_free_tot_ in response different levels of *CLASS*_I-V_ with *k’*_6_ at default value. **(G)** Steady-state dose-response curves of various NRF2 species and *KEAP1*_free_tot_ on double-log scale with *k’*_6_ at default value. Shown are *n*_H_ and *LRC*_max_ for *NRF2*_free_; *n*_H_ and *LRC*_max_ for *NRF2*_total_ are 1.22 and 0.42 respectively (not shown). **(H-I)** Flux analyses for conditions when NRF2 in *KEAP1_NRF2*_closed_ is stabilized by setting *k*_6_ to 30% (H) and 10% (I) of default value.

At the basal condition, with the default parameter setting of Model 3a, the second step of ETGE binding (*k*_1.2_ and *k*_2.1_) does not operate in equilibrium mode. This is because *flux*_k1.1_ = 0.133 and *flux*_k2.1_ = 1.73E-3 nM/S, thus only a tiny fraction of *KEAP1_NRF2*_open2_ is returned to *KEAP1_NRF2*_open1_ (Table S10). Another small fraction is degraded through *flux*_k9.1_ at 4.1E-3 nM/S. Over 95% of *KEAP1_NRF2*_open2_ is moved forward to become *KEAP1_NRF2*_closed_ at a net flux (*flux*_k3_ - *flux*_k4_) of 0.127 nM/S. In contrast, both the first step of ETGE binding (*k*_1_ and *k*_2_) and the step of DLG binding (*k*_3_ and *k*_4_) operate in equilibrium mode, with *flux*_k1_ and *flux*_k2_ at 16.48 and 16.33 nM/S respectively, and *flux*_k3_ and *flux*_k4_ at 14.19 and 14.07 nM/S respectively, all of which are >100 fold higher than other connected turnover fluxes (Table S10). As a result, the *NRF2*_free_:*KEAP1_NRF2*_open1_ ratio, which is 1:9.4, is largely determined by the *k*_2_:(2\**k*_1_\**Keap1*_free_) ratio, and the *KEAP1_NRF2*_open2_:*KEAP1_NRF2*_closed_ ratio, which is 1:5, is largely determined by the *k*_4_:*k*_3_ ratio. At 58 and 14 nM respectively, *KEAP1_NRF2*_open1_ dominates *KEAP1_NRF2*_open*2*_, accounting for 80% of the total open KEAP1-NRF2 complex (Fig. 2B and Table S8).

When setting *k*_0_=0 to examine the decay of NRF2 species from their basal steady states, *NRF2*_free_:*KEAP1_NRF2*_open1_ and *KEAP1_NRF2*_open2_:*KEAP1_NRF2*_closed_ remain at the same equilibrium ratios as above as all NRF2 species decrease (Fig. 2B). *NRF2*_free_ and *KEAP1_NRF2*_open1_ decrease quickly with a half-life of about 4-5 min, due primarily to the depletion of *KEAP1_NRF2*_open1_ through *flux*_k1.1_, which is about 8-fold greater than *flux*_k9_. In contrast, *KEAP1_NRF2*_open2_ and of *KEAP1_NRF2*_closed_ do not decrease as fast because of the continued supply of KEAP1-NRF2 complex through *flux*_k1.1_. Because of the differential decay rates, the relative abundance of *KEAP1_NRF2*_open1_ and *KEAP1_NRF2*_open2_ switches positions over time, with *KEAP1_NRF2*_open2_ becoming the dominant form of the open KEAP1-NRF2 complex eventually. Furthermore, the levels of *KEAP1_NRF2*_open_ and *KEAP1_NRF2*_closed_ diverge quickly from the basal ratio of 1:1 to 1:2.5 by 15 min, and to about 1:4.5 eventually. Since toward the end of the decay process *KEAP1_NRF2*_open2_ is the dominant form of the open KEAP1-NRF2 complex, this 1:4.5 ratio closely reflects the equilibrium ratio of *KEAP1_NRF2*_open2_:*KEAP1_NRF2*_closed_, which is determined primarily by the *k*_4_:*k*_3_ ratio.

To examine the behavior of Model 3a when NRF2 in *KEAP1_NRF2*_closed_ is stabilized, we first lowered *k*_6_ to different values, while keeping *CLASS*_I-V_=0. As *k*_6_ decreases from the default 1.775E-3 S^-1^ (equivalent *t*_1/2_=6.5 min) to 1.252E-4 (which is the default value of *k’*_6_, equivalent *t*_1/2_=92 min), all NRF2 species (except *KEAP1_NRF2*_open1_) increase and reach steady states in about 400 min (Fig. 2C). The open:closed ratio decreases and reaches about 1:2.8 at 1 h, and settles to 1:3.5 at steady state, with *KEAP1_NRF2*_open2_ switching to the dominant form of the open-state complex. When reaching steady states, *NRF2*_tot_ increases by 5-fold, while *NRF2*_free_ increases by a much greater fold, from 6.2 to 227 nM (36.6-fold). At this activated state, by setting *k*_0_=0, all NRF2 species decrease, with a half-life of 68 min for *NRF2*_tot_, while *NRF2*_free_ disappears much more quickly. By setting *k*_6_ to even lower values, the maximal levels of both *NRF2*_free_ and *NRF2*_tot_ increase but to a limited extent and the half-life of *NRF2*_tot_ lengthens to 126 min in the extreme case when *k*_6_=0. (Figs. S5A and S5C).

Therefore, at the basal condition, *KEAP1_NRF2*_open1_ is the dominant form of the open KEAP1-NRF2 complex. However, upon perturbation, either by setting *k*_0_=0 or setting *k*_6_ to a lower value than the default, the relative abundance between the two open states is switched, such that *KEAP1_NRF2*_open2_ becomes dominant, and then the open:closed ratio will be following the *KEAP1_NRF2*_open2_:*KEAP1_NRF2*_closed_ ratio, which is determined largely by *k*_3_ and *k*_4_ as an equilibrium step. In this sense, although Model 3a behaves as shown above in a cycle mode globally due to the slow *k*_1.1_/*k*_2.1_ steps, locally, some species of the KEAP1-NRF2 complexes still maintain an equilibrium relationship, due to the high fluxes of the *k*_1_/*k*_2_ and *k*_3_/*k*_4_ binding steps.

With increasing *CLASS*_I-V_ levels, the temporal behaviors of *NRF2*_tot_ (Fig. 2D), *NRF2*_free_ (Fig. 2E), and *KEAP1*_free_ (Fig. 2F) are similar to Model 2. It takes a longer time for *NRF2*_tot_ to reach steady states, while the *NRF2*_free_ response is initially delayed but its rising time is shortened with increasing *CLASS*_I-V_ levels. For steady-state dose-response relationships, *KEAP1_NRF2*_open2_tot_ (*KEAP1_NRF2*_open2_ + *KEAP1*_o_*_NRF2*_open*2*_) and *KEAP1_NRF2*_closed_tot_ (*KEAP1_NRF2*_closed_ + *KEAP1*_o_*_NRF2*_closed_) both increase while remaining at a constant equilibrium ratio with increasing *CLASS*_I-V_ levels (Fig. 2G). In contrast, steady-state *KEAP1_NRF2*_open1_tot_ (*KEAP1_NRF2*_open1_ + *KEAP1*_o_*_NRF2*_open1_) first increases slightly then decreases (Fig. 2G). Steady-state *NRF2*_free_ exhibits an ultrasensitive, sigmoidal dose-response with respect to *CLASS*_I-V_ levels, with *n*_H_ of 1.78 and *LRC*_max_ of 2.24 (Fig. 2G).

Flux analysis shows that the *total degradation rate* curve exhibits an S-shape as in Model 2 (Figs. 2H and 2I). However, because of the two-step ETGE binding, higher concentrations of *NRF2*_free_ are required to produce levels of turnover fluxes similar to Model 2, with the *flux*_k6_ and *flux*_k9_ curves shifted to the right and closer to *flux*_k5_. This shift leads to a shorter second phase of the *total degradation rate* curve that is not as flat as in Model 2. As the *k*_6_ value is varied mimicking different stress levels, the intersection point between *synthesis rate* and *total degradation rate* curves still swings quite dramatically, albeit not as dramatic as in Model 2 (Figs. S3G and S3H). As shown in Figs. 2H and 2I, when *k*_6_ is lowered from 5.325E-4 to 1.775E-4, a 3-fold decrease, the corresponding steady-state *NRF2*_free_ concentration increases by 13-fold, indicating signal amplification.

#### *Effects of k*_1_ *(k’*_1_) and *k*_2_ *(k’*_2_)

We next examined the effects of different parameters on the NRF2 response in Model 3a. Enhancing the ETGE-mediated first-step binding affinity between free KEAP1 and free NRF2, by increasing *k*_1_ and *k’*_1_ by 10-fold, only marginally decreases the basal *NRF2*_tot_ level and half-life (Fig. S6A) with nearly no effect on the steady-state dose-response curve (Fig. 3A). Neither the basal levels of different open and closed KEAP1-NRF2 complexes nor their steady-state dose-response curves are affected (Figs. S6E-S6H). In contrast, the basal *NRF2*_free_ level decreases dramatically and the ultrasensitivity of the dose-response curve is enhanced markedly without much change in the maximal level (Fig. 3B). Decreasing *k*_1_ and *k’*_1_ by 10-fold appears to have slightly larger albeit opposite effects on the various NRF2 species (Figs. 3A and S6), and dramatically increases the *NRF2*_free_ level and reduces its ultrasensitivity (Fig. 3B). The time delay in the *NRF2*_free_ response disappears with decreasing *k*_1_ and *k’*_1_ (Fig. S6C) and is further increased with increasing *k*_1_ and *k’*_1_ (Fig. S6D). Varying *k*_2_ and *k’*_2_ has opposite effects as varying *k*_1_ and *k’*_1_ (simulation results not shown).

**Figure 3.**
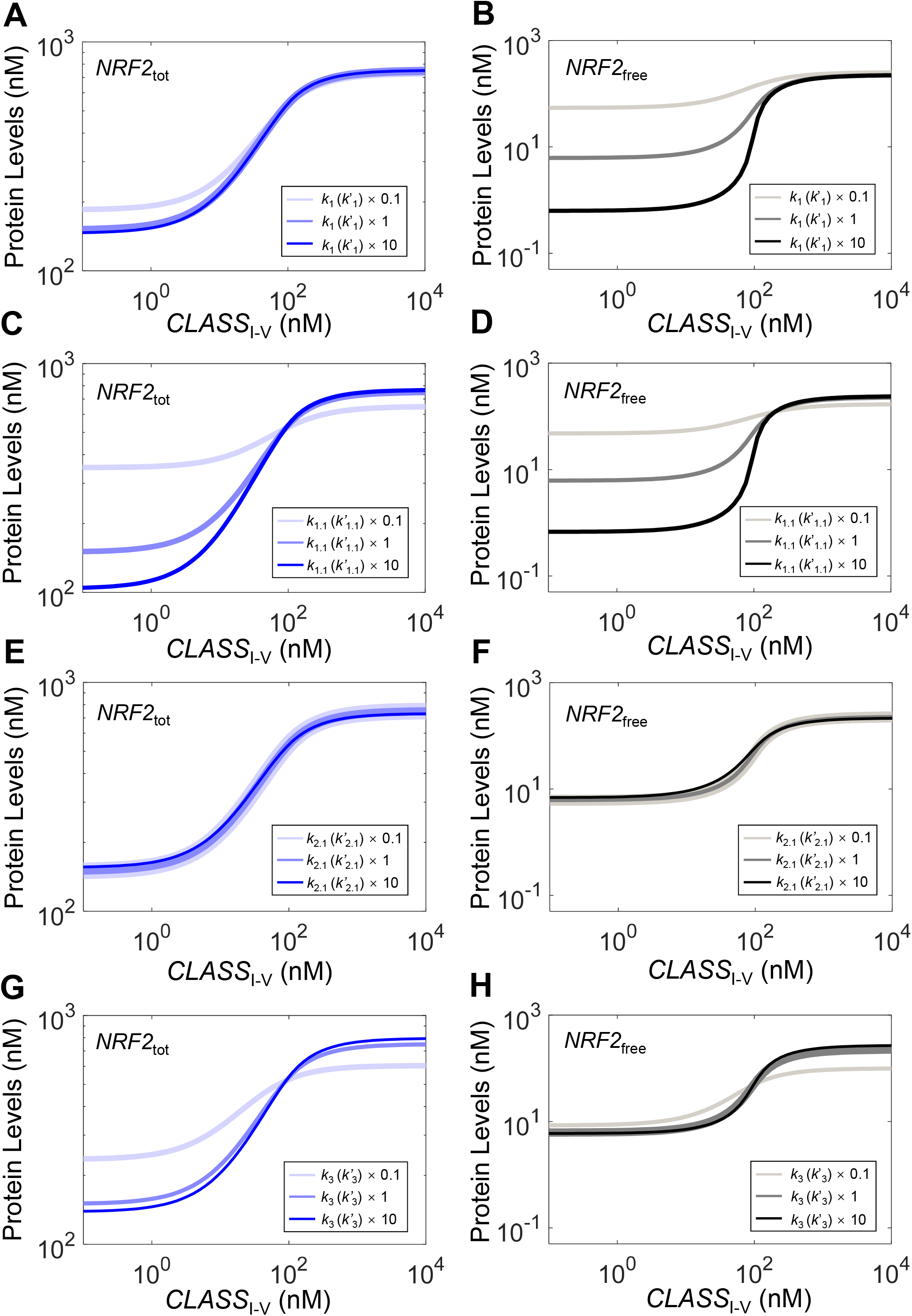
Effects of KEAP1-NRF2 binding parameters on NRF2 responses in Model 3a. Effects of varying **(A-B)** *k*_1_ (*k’*_1_), **(C-D)** *k*_1.1_ (*k’*_1.1_), **(E-F)** *k*_2.1_ (*k’*_2.1_), and **(G-H)** *k*_3_ (*k’*_3_) on steady-state dose-response curves of *NRF2*_tot_ (left panels) and *NRF2*_free_ (right panels). Note *k*_i_=*k’*_i_ for i = 1, 1.1, 2.1, or 3; x0.1, x1, and x10 denote 0.1, 1, and 10 times default values.

#### *Effects of k*_1.1_ *(k’*_1.1_) and *k*_2.1_ (*k’*_2.1_)

We next examined the effects of the ETGE-mediated second-step binding, which is much slower than the first step and is the key step for making Model 3a behave in a cycle mode. Increasing *k*_1.1_ and *k’*_1.1_ shifts the balance between the two open states, causing a reduction in *KEAP1_NRF2*_open1_tot_ (Fig. S7E) but only a slight increase in *KEAP1_NRF2*_open2_tot_ (Fig. S7F) and *KEAP1_NRF2*_closed_tot_ (Fig. S7H), resulting in a net decrease of the total open state *KEAP1_NRF2*_open_tot_ (Fig. S7G). As a result, the basal level of *NRF2*_tot_ is reduced with a slight decrease in its half-life (Fig. S7A) and the steady-state dose-response curve becomes steeper (Fig. 3C). In comparison, the basal level of *NRF2*_free_ is considerably reduced and the ultrasensitivity of the dose-response curve is dramatically enhanced with little change in the maximal response level (Fig. 3D). Decreasing *k*_1.1_ and *k’*_1.1_ has the opposite but generally larger effects. It causes an increase in *KEAP1_NRF2*_open1_tot_ (Fig. S7E) and a decrease in *KEAP1_NRF2*_open2_ (Fig. S7F) and *KEAP1_NRF2*_closed_ (Fig. S7H), resulting in a net increase of *KEAP1_NRF2*_open_tot_ which seems to have a flat response to *CLASS*_I-V_ (Fig. S7G). As *k*_1.1_ and *k’*_1.1_ are decreased by 10-fold, the basal level of *NRF2*_tot_ is dramatically increased with its half-life lengthened (Fig. S7A) and the steady-state dose-response curve becomes much shallower (Fig. 3C). The basal level of *NRF2*_free_ is considerably elevated and the ultrasensitivity of its dose-response curve is dramatically reduced (Fig. 3D). The time delay in the *NRF2*_free_ response disappears with decreasing *k*_1.1_ and *k’*_1.1_ (Fig. S7C) and is further increased with increasing *k*_1.1_ and *k’*_1.1_ (Fig. S7D). Varying *k*_2.1_ and *k’*_2.1_, especially when lowering the values, seems to affect *KEAP1_NRF2*_open1_tot_ the most, with a minimal effect on all other NRF2 species (Figs. 3E-3F and S7I-S7P), which is consistent with the low backward flux nature of this second-step ETGE binding, where the backward flux (*flux*_k2.1_ + *flux*_k’2.1_) is only a tiny fraction of the forward flux (*flux*_k1.1_ + *flux*_k’1.1_).

#### *Effects of k*_3_ *(k’*_3_) and *k*_4_ *(k’*_4_)

We next examined the effects of DLG-mediated binding. Increasing *k*_3_ and *k’*_3_ by 10-fold reduces the *KEAP1_NRF2*_open2_tot_ level dramatically across the range of *CLASS*_I-V_ levels as expected (Fig. S8F). However, it only marginally decreases the basal *KEAP1_NRF2*_open1_tot_ (Fig. S8E) and increases the basal *KEAP1_NRF2*_closed_tot_ (Fig. S8H) levels. At high *CLASS*_I-V_ levels, *KEAP1_NRF2*_open1_tot_ is suppressed considerably and *KEAP1_NRF2*_closed_tot_ increases to higher levels. These changes have slight effects on the basal *NRF2*_tot_ level and its half-life (Fig. S8A), and the steady-state dose-response curve (Fig. 3G). The basal *NRF2*_free_ level decreases marginally and the ultrasensitivity of the steady-state dose-response curve barely increases with a slightly higher maximal level (Fig. 3H). Decreasing *k*_3_ and *k’*_3_ by 10-fold has opposite but larger effects on the various species. With *KEAP1_NRF2*_open2_tot_ at higher levels (Fig. S8F), *KEAP1_NRF2*_open1_tot_ (Fig. S8E) becomes higher and *KEAP1_NRF2*_closed_tot_ (Fig. S8H) becomes lower. Both basal *NRF2*_tot_ and *NRF2*_free_ levels increase and maximal response levels decrease, reducing their ultrasensitivity (Figs. 3G and 3H). The time delay in the *NRF2*_free_ response does not appear to be affected by *k*_3_ and *k’*_3_ (Figs. S8B-S8D). Varying *k*_4_ and *k’*_4_ has opposite effects as varying *k*_3_ and *k’*_3_, and reducing *k*_4_ and *k’*_4_ to zero thus making the DGL-mediated binding irreversible has a similar effect as reducing *k*_4_ and *k’*_4_ by 10-fold (simulation results not shown).

#### Effects of hinge-latch mode of operation

The hinge-latch hypothesis states that under oxidative stress by class I-V oxidants, the DLG-mediated binding is weakened, likely due to the cysteine modification on KEAP1 in multiple domains, and the level of the closed KEAP1-NRF2 complex is reduced so that NRF2 is no longer destabilized (Tong et al. 2006, Tong et al. 2007, Fukutomi et al. 2014). Here we used Model 3a to explore the effects of the hinge-latch hypothesis. When setting *k’*_3_ (which is the association rate constant for the intramolecular binding between oxidized KEAP1 and DLG motif) to a lower value (0.1 of default) to mimic a hinge-latch mode of operation, a high *CLASS*_I-V_ level lead to increases in both the open and closed states (Figs. 4A and 4B). However, the open state level is higher than the closed state, which runs counter to the decreasing open:closed ratio under oxidative stress as expected (Baird et al. 2013). The hinge-latch simulation also predicts more muted maximal responses of *NRF2*_tot_ (Fig. 4E) and *NRF2*_free_ (Fig. 4F). Interestingly, increasing *k’*_3_ to simulate strengthened DLG binding under oxidative stress has the opposite effect: the open:closed ratio further increases (Figs. 4C and 4D) and the *NRF2*_tot_ (Fig. 4E) and *NRF2*_free_ (Fig. 4F) dose-responses exhibit higher maximal levels and enhanced ultrasensitivity, although these changes approach a limit as *k’*_3_ is increased by > 10-fold. Changing the DLG binding affinity by varying *k’*_4_ has opposite effects as varying *k’*_3_ (simulation results not shown). Therefore, with current parameter settings, , the hinge-latch mode of operation is predicted to be less effective in activating NRF2 by class I-V compounds.

**Figure 4.**
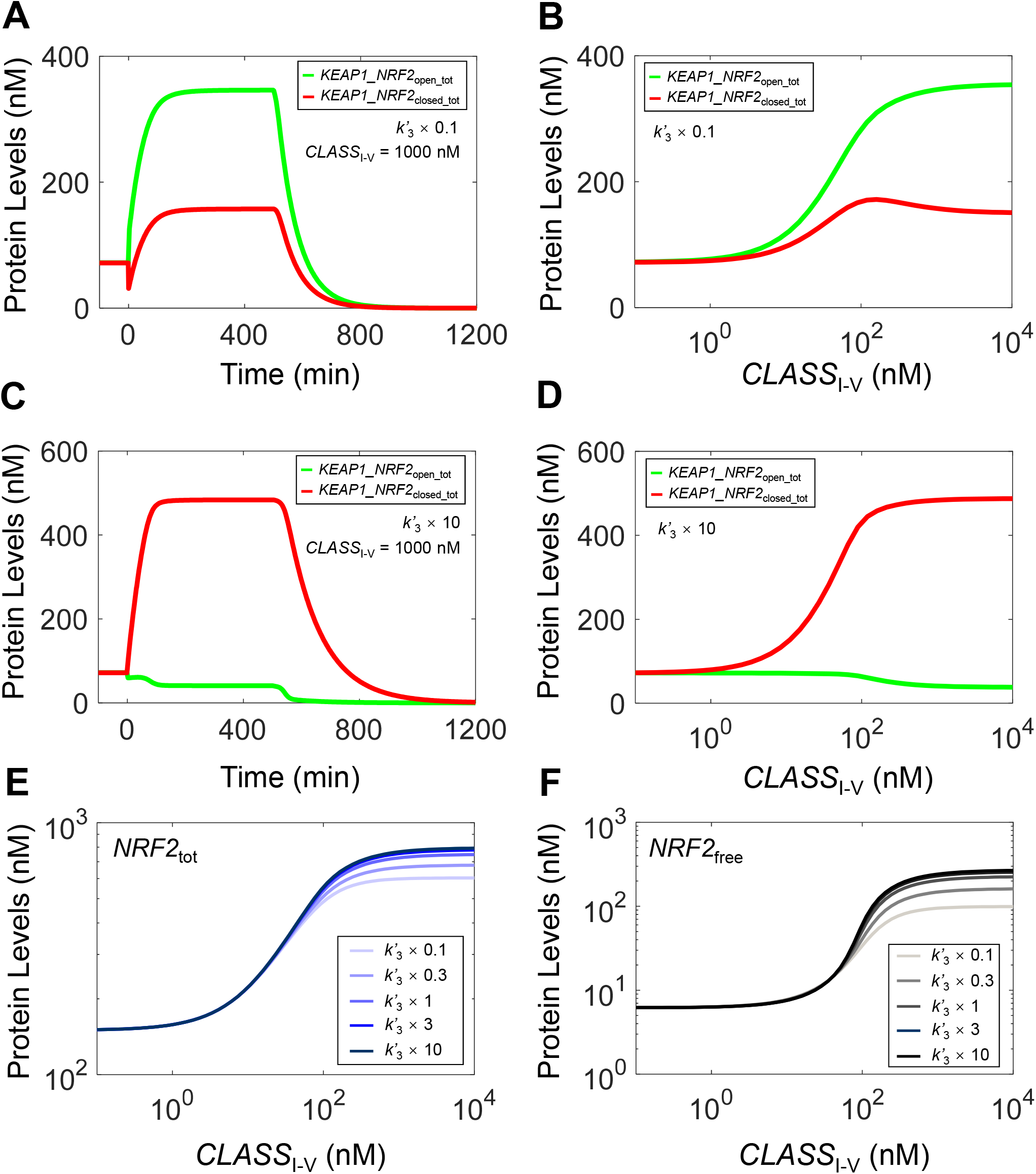
Effects of varying parameter *k’*_3_ alone on NRF2 responses in Model 3a to test hinge-latch hypothesis - with *k’*_6_ at default value. Dynamical changes of *KEAP1_NRF2*_open_tot_ and *KEAP1_NRF2*_closed_tot_ in response to a high level of *CLASS*_I-V_ at 1000 nM starting at 0 min and in response to termination of NRF2 synthesis (by setting *k*_0_=0) starting at 500 min, when *k’*_3_ is **(A)** 0.1 and **(C)** 10 times of default value. Steady-state dose-response curves of *KEAP1_NRF2*_open_tot_ and *KEAP1_NRF2*_closed_tot_ when *k’*_3_ is **(B)** 0.1 and **(D)** 10 times of default value. **(E-F)** Effects of varying *k’*_3_ on steady-state dose-response curves of *NRF2*_tot_ and *NRF2*_free_ respectively.

#### Effects of KEAP1 abundance

The relative abundance of KEAP1 and NRF2 can have important effects on NRF2 activation. The current default basal *NRF2*_tot_:*KEAP1*_tot_ ratio is about 1:4. Increasing *KEAP1*_tot_ by up to 10-fold has little effect on the basal *NRF2*_tot_ level and its half-life (Fig. S10A). This lack of effect is because at the default *KEAP1*_tot_ level, there is already sufficient KEAP1 to sequester the majority of NRF2, so increasing *KEAP1*_tot_ further does not alter the fraction of NRF2 in complex with KEAP1 much, including the closed state which is most actively degraded. But the maximal level of the dose-response curve of *NRF2*_tot_ increases (Fig. 5A) and this occurs because NRF2 in *KEAP1*_o_*_NRF2*_closed_ is not degraded as readily as *NRF2*_free_ or NRF2 in the open state. Increasing total KEAP1 abundance reduces basal *NRF2*_free_ and the maximal response levels dramatically (Fig. 5B). The muted response is mostly due to the increased sequestering effect of higher KEAP1 abundance. When *KEAP1*_tot_ is reduced from its default value, basal *NRF2*_tot_ levels and its half-life increase (Fig. S10A), and the dose-response curve becomes shallower with lower maximal response levels (Fig. 5A). Basal *NRF2*_free_ increases dramatically with little further increase in response to *CLASS*_I-V_ at higher levels, indicating constitutive activation of NRF2 (Fig. 5B). These results suggest that there is an optimal NRF2:KEAP1 ratio that can maximize the dynamic range of free NRF2 in response to oxidative stress. If the ratio is too low or too high, the response of free NRF2 is muted.

**Figure 5.**
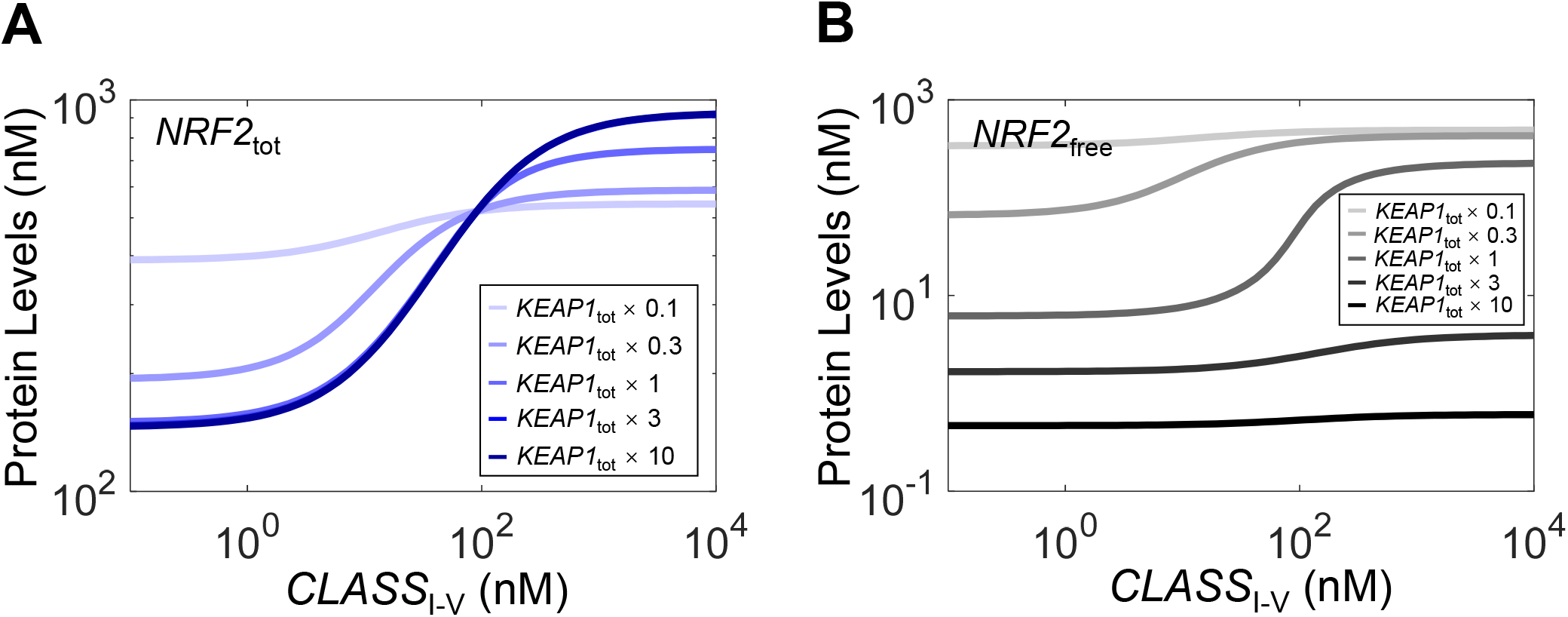
Effects of total KEAP1 abundance on NRF2 responses in Model 3a. Steady-state dose-response curves of **(A)** *NRF2*_tot_ and **(B)** *NRF2*_free_ under different values of total KEAP1 abundance relative to default value.

### Model 3b (Two-Step ETGE-Binding Cycle Mode for Class VI inducer)

Since Model 3a represents the most updated biology of KEAP1 and NRF2 interactions, the remaining Models (3b, 4a and 4b) are based on this model. In Model 3b, we simulated class VI NRF2 activators, which activate NRF2 by competing with NRF2 for binding to the DC domain of KEAP1 (Hancock et al. 2012, Jiang et al. 2014, Lazzara et al. 2020). Model 3b keeps the interactions between KEAP1 and NRF2 at the basal condition as in Model 3a, but differ in how *CLASS*_VI_ interacts with KEAP1 (Fig. 6A). We assume that a *CLASS*_VI_ molecule can bind equally to either of the two monomeric subunits in KEAP1 dimer that is not occupied by NRF2. It is thus possible that a KEAP1 dimer can be occupied by 2 molecules of a *CLASS*_VI_ compound such that no NRF2 is able to bind to this KEAP1 dimer. This assumption is well justified as It has been recently demonstrated that NRF2 can be progressively and ultimately completely liberated off KEAP1 by increasing concentrations of p62 and other KEAP1-NRF2 interaction inhibitors (Horie et al. 2021). Unlike the case with *CLASS*_I-v_ activators, in response to *CLASS*_VI_, *NRF2*_free_ increases immediately without delay, followed by a slower rise over time to reach the steady state in about 300 min (Fig. 6C). The initial rapid response of *NRF2*_free_ is due to the titration of KEAP1 by *CLASS*_VI_, resulting in immediate liberation of NRF2 from the KEAP1-NRF2 complexes. The subsequent slow *NRF2*_free_ rise occurs because more KEAP1-NRF2 complex shifts away from the rapidly-degrading closed state, resulting in NRF2 stabilization (Fig. 6E). Contrary to Model 3a for *CLASS*_I-v_, the higher the *CLASS*_VI_ level, the long it takes for *NRF2*_free_ to reach the steady state (Fig. 6C). *NRF2*_tot_ has a similar temporal profile to *NRF2*_free_ except the initial fast-rising phase (Fig. 6B). The steady-state dose responses of *NRF2*_free_ and *NRF2*_tot_ are shown in Fig. 6D, with *NRF2*_free_ exhibiting an *n*_H_ of 1.09 and *LRC*_max_ of 0.92. The *NRF2*_free_ and *NRF2*_tot_ responses to low *CLASS*_VI_ levels are nearly flat, as *CLASS*_VI_ molecules are first sequestered by free KEAP1. Contrary to the decreasing open:closed ratio of KEAP1-NRF2 complexes under *CLASS*_I-v_, this ratio increases by *CLASS*_VI_ (Fig. 6E). At high *CLASS*_VI_ levels, the half-life of *NRF2*_tot_ approaches 40 min, which is also the half-lives of *NRF2*_free_ and *KEAP1_NRF2*_open_tot_ (Fig. S11). We also explored the situation when only one KEAP1 monomeric subunit can be occupied by class VI activators by setting both *k’*_7_ and *k’*_8_ to zero. As shown in Figs. 6F and 6G, this configuration does not affect *NRF2*_tot_, but weakens the *NRF2*_free_ response as its maximal level cannot reach as high as when both KEAP1 monomeric subunits can be occupied by class VI activators. This more muted response is because without class VI activators blocking both binding sites on KEAP1 dimer, NRF2 can still be sequestered by KEAP1 through the ETGE motif, resulting in lower *NRF2*_free_ levels.

**Figure 6.**
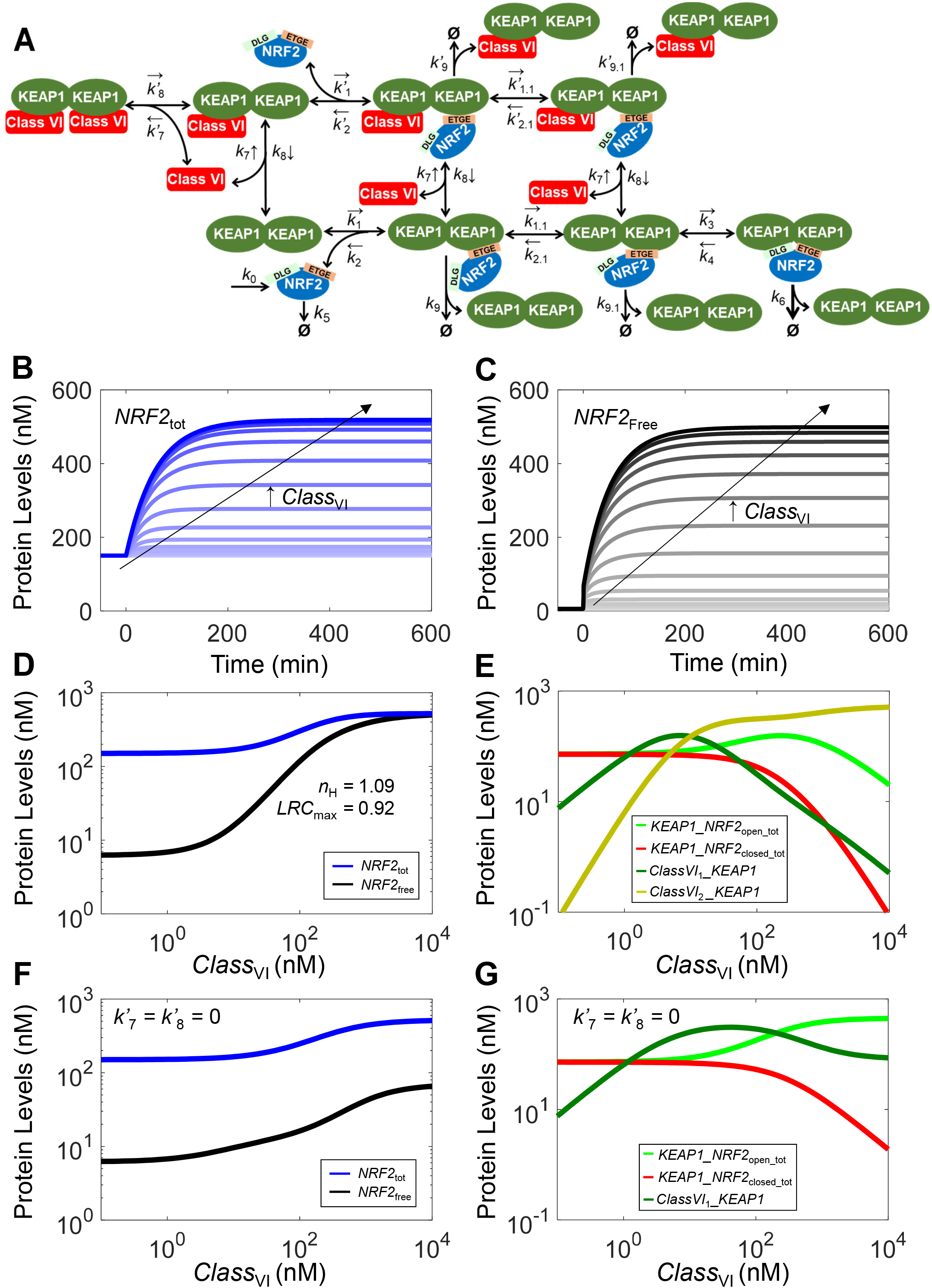
Structure, and dynamical and steady-state behaviors of Model 3b. **(A)** Structure of Model 3b featuring two-step ETGE binding and interaction with class VI activator. Dynamical changes of **(B)** *NRF2*_tot_ and **(C)** *NRF2*_free_ in response different levels of *CLASS*_VI_. **(D)** Steady-state dose-response curves of *NRF2*_tot_ and *NRF2*_free_. Shown are *n*_H_ and *LRC*_max_ for *NRF2*_free_; *n*_H_ and *LRC*_max_ for *NRF2*_total_ are 1.35 and 0.35 respectively (not shown). **(E)** Steady-state dose-response curves of *KEAP1_NRF2*_open_tot_*, KEAP1_NRF2*_closed_tot_, *ClassVI*_1__*KEAP1* (Class VI activator-KEAP1 complex containing one activator molecule) and *ClassVI*_2__*KEAP1* (containing two activator molecules). **(F-G)** Steady-state oxidant-response curves of *NRF2*_tot_, *NRF2*_free_, *KEAP1_NRF2*_open_tot_*, KEAP1_NRF2*_closed_tot_, and *ClassVI*_1__*KEAP1* under condition when only one class VI activator molecule is allowed to bind to KEAP1 by setting *k’*_7_=*k’*_8_=0.

### Model 4a (With Nucleus for Class I-V activators)

Since NRF2 that translocates to the nucleus is what ultimately drives target gene expression, we next explored the situation when a nuclear compartment is added. The following assumptions were made regarding NRF2 translocation between the cytosol and nucleus (Fig. 7A). (i) The binding kinetics between free nuclear NRF2 (*NRF2*_free_nucleus_) and free nuclear KEAP1 dimer (*KEAP1*_free_nucleus_) are the same as in the cytosol. (ii) KEAP1-mediated NNRF2 ubiquitination and degradation does not occur in the nucleus. Therefore, the degradation rate constants of various NRF2 species in the nucleus are the same as in the cytosol, except for the closed KEAP1-NRF2 complex, which is degraded with the same rate constant as other NRF2 species. (iIi) NRF2 activators do not modify or bind to KEAP1 in the nucleus to regulate NRF2 stability.

**Figure 7.**
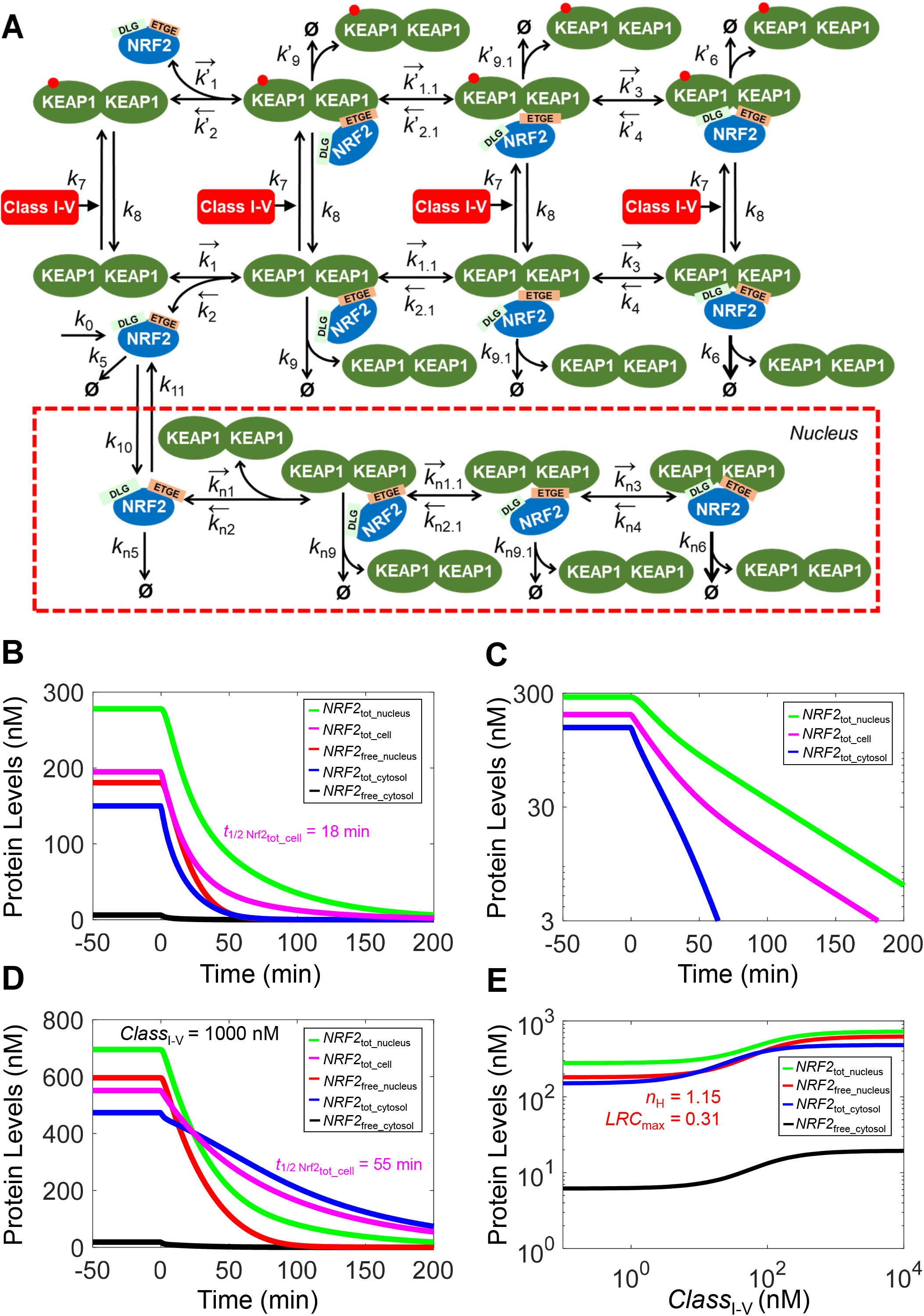
Structure, and dynamical and steady-state behaviors of Model 4a. **(A)** Structure of Model 4a featuring two-step ETGE binding, nuclear NRF2 translocation, and interaction with class I-V activator. **(B)** Dynamical changes of basal *NRF2*_tot_cell_, *NRF2*_tot_nucleus_, *NRF2*_tot_cytosol_, *NRF2*_free_nucleus_, and *NRF2*_free_cytosol_ in response to termination of NRF2 synthesis (by setting *k*_0_=0) starting at 0 min. **(C)** *NRF2*_tot_cell_, *NRF2*_tot_nucleus_, and *NRF2*_tot_cytosol_ in (B) shown in log Y scale. **(D)** Dynamical changes of various NRF2 species previously induced by a high level of *CLASS*_I-V_ at 1000 nM in response to termination of NRF2 synthesis (by setting *k*_0_=0) starting at 0 min. **E)** Steady-state dose-response curves of various NRF2 species. *n*_H_ and *LRC*_max_ of *NRF2*_free_nucleus_ curve are indicated.

At the basal condition, total nuclear NRF2 (*NRF2*_tot_nucleus_) is at 278 nM as observed in RAW 264.7 cells (Iso et al. 2016) and a significant fraction of which is titrated by KEAP1 such that *NRF2*_free_nucleus_ is at 180 nM (Table S8). When setting *k*_0_=0, *NRF2*_tot_cytosol_, *NRF2*_tot_nucleus_, and *NRF2*_tot_cell_ all decay but at different paces, with corresponding half-lives of about 11, 28, 18 min, respectively (Fig. 7B). When viewed on log scale, it is apparent that *NRF2*_tot_cell_ decays in two phases, a fast phase followed by a slow one (Fig. 7C). This two-phase decay profile of total cellular NRF2 is caused by the fast cytosolic and slow nuclear decay and has been observed experimentally in a variety of cell lines (Khalil et al. 2015). Under high stress when *CLASS*_I-v_ = 1000 nM, the half-life of *NRF2*_tot_cell_ markedly lengthens to 55 min (Fig. 7D). In response to a range of *CLASS*_I-v_ levels, free and total NRF2 in both cytosol and nucleus rise and reach steady states in about 300 min (Fig. S12). In contrast to Model 3a which does not have the nucleus compartment, *NRF2*_free_cytosol_ rises to much lower levels (Fig. S12A) as most of it translocates into the nucleus elevating *NRF2*_free_nucleus_ (Fig. S12C). The steady-state dose-response relationship for *NRF2*_free_nucleus_ exhibits a shallow response, with *n*_H_ of 1.15 and of *LRC*_max_ of 0.31 (Fig. 7E). The maximal response levels of *NRF2*_tot_nucleus_ and *NRF2*_free_nucleus_ increase by 2.6 and 3.5-fold respectively, while those of *NRF2*_tot_cytosol_ and *NRF2*_free_cytosol_ both increase by about 3.2 and 3.1-fold, respectively (Tables S8 and S9). Thus, with a nuclear load, NRF2 activation is not as robust as when the action is limited to the cytosol only. This lessor response contrasts with the nearly 10-fold increase in nuclear NRF2 under exposure to DEM at 100 µM for 3 h observed in RAW 264.7 cells which our model is partially based upon (Iso et al. 2016).

The overall muted response of Model 4a is due to the following reasons. At the basal condition, the net influx of NRF2 from the cytosol to nucleus is *flux*_k10_ - *flux*_k11_ = 0.0434 nM/s, which is about 22% of *k*_0_ (0.1933), the NRF2 synthesis rate in the cytosol. Therefore, even if a *CLASS*_I-v_ activator can divert all synthesized NRF2 into the nucleus, the total nuclear NRF2 can only increase by a maximal 4.45-fold (0.193/0.0434) with a constant nuclear NRF2 half-life. We wondered if the relative abundance of nuclear KEAP1 and NRF2 plays a role in determining the magnitude of the nuclear NRF2 response. When *KEAP1*_tot_nucleus_ abundance is increased (with *k*_10_ adjusted simultaneously to maintain the same basal *NRF2*_tot_cytosol_ and *NRF2*_tot_nucleus_ concentrations), the simulations showed that both the basal and maximally-induced levels of *NRF2*_free_nucleus_ decease because of the sequestering effect of KEAP1 (Fig. S13C). However, the degree of ultrasensitivity of the *NRF2*_free_nucleus_ dose-response curve seems to be optimal when *KEAP1*_tot_nucleus_ is at an intermediate abundance. Increasing *KEAP1*_tot_nucleus_ also leads to changes in the maximal levels of *NRF2*_tot_cytosol_, *NRF2*_free_cytosol_, and *NRF2*_tot_nucleus_ (Fig. S13), but the fold-increase of *NRF2*_tot_nucleus_ remains relatively low. These results suggest that other mechanisms, as described in the Discussion, may operate *in vivo* to produce a more robust nuclear NRF2 response. One possibility is a smaller nuclear NRF2 load at the basal condition. As shown in Fig. S14, when reducing the basal *NRF2*_tot_nucleus_ level and thus the nuclear load through simultaneously adjusting *k*_0_ and *k*_10_ while maintaining the same basal *NRF2*_tot_cytosol_ level, the magnitude of both *NRF2*_tot_nucleus_ and *NRF2*_free_nucleus_ responses improve considerably.

### Model 4b (With Nucleus for Class VI activators)

We next considered the situation of a class VI activator which competes with NRF2 for binding to KEAP1 in Model 4b (Fig. 8A). The model assumptions are similar to Model 4a and the class VI activator only operates in the cytosol. Model 4b has an interesting dynamic. In response to a range of *CLASS*_VI_ levels, there is a quick spike in *NRF2*_free_cytosol_ within a couple of minutes followed by a slow rise (Fig. 8B). Correspondingly, *NRF2*_tot_cytosol_ decreases immediately followed by a slower increase before setting to steady states (Fig. 8C). The rapid increase in *NRF2*_free_cytosol_ results from the immediate liberation of NRF2 from the KEAP1-NRF2 complex, and the liberated NRF2 moves quickly into the nucleus, causing *NRF2*_free_nucleus_ (Fig. 8D) and *NRF2*_tot_nucleus_ to rise quickly followed by a slower increase to steady states. Under high stress when *CLASS*_VI_ = 1000 nM, the half-life of *NRF2*_tot_cell_ lengthens to 40 min (Fig. 8E), shorter than that in Model 4a. However, the steady-state *NRF2*_free_nucleus_ and *NRF2*_tot_nucleus_ levels can increase to higher levels, maximally by 6 and 4.2-fold from their basal levels, respectively (Tables S8 and S9). This is because by outcompeting NRF2 for KEAP1, *CLASS*_VI_ can drive more NRF2 into the nucleus (Fig. S12C vs. Fig. 8D). The steady-state dose-response curve *NRF2*_free_nucleus_ is shallow, with *n*_H_ of 1.09 and of *LRC*_max_ of 0.46 (Fig. 8F). Interestingly, the steady-state dose-response curve of *NRF2*_tot_cytosol_ monotonically decreases, from the basal 150 nM to 33 nM for higher levels of *CLASS*_VI_. This decrease occurs because KEAP1 dimer is gradually titrated away by *CLASS*_VI_ activator, leaving fewer NRF2 in the KEAP1-bound form (Fig. 8G), and more NRF2 translocates to the nucleus. As in Model 4a, varying nuclear KEAP1 and lowering basal nuclear load of NRF2 turnover can also improve the magnitude and ultrasensitivity of *NRF2*_free_nucleus_ (Figs. S15 and S16).

**Figure 8.**
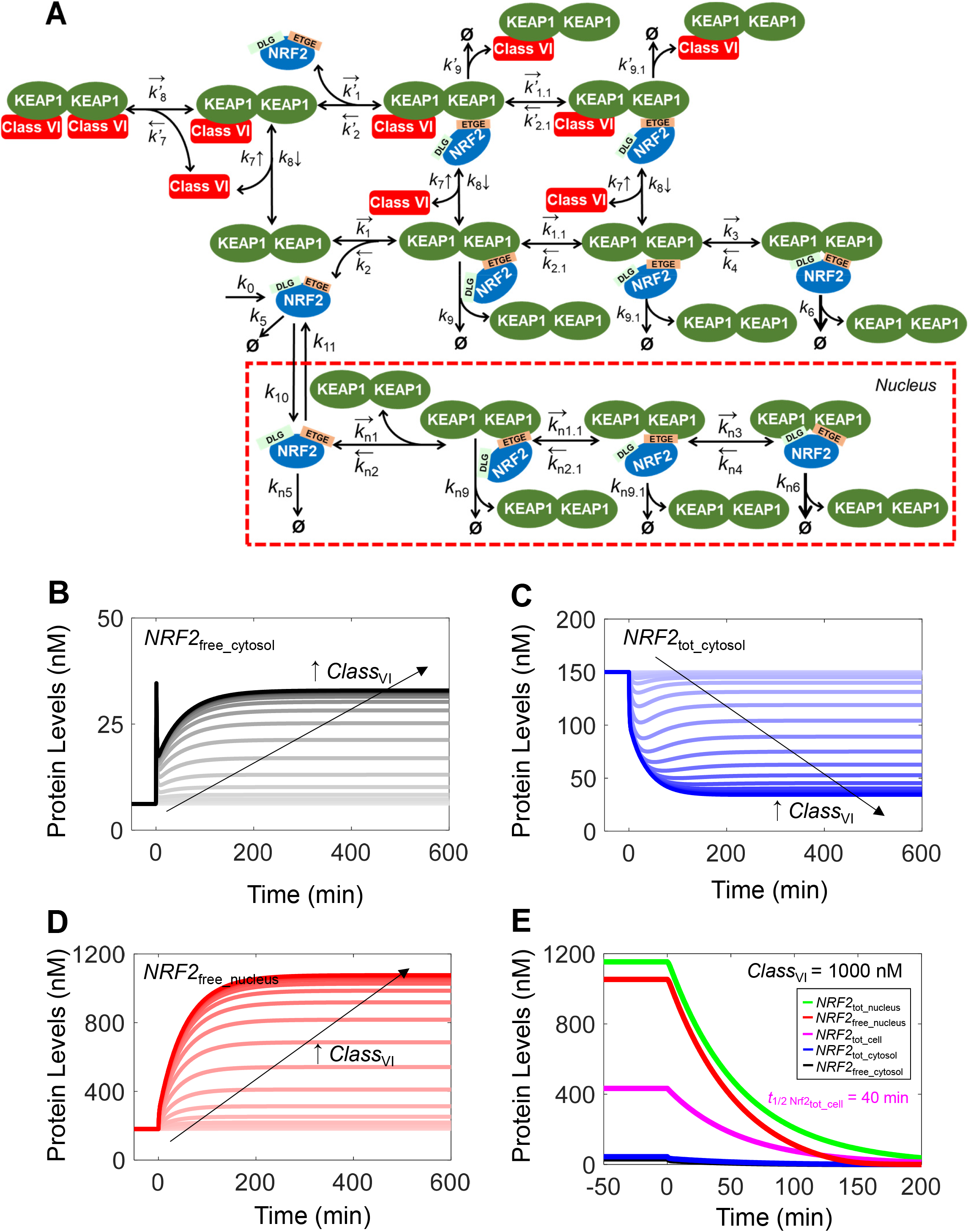
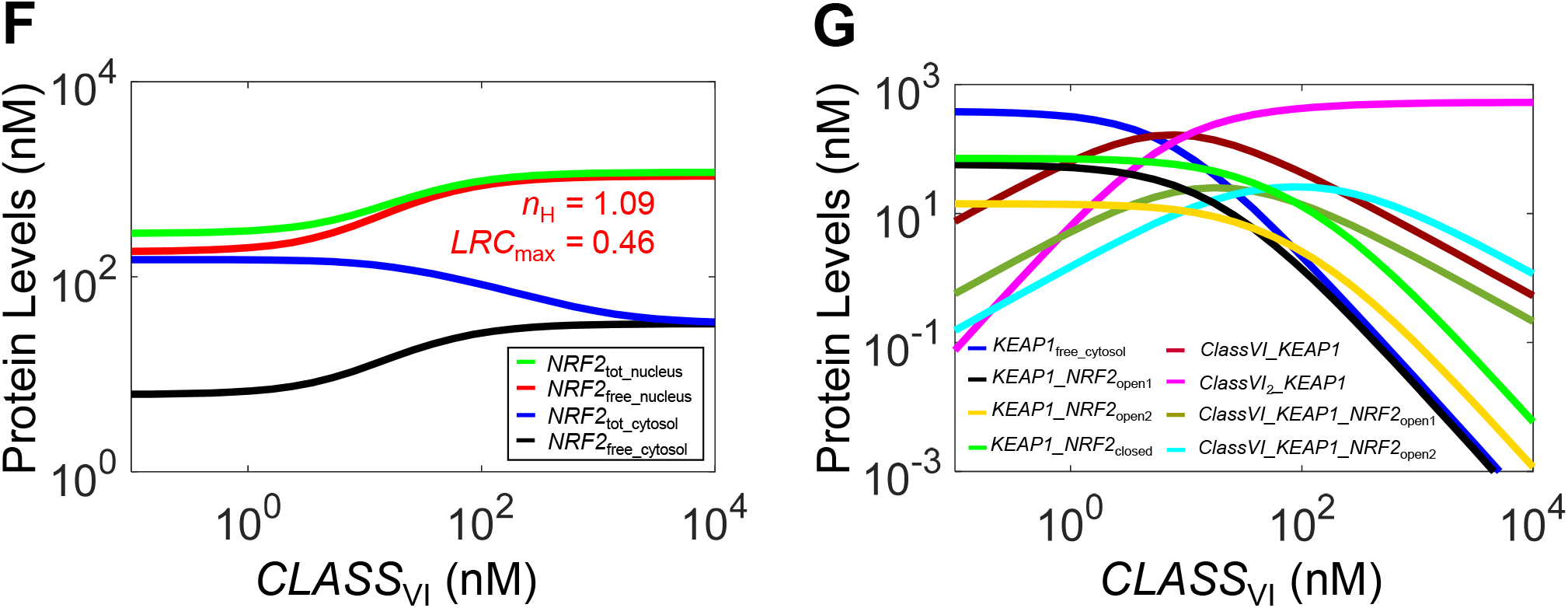
Structure, and dynamical and steady-state behaviors of Model 4b. **(A)** Structure of Model 4b featuring two-step ETGE binding, nuclear NRF2 translocation, and interaction with class VI activator. Dynamical changes of **(B)** *NRF2*_free_cytosol_, **(C)** *NRF2*_tot_cytosol_, and **(D)** *NRF2*_free_nucleus_ in response to different levels of *CLASS*_VI._ **(E)** Dynamical changes of various NRF2 species previously induced by a high level of *CLASS*_VI_ at 1000 nM in response to termination of NRF2 synthesis (by setting *k*_0_=0) starting at 0 min. **(F-G)** Steady-state dose-response curves of various NRF2 species and KEAP1 species respectively.

## DISCUSSION

NRF2 activation is an essential step toward the transcriptional induction of adaptive antioxidant responses. It is mediated via a unique mechanism of protein stabilization where KEAP1 functions as both a redox sensor and regulator. While the molecular interactions involved in this process have been well characterized qualitatively and to some extent quantitatively, the quantitative systems-level behaviors of this redox transducer module are still poorly understood. In the present study, we explored the steady-state and dynamic behaviors of the KEAP1-NRF2 interactions through a series of mathematical models of increasing complexity. Our simulations demonstrated that the kinetic details of the molecular interactions between KEAP1 and NRF2 play critical roles in determining the redox signaling properties.

### Basal NRF2 half-life in relation to different NRF2 states

A prominent function of KEAP1 is to act as an E3 ligase adaptor to promote NRF2 ubiquitination and degradation. This function is critically dependent on the configuration of the KEAP1-NRF2 complex. It is well-established that for the ubiquitination and degradation of NRF2 to occur, the KEAP1-NRF2 complex has to be in the closed state, i.e., both of the two binding sites in the cysteine-unmodified KEAP1 dimer have to be engaged by the ETGE and DLG motifs of the same NRF2 molecule. Therefore, the fraction of this closed state and the rate at which NRF2 within this closed KEAP1-NRF2 complex is ubiquitinated and degraded are key determinants for the half-life of NRF2 in the cytosol.

In the confine of the present model structure, NRF2 exists in three forms: free, open KEAP1-NRF2 complex, and closed KEAP1-NRF2 complex. The relative abundances of these forms at the basal steady state are determined by the binding parameters as well as the degradation rate constant of each NRF2 form. Given the high binding affinity between KEAP1 and ETGE, it is expected in theory and shown by our simulation that the fraction of free cytosolic NRF2 is very small, when KEAP1 is not limiting, and NRF2 exists predominantly in the complex forms at the basal condition. Using FRET to track the open and closed states of the KEAP1-NRF2 complex, Baird 2013 showed that at least in HEK293 cells, the open:closed ratio of the KEAP1-NRF2 complex is near 1:1 under nonstressed conditions (Baird et al. 2013). The half-life of total NRF2 in whole cells at basal conditions is short, mostly ranging between 6-20 min depending on cell types (Kwak et al. 2002, Alam et al. 2003, Itoh et al. 2003, Stewart et al. 2003, Kobayashi et al. 2004, He et al. 2006, Khalil et al. 2015, Crinelli et al. 2021). Since nuclear NRF2 is relatively more stable than cytosolic NRF2 (Itoh et al. 2003, Burroughs et al. 2018) and often constitutes a considerable fraction of total NRF2 at the basal condition (Khalil et al. 2015, Iso et al. 2016), it is expected that cytosolic NRF2 is actually degraded at even faster rates than measured in whole cells. The comparable basal abundance of open and closed KEAP1-NRF2 complexes suggests that NRF2 in the closed form has to be degraded very fast with a half-life of its own that is much shorter than the averaged half-life of total NRF2. In our models, parameter *k*_6_ governs the degradation of this NRF2 form. With an apparent half-life of cytosolic total NRF2 around 10 min at the basal condition, the default values of *k*_6_ across the six models correspond to an half-life of 5.7-6.6 min. In comparison, the half-lives of free NRF2 and NRF2 in the open KEAP1-NRF2 complex, as determined by parameters *k*_5_ and *k*_9_ (and *k*_9.1_ in the case of two-step ETGE binding) respectively, are much longer, which is 40 min here, as reported for COS-1 and HEK293T cells (McMahon et al. 2004, Rada et al. 2011). If *k*_5_, *k*_9_, and *k*_9.1_ are set lower than the current default value, *k*_6_ needs to be even higher to maintain the same basal total NRF2 half-life. Therefore, the turnover of basal NRF2 is predominantly routed through the closed KEAP1-NRF2 complex, and the apparent half-life of cytosolic total NRF2 is determined by the fraction of the closed complex. In Model 1 which operates in an equilibrium mode, this fraction remains constant at 50% at all times (Fig. S1A), therefore the instantaneous half-life of total NRF2 at any given moment is fixed. In the remaining models which operate in a cycle mode, the fraction increases dynamically and becomes dominant over other NRF2 forms during the decay process (Figs. S3A and 2B), resulting in a nonlinear degradation of total NRF2 with shortening instantaneous half-life. In Models 4a and 4b, which has the nucleus compartment, cellular total NRF2 decays with a two-phase profile, which has been observed experimentally in a variety of cell types (Khalil et al. 2015), reflecting the differential half-lives of cytosolic and nuclear NRF2.

### Equilibrium vs cycle mode of operation

The comparable abundance of the open and closed states of the KEAP1-NRF2 complex at the basal condition can be achieved in theory in two ways, depending on the transition fluxes (*flux*_k3_ and *flux*_k4_) between the two states relative to other turnover fluxes (*flux*_k5_, *flux*_k6_*, flux*_k9_ and *flux*_k9.1_). If the transition fluxes are much higher than the turnover fluxes, then the open and closed states of the KEAP1-NRF2 complex operate in an equilibrium mode, which means that the ratio of the two states is predominantly determined by the *k*_3_:*k*_4_ ratio regardless of other parameter values. Parameters *k*_3_ and *k*_4_ describe the DLG-mediated KEAP1 and NRF2 binding. In the literature, its binding kinetics was determined *in vitro* by using mouse KEAP1-DC fragment and NRF2-Neh2 domain fragment (Tong et al. 2006) or extended DLG motif peptide (DLGex) (Fukutomi et al. 2014). However, *in vivo* the DLG binding is mostly an intra-molecular event within the open KEAP1-NRF2 complex, since the ETGE motif has a much higher binding affinity for KEAP1 and ETGE-mediated binding almost always occurs first to form the open-state complex (Stewart et al. 2003). In such *in vivo* scenario, *k*_3_ is actually a first-order, as opposed to a second-order, association rate constant while *k*_4_ remains as a first-order dissociation rate constant. It is unclear whether in the open state the DLG binding is enhanced since the DLG is in a closer vicinity to the unoccupied KEAP1 binding site than the DLG in a free NRF2 molecule not yet attached to KEAP1. Regardless, in our first trial, in Model 1 we used the *k*_4_ value measured *in vitro* (Fukutomi et al. 2014) and adjusted *k*_3_, as detailed in Table S1 footnote, to achieve a 1:1 ratio for the basal open:closed states. Examining the fluxes clearly revealed that with these parameter settings, *flux*_k3_ and *flux*_k4_ are absolutely dominant over other turnover fluxes (Table S10). As shown in Fig. S1, the open and closed KEAP1-NRF2 complexes remain at a 1:1 ratio in all perturbed conditions including shutdown of NRF2 synthesis (Fig. S1A), stabilization of NRF2 in the closed state (Fig. S1B), and under a wide range of *CLASS*_I-V_ levels (Fig. S1C), demonstrating that Model 1 definitely operates in an equilibrium mode. However, the experimental study by Baird clearly demonstrated that under various perturbations similar to above, the closed KEAP1-NRF2 state will eventually dominate over the open state, thus negating an equilibrium mode of operation (Baird et al. 2013). It was further suggested that the KEAP1-NRF2 interaction may operate instead in a global cycle mode where the transition fluxes between the open and closed states are not overwhelmingly higher than the turnover fluxes, such that KEAP1 in the complex is almost always moved forward to the next state and eventually exists via the closed complex along with NRF2 degradation, and recycled to join the free KEAP1 dimer pool.

The discrepancy between our Model 1’s behavior and the cycle mode of operation led us to Model 2 where the parameters *k*_3_ and *k*_4_ were lowered to alleviate the equilibrium mode of operation. Model 2 indeed exhibits the behavior consistent with the cycle mode, where the open:closed ratio decreases in all perturbed conditions (Figs. S3A, S3B, and S3F). The issue with Model 2 is that parameter *k*_4_, which describes the dissociation rate constant of DLG binding, is 1.0E-4 S^-1^, only about 1/2000 of the *in vitro* measured value (Table S1). This value translates into an average lifetime of 167 min for the closed state before it can revert back into the open state, which is considered too long for such weak binding (Fukutomi et al. 2014). In the same study it was also demonstrated that ETGE-mediated binding is actually a two-step process, involving an initial fast binding step followed by a subsequent slow binding step. We therefore wondered whether the slow binding here may be responsible for the cycle mode behavior. When this idea was implemented in Model 3, simulations indeed showed such effects on the open:closed ratio under all perturbed conditions, including shutdown of NRF2 synthesis (Fig. 2B), stabilization of NRF2 in the closed state (Fig. 2C), and under a wide range of *CLASS*_I-V_ levels (Fig. 2G). With the parameter setting for the second-step, slow ETGE binding (*k*_1.1_ and *k*_2.1_), *flux*_k1.1_ is much greater than *flux*_k2.1_ (Table S10). *KEAP1_NRF2*_open1_ is the dominant form of the open state at the basal condition (Fig. 2B and Table S8), and is not in equilibrium with *KEAP1_NRF2*_open2_ (Fig. 2C and 2G). In contrast, *KEAP1_NRF2*_open2_ is always in equilibrium with *KEAP1_NRF2*_closed_ at an approximate 1:5 ratio as determined by the *k*_4_:*k*_3_ ratio (Fig. 2G). During *CLASS*_I-V_ perturbation, *KEAP1_NRF2*_open1_ decreases while *KEAP1_NRF2*_open2_ increases and becomes the dominant open form, resulting in an overall open:closed ratio that is close to 1:3.5. Therefore, although Model 3 behaves globally in a cycle mode at the basal condition with *KEAP1_NRF2*_open1_ as the dominant open form, at high stress levels *KEAP1_NRF2*_open2_ becomes the dominant open form and the system switches to operate largely in equilibrium mode as far as the overall open:closed ratio is concerned.

### Hinge-latch hypothesis and class I-V vs. class VI NRF2 activators

An important theory of NRF2 activation is the hinge-latch hypothesis which postulates that the ETGE-mediated association (i.e., the open-state complex) is always there functioning as a hinge between KEAP1 dimer and NRF2, while the weaker DLG-mediated association can be latched on (i.e., forming the closed-state complex) or off (i.e., reverting to the open state complex) by oxidative stressors (Yamamoto et al. 2018). With Model 3a we tested the effects of hinge-latch mode of operation on NRF2 activation by altering the *k’*_3_:*k’*_4_ ratio which governs the intramolecular DLG binding affinity between oxidized/conjugated KEAP1 and NRF2. Our simulations showed that when the DLG binding affinity is lowered to mimic a hinge-latch, the maximally induced steady-state levels of total NRF2 and particularly free NRF2 is tangibly reduced (Figs. 4E and 4F). In contrast, when the *k’*_3_:*k’*_4_ ratio is made higher, i.e., a strengthening of the latched-on state under oxidative stress, there is an increase, albeit limited, in the maximal NRF2 levels. These results suggest that a hinge-latch mode of operation may lead to a lessor NRF2 response to class I-V compounds. The reason for the more muted response in our model is because under oxidative stress, the closed KEAP1-NRF2 complex in which the KEAP1 molecule is modified on the sensor cysteine residues, i.e., *KEAP1*_o__*NRF2*_closed_, has a half-life (determined by *k’*_6_) even longer than those of NRF2 in the free or open complex forms. Therefore, KEAP1 here reverses the normal role of promoting NRF2 degradation as at the basal condition, and becomes instead protective of the NRF2 molecule. We also examined the situation when *KEAP1*_o__*NRF2*_closed_ is not protective of NRF2 by setting *k’*_6_ equal to *k*_5_ such that it degrades with the same half-life as free NRF2 and as open complex. In this case, the hinge-latch mode of operation slightly improves the ultrasensitivity of free NRF2 and total NRF2 (Figs. S9E and S9F). But regardless, when in the hinge-latch mode of operation, the open:closed ratio of KEAP1-NRF2 complexes increases in response to high *CLASS*_I-V_ levels (Figs. 4A-4B and S9A-S9B), as opposed to the expected decrease, suggesting that the hinge-latch hypothesis may not be valid for class I-V compounds-induced NRF2 activation. Indeed, using titration NMR spectroscopy, the most recent study by Yamamoto group clearly demonstrated that modifications of reactive cysteines of KEAP1 by class I-V oxidants and electrophiles, including CDDO-Im and sulforaphane targeting Cys151 and 15d-PGJ2 targeting Cys288, do not break the DLG-mediated binding (Horie et al. 2021).

Class VI compounds are those that can bind to the DC region of KEAP1 and thus disrupt DLG-mediated, and also potentially, ETGE-mediated NRF2 binding. Therefore, the NRF2 stabilization effect of class VI compounds is indirect, by shifting KEAP1-NRF2 complex away from the ubiquitinatible closed state. An endogenous ligand is p62, which has a KEAP1-interacting region (KIR) containing a DPSTGE motif that is similar to the ETGE motif of NRF2 (Lau et al. 2010, Jiang et al. 2015). The motif has similar or even higher binding affinities for KEAP1 than the DLG motif of NRF2, depending on its phosphorylation status (Komatsu et al. 2010, Ichimura et al. 2013). Small-molecule compounds have also been identified recently as disruptors of the protein-protein interaction between KEAP1 and NRF2, such as Cpd16 (Jiang et al. 2014) and several others (Yasuda et al. 2017, Lazzara et al. 2020, Lee and Hu 2020). By displacing DLG binding preferentially, these compounds make the KEAP1-NRF2 complex function as a hinge-latch as recently demonstrated experimentally (Horie et al. 2021). Our Model 3b captures the hinge-latch behavior in response to class VI NRF2 activators (Fig. 6), and shows that the open:closed ratio of KEAP1-NRF2 complex actually decreases with increasing concentrations of class VI activator (Fig. 6E). Simulations of Model 3b also suggest that when the two monomeric subunits of KEAP1 dimer can both be occupied by a class VI compound, NRF2 activation is more robust because of simultaneous sequestration of free KEAP1 by the compound (Figs. 6D vs. 6F). Horie et al. indeed showed that with high enough concentrations, p62 and small-molecule class I-V compounds can completely dissociate NRF2 from KEAP1 dimers, breaking the ETGE-mediated hinge (Horie et al. 2021). A recently identified endogenous protein, FAM129B, has both DLG and ETGE motifs on the C terminal and can compete with NRF2 for KEAP1 binding (Cheng 2019). FAM129B is found to be upregulated in many cancers which have poor prognosis by promoting NRF2 activation and thus chemoresistance.

### Maximal NRF2 activation, ultrasensitivity, and floodgate hypothesis

The maximal fold increase of total NRF2 is determined by the differential half-lives at the basal vs. severely stressed conditions. With a basal half-life of about 10 min in the cytosol in our models, and the lengthening of the half-life to over an hour under simulated oxidative stress such as in Model 3a, total NRF2 increases by 5-fold (Fig. 2D and Tables S8-S9). As discussed above, under oxidative stress, this model switches from KEAP1-mediated degradation to KEAP1-mediated stabilization of NRF2, therefore the fold-increase can be even higher when parameter *k’*_6_ is of lower values (Figs. S5B and S5D). Conversely, the fold-increase becomes smaller when *k’*_6_ is higher (Fig. S5F). It is also evident that at the basal condition when there is a higher fraction of the closed KEAP1-NRF2 complex that is rapidly degraded, the system is poised to produce higher levels of NRF2 in response to stresses, as the closed complex becomes stabilized by a class I-V compound or dissociated by a class VI compound. In Model 3b which simulates class VI compounds, because there is no closed KEAP1-NRF2 complex with a *CLASS*_VI_ molecule attached, the maximal fold-increase of total NRF2 is limited by the half-lives of free NRF2 and NRF2 in the open KEAP1-NRF2 complexes. In our study parameters *k*_5_, *k’*_5_, *k*_9_, *k’*_9_, *k*_9.1_ and *k’*_9.1_ govern the degradation of these NRF2 species, which have an equal half-life of 40 min. As a result, the maximal fold-increase of total NRF2 in Model 3b cannot exceed 40/10 = 4-fold (Fig. 6B and Tables S8-S9).

As aforementioned in Introduction, for an adaptive stress response, it is ideal that some degree of signal amplification, i.e., ultrasensitivity, can be embedded in the feedback circuit to ensure robust resistance to perturbation. In Models 1, 2, and 3a, free NRF2 exhibits some decent ultrasensitivity. Molecular mechanisms producing ultrasensitivity can arise from six common ultrasensitive motifs (Zhang et al. 2013). In the KEAP1-NRF2 module here, it appears that both zero-order degradation and protein sequestration (molecular titration) are at play simultaneously to produce NRF2 ultrasensitivity, where the sequestration is mediated by ETGE binding and zero-order degradation is mediated by saturation of DLG binding. As shown in Figs. S3G-S3H and 2H-2I, KEAP1-mediated degradation of NRF2 in the closed KEAP1-NRF2 complex will eventually saturate when all KEAP1 dimers are occupied by NRF2. Around this saturation point, KEAP1-mediated NRF2 degradation becomes zero order such that any additional increase in NRF2 will have to rely on KEAP1-independent mechanism to degrade. As a result of the nonlinear, zero-order degradation, the steady-state total NRF2 abundance may experience some steep changes around the point of KEAP1 saturation than when no saturation occurs. Indeed, for Models 1-3b, the *n*_H_ of total NRF2 is between 1.17-1.35. But with *LRC*_max_ between 0.35-42, total NRF2 does not exhibit overt ultrasensitivity because of the high basal level. From the perspective of free NRF2, this KEAP1 saturation point is also a moment when free NRF2 can no longer be sequestered by KEAP1, and as a result any additional NRF2 synthesized *de novo* will remain as free NRF2, leading to a steep increase in its abundance. Therefore, both zero-order degradation (by providing more NRF2 overall) and protein sequestration are at play simultaneously to produce the ultrasensitivity of free NRF2.

Conceivably, the abundance of total KEAP1 and whether NRF2 can accumulate to a level that surpasses this abundance play a critical role in quantitative NRF2 activation. If total NRF2 can never increase to a level higher than KEAP1 dimer, then NRF2 cannot escape the sequestration by KEAP1 and there will be no ultrasensitivity of free NRF2. This is first illustrated by setting *k’*_6_ to a higher value such that total NRF2 can only barely match the level of total KEAP1 (Figs. S4F and S5F). The intracellular NRF2:KEAP1 ratio at basal conditions varies among different cell types, which can be lower or higher 1:1 (Khalil et al. 2015, Iso et al. 2016). Since nuclear NRF2 often constitutes a considerable fraction of total NRF2 at basal conditions (Khalil et al. 2015, Iso et al. 2016), the cytosolic NRF2:KEAP1 ratio can be actually even lower than the values reported for the whole cells. Varying the abundance of KEAP1 in the models has some interesting results. By increasing KEAP1, its role in further destabilizing Nrf2 is limited because it is already in excess, and as a result total NRF2 does not increase further (Fig. 5A). But increasing KEAP1 will be more effective as a sequester to inhibit Nrf2. There seems to be an optimal KEAP1 abundance that can produce the steepest *NRF2*_free_ response (Fig. 5B). When KEAP1 is too low, NRF2 is constitutively activated, but when KEAP1 is too high, free NRF2 is constitutively suppressed.

This sequestering role of KEAP1 is consistent with the floodgate hypothesis which postulates that stabilization of NRF2 due to loss of KEAP1 activity as an E3 ligase adaptor protein is not sufficient to initialize NRF2 nuclear translocation; NRF2 has to accumulate to a higher level to overflood the KEAP1 gate to move to the nucleus (Yamamoto et al. 2018). A potential caveat of this mechanism is that it takes some time to produce enough NRF2 to saturate KEAP1, therefore the free NRF2 response can be delayed as we demonstrated with our models. However, it is likely that at the basal condition, the cytosolic NRF2:KEAP1 ratio is near parity in some cells such that KEAP1 is near saturation. As a result, the system is at a tipping point, poised to respond to a slight increase in oxidative stress to overwhelm KEAP1 and cause an immediate and steep increase in free NRF2 (Baird et al. 2013).

While both Models 2 and 3a are cycle models, the ultrasensitivity of free NRF2 exhibited by Model 3a, which has two-step ETGE binding, is somehow weaker than Model 2. As described in Results, this is partly because a higher free NRF2 level is required to maintain the same turnover fluxes through different NRF2 species (Figs. 2H and 2I) at the basal steady state in Model 3a, resulting in a lesser zero-order degradation effect. The apparent dissociation constant for the ETGE-mediated two-step binding is 7.54 nM, which is lower than the 20 nM used in Model 2. However, at the basal condition, free NRF2 is higher in Models 3a than Model 2 as a result of the two-step binding and slow fluxes through the second-step binding. A higher basal level will always reduce the degree of ultrasensitivity (Zhang et al. 2013), despite that Models 2 and 3a have comparable maximally induced free NRF2 levels (Table S9). By increasing the binding affinity of ETGE, e.g., through increasing *k*_1_ and *k*_1.1_, to reduce basal free NRF2, the ultrasensitivity of Model 3a is improved dramatically (Figs. 3B and 3D).

With both *n*_H_ and *LRC*_max_ of free NRF2 close to unity, the ultrasensitivity for class VI compounds as in Model 3b is basically absent. In contrast to class I-V compounds, a class VI compound does not need to induce total NRF2 to a level that exceeds total KEAP1 to produce tangible increase in free NRF2. This is because when a class VI compound can bind to both of the monomeric subunits of KEAP1 dimer, it would titrate free KEAP1 away, essentially lowing the amount of available KEAP1 that can sequester NRF2. This lowering of the “floodgate” can result in a much higher level of free NRF2 that can be maximally induced by class VI compounds than class I-V compounds (Figs. 6D vs. 2G and Table S9). However, because of the reduced sequestration by KEAP1, the ultrasensitivity of free NRF2 is lost.

### Response of nuclear NRF2

To transcriptionally regulate its target genes, NRF2 needs to translocate into the nucleus where it dimerizes with sMaf to gain affinity for the AREs in promoters. The flux of nuclear translocation constitutes a load to the cytosolic NRF2. At a constant NRF2 production rate in the cytosol, this nuclear load is expected to alter the dynamics of NRF2 activation. With Models 4a and 4b we made the assumptions that KEAP1 and NRF2 interactions in the nucleus follow the same kinetic parameters as in the cytosol except that nuclear KEAP1 is not able to mediate the ubiquitination and degradation of NRF2 and is not subject to redox modification by class I-V compounds or binding by class VI compounds. With higher abundance of nuclear NRF2 than KEAP1, as observed in RAW 264.7 cells and potentially many other cell types (Iso et al. 2016), nuclear KEAP1 is nearly saturated by NRF2, resulting in low basal free nuclear KEAP1 dimer and high free nuclear NRF2 levels. These configures result in a net nuclear influx of NRF2 that is 22% of the NRF2 production rate in the cytosol (Table S10). Therefore, net nuclear importing of NRF2 constitutes a significant load of NRF2 production. It can thus be estimated that even under oxidative stress that completely terminates cytosolic NRF2 degradation and all cytosolic NRF2 translocates into the nucleus, total nuclear NRF2 cannot increase by > 5-fold. Our simulations confirmed this prediction (Figs. 7E and 8F, Tables S8-S9), and the fold increase of nuclear free NRF2 is only slightly higher than the total. Class VI activators seem to has a larger effect on maximal nuclear NRF2 (4.2-fold) than class I-V activators (2.6-fold). This is due to the KEAP1-titrating effects of a class VI activator, which reduces free cytosolic KEAP1, pushing more NRF2 into the nucleus. Nonetheless these lessor responses contrasts with the nearly 10-fold increase in nuclear NRF2 under exposure to DEM at 100 µM in RAW 264.7 cells which our model is partially based upon (Iso et al. 2016). For nuclear Nr2 to increase to higher levels, additional mechanisms have to be at play which are not included in our models. These include (i) increased NRF2 production through transcriptional autoregulation, which has been confirmed in many cell types including RAW 264.7 cells (Kwak et al. 2002, Pi et al. 2003, Pi et al. 2008); (ii) reduced nuclear exporting of NRF2 due to redox-sensitive cysteine modification of the nuclear export signal (NES) sequence in the Neh5 domain of NRF2 (Li et al. 2006); (iii) stabilization of nuclear NRF2 under oxidative stress; (iv) lower nuclear NRF2 load at the basal condition such that there is still more reserve capacity for nuclear NRF2 accumulation.

Free nuclear NRF2 does not exhibit overt ultrasensitivity in either of the two models. Part of the reason is due to its high basal level and smaller fold increase of total nuclear NRF2 discussed above. However, a number of mechanisms that can potentially contribute to ultrasensitivity have been confirmed in the KEAP1-NRF2 system. These mechanisms include (i) positive transcriptional autoregulation of both NRF2 and sMaf (Kwak et al. 2002, Katsuoka et al. 2005), (ii) molecular titration of sMaf by inhibitor Bach1 (Igarashi et al. 1998), (iii) positive feedback through NRF2 induction of p62 which can titrate KEAP1 away from NRF2 and also promote KEAP1 autophagy (Katsuragi et al. 2016), and (iv) multi-step signaling through (a) enhanced nuclear NRF2 accumulation due to redox modification of NES as mentioned above (Li et al. 2006) and (b) redox-sensitive nuclear exporting of Bach1 (Suzuki et al. 2003, Dhakshinamoorthy et al. 2005). It is highly likely that these mechanisms converge to produce ultrasensitive nuclear free NRF2 accumulation.

### Limitations

The KEAP1-NRF2 module has been modeled mathematically as part of larger networks. We have constructed NRF2-mediated pathways of antioxidant induction and phase II enzyme induction, containing negative feedback, incoherent feedforward, and a variety of ultrasensitive motifs to understand the nonlinear dose-response relationship under oxidative stress (Zhang and Andersen 2007, Zhang et al. 2009). Blis and his colleagues adapted these models to interpret and predict antioxidant gene induction in human renal cells in response to cyclosporine (Hamon et al. 2014), and glutathione depletion in liver microfluidic chips in response to flutamide (Leclerc et al. 2014). Khalil et al. constructed a model of KEAP1-NRF2/sMaf-ARE activation and its interaction with the peroxiredoxin and thioredoxin antioxidant enzymes in controlling intracellular H_2_O_2_ levels and regulating the reduction of KEAP1, in which one-step ETGE binding was considered (Khalil et al. 2015). Xue et al. observed a basal NRF2 cytosol-nucleus oscillation behavior in cells with a period of about 2 hours for which they constructed a mathematical model of negative feedback through NRF2 phosphorylation and dephosphorylation without involving changes in the abundance (Xue et al. 2015). Kolodkin et al. has recently incorporated the KEAP1-NRF2 component into an ROS dynamic network to explore the design principles relevant to network-based therapies for Parkinson disease (Kolodkin et al. 2020). Compared to the previous work, our present study provided a much more detailed analysis of the KEAP1-NRF2 module itself, which can be adapted and included in future systems-level models of antioxidant and detoxification responses.

There are several limitations of the present study, however. In the models we have limited the action of class I-V and VI compounds on the KEAP1 molecules in the cytosol only, however it is possible that these compounds, especially class VI, may still compete for KEAP1 in the nucleus to further drive NRF2 activation. We have assumed separate pools of cytosolic and nuclear KEAP1 without exchange. However, it has been shown that KEAP1 may also control postinduction repression of the NRF2-mediated antioxidant response by escorting NRF2 out of the nucleus (Sun et al. 2007). The DLG-mediated binding kinetics has been measured *in vitro* with peptide fragments of KEAP1 and NRF2 as inter-molecular event following the law of mass action. Measuring the binding kinetics as an intra-molecular event as occurring with full-length proteins will help reduce the uncertainty of model parameterization. It is also unclear whether the modification of cysteine residues of both KEAP1 subunits of the dimer will have any differential effects on NRF2 ubiquitination than only one subunit is modified. Lastly, the parameterization and calibration of our models are based on experimental measurements from multiple cell types, such as RAW 264.7 and HEK293 cells, and under various experimental conditions. Therefore, the parameter values and model responses do not represent an ideal “average” cell. However, we systemically varied parameters where applicable in our study to explore their effects on NRF2 response. All in all, future iterations of the KEAP1-NRF2 model should address these limitations as more quantitative information, such as binding and degradation kinetics of all NRF2 forms in complex with KEAP1, is obtained.

### Conclusions

Robustly inducing antioxidant and detoxification genes to cope with cellular stress imposed by oxidative and electrophilic chemicals requires timely and sufficient NRF2 accumulation and translocation into the nucleus. KEAP1 plays a dual role in repressing NRF2 – promoting its degradation to keep its total abundance low and sequestering to keep its free abundance low. The floodgate hypothesis captures some of the dual actions of KEAP1 (Iso et al. 2016, Suzuki and Yamamoto 2017). Our modeling revealed here that the quantitative aspect of protein stabilization of NRF2 and nuclear translocation can be better understood as a water tank model that overflows due to drain closure (Fig. 9A), which we believe is an improvement over the floodgate analogy. Here, the water is poured into the large water tank at a constant rate, just as NRF2 is produced in the cytosol. Since the drain is open, most of the water will leave the tank with a small amount remaining and leaking to the small tank (nucleus). These events are like NRF2 being actively degraded by KEAP1 and cytosolic and nuclear NRF2 levels are low. If a stopper is partially put in place, the water will drain slowly, and the water level in the large tank will rise, however it is still being held by the large tank without much going into the small tank. This is like under mild stress, KEAP1-dependent NRF2 degradation is partially stopped, total NRF2 will increase but because of the sequestration by KEAP1, it still remains largely in the cytosol. Therefore, the height of the large water tank here is equivalent to the total cytosolic KEAP1 dimer. When the stopper is further pushed in to completely block the drain, the water level will rise and eventually overflow the large tank and flood the small tank. This is like under severe oxidative stress, KEAP1-mediated NRF2 degradation is totally shut down, NRF2 accumulates to a level exceeding cytosolic KEAP1 dimer, and free NRF2 rises sharply and translocates into the nucleus. Modification of KEAP1 cysteine by class I-V compounds is like slowing the drain without affecting the height of the large water tank, while binding of class VI compounds to both of the two subunits of KEAP1 dimer is like simultaneously slowing the drain and lowering the height of the large water tank. The differential action of this water-tank model can be captured by a reduced mathematical model of KEAP1-NRF2 interaction (Fig. 9B). The free nuclear NRF2 response to class I-V compounds is potentially more ultrasensitive than that to class VI compounds, while at lower concentrations class VI compounds may activate more nuclear NRF2 than class I-V compounds (Fig. 9C).

**Figure 9.**
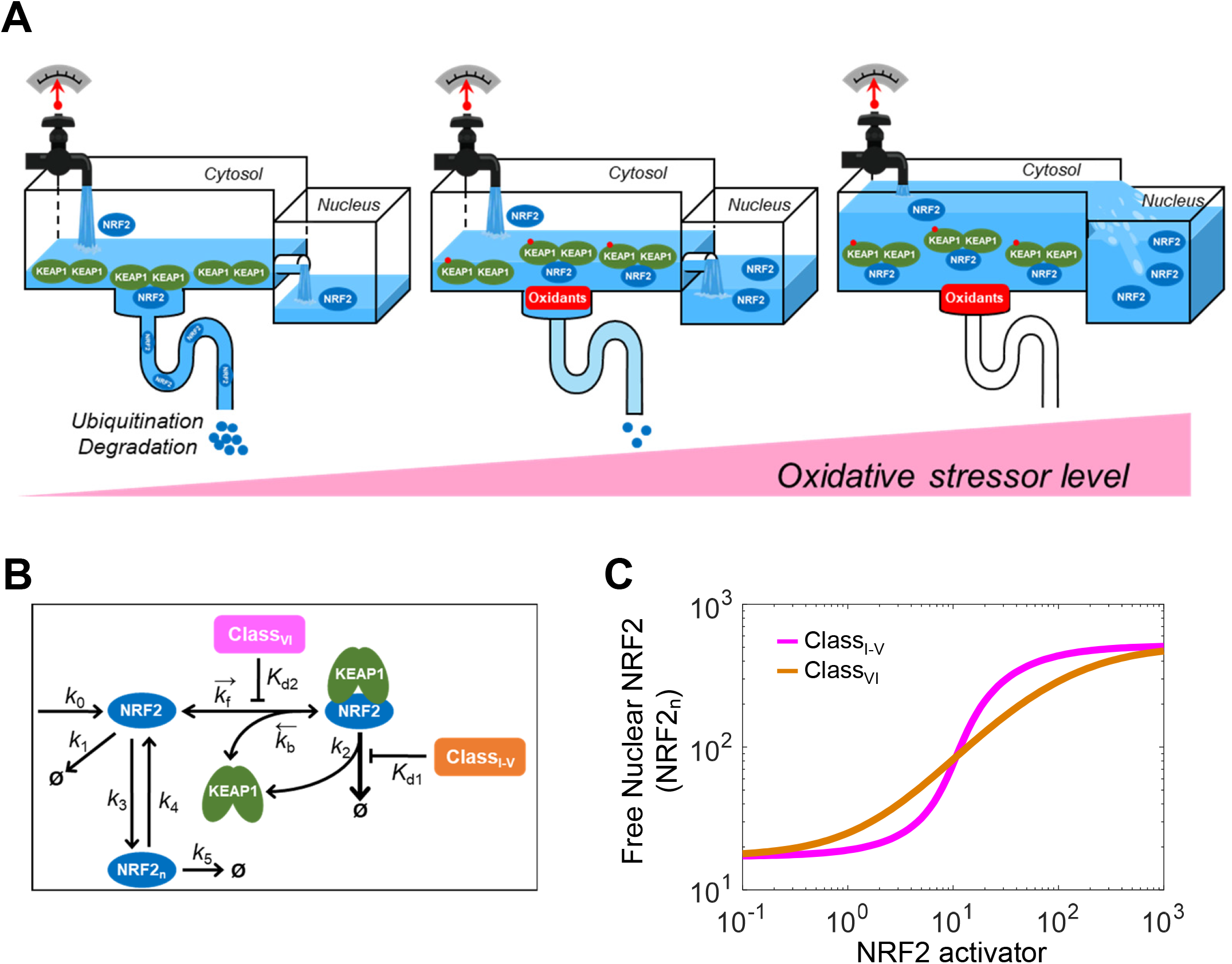
Water-tank analogy and reduced KEAP1-NRF2 mathematical model. **(A)** Schematic illustration of the water-tank analogy for KEAP1-dependent NRF2 degradation, sequestration, and nuclear translocation. Large tank: cytosol, small tank: nucleus, height of large tank: total cytosolic KEAP1 abundance, water: NRF2, tap: NRF2 production, drain: KEAP1-mediated NRF2 degradation, stopper: oxidant or NRF2 inducer. To reduce clutter for clarity, KEAP1-independent NRF2 degradation and nuclear NRF2 degradation are not shown. **(B)** Reduced KEAP1-NRF2 mathematical model for NRF2 activation by class I-V and class VI activators. **(C)** Predicted differential free nuclear NRF2 dose-response for class I-V and class VI activators.

Quantitative understanding of NRF2 activation can have many implications. A detailed kinetic model like the one we presented here can help to explore the systems behavior of cellular oxidative stress responses. It may help to better understand cancer chemoresistance, where mutation in either NRF2 or KEAP1 can lead to constitutive NRF2 activation or a more prompt and robust activation in response to chemo-drugs. Our modeling suggests that threshold concentrations may exist for certain class I-V compounds and at suprathreshold concentrations they can readily activate NRF2, while to finely control NRF2 activation with precision, class VI compounds are preferred. The model may thus help with using a synthetic biology approach to improve current and design novel classes of NRF2 activators or inhibitors. The mechanistically-based KEAP1-NRF2 model can also help to understand the commonality of toxic actions of environmental oxidative stressors.

## Supporting information

Model Code in Berkeley Madonna and R

## ACKNOWLEDGEMENTS

This research was supported in part by the National Natural Science Foundation of China: 81830099 (J.P.), 82020108027 (J.P.) and 81602824 (S.L.); Liaoning Key Research and Development Guidance Plan 2019JH8/10300012 (J.P.); NIEHS Superfund Research grant P42ES04911, and NIEHS HERCULES grant P30ES019776.

## CONFLICT OF INTEREST

The authors declare no conflict of interest.

## Notes

### Competing Interest Statement

The authors have declared no competing interest.

### Summary of Updates

(1) The results for Models 1 and 2 and two corresponding figures are moved to Supplemental Material to reduce the main text length. Instead, a summary of Models 1 and 2 is provided in the main text. (2) R codes for the Models are provided. (3) Some editorial changes are made to Abstract and Discussion.

