## Supplemental Materials for "Mathematical Modeling Reveals Quantitative Properties of KEAP1-NRF2 Signaling"

### Results for Models 1 and 2

#### Model 1 (Equilibrium Mode)

Model 1 is the minimal model, involving only basic cytosolic KEAP1 and NRF2 interactions through ETGE and DLG motifs (Fig. 1). At the basal steady state, due to the strong binding between KEAP1 and NRF2 through ETGE, the majority of NRF2 is sequestered by KEAP1, leaving *free NRF2* ($NRF2_{free}$), at 2 nM, just a tiny fraction of *total NRF2* ($NRF2_{tot}$), which is at 150 nM (Fig. S1A and Table S8). The open state of the KEAP1-NRF2 complex ($KEAP1\_NRF2_{open}$), in which the association is through ETGE only, and the closed state of the complex ($KEAP1\_NRF2_{closed}$), in which the association is through both ETGE and DLG, are equal to each other in concentration at 74 nM and much higher than $NRF2_{free}$. The comparable levels between the open and closed states are consistent with what was observed experimentally in HEK293 cells (Baird et al. 2013). When the synthesis of NRF2 is terminated by setting $k_0$=0, all NRF2 species including $NRF2_{free}$, $KEAP1\_NRF2_{open}$ and $KEAP1\_NRF2_{closed}$, degrade exponentially, and the half-life of $NRF2_{tot}$ is 10 min (Fig. S1A).

To examine the basic behavior of the model when NRF2 in $KEAP1\_NRF2_{closed}$ is stabilized, as would occur during oxidative stress, we first lowered $k_6$ to different values, while keeping *class I-V activator* at zero ($CLASS_{I\text{-}V}$=0) for simplicity. As $k_6$ decreases from the default 2.03E-3 $S^{-1}$ (equivalent half-life $t_{1/2}$=5.7 min for $KEAP1\_NRF2_{closed}$) to 1.178E-4 (which is the default value of $k'_6$ for degradation of $KEAP1_o\_NRF2_{closed}$, equivalent $t_{1/2}$=98 min), all NRF2 species increase and reach steady states in about 300 min (Fig. S1B). $KEAP1\_NRF2_{open}$ and $KEAP1\_NRF2_{closed}$ reach the steady states the fastest, follows by $NRF2_{free}$ and $NRF2_{tot}$. There is an apparent delay in the $NRF2_{free}$ response. $NRF2_{tot}$ increases by 5-fold, from 150 to 750 nM, while $NRF2_{free}$ increases by a much greater fold, from 2 to 230 nM. At this activated state, by setting $k_0$=0, all NRF2 species degrade but at different rates with $NRF2_{free}$ disappearing much more quickly, and the half-life of $NRF2_{tot}$ is 54 min (Fig. S1B). By setting $k_6$ to an even lower value (0.589E-4), the steady-state levels of both $NRF2_{free}$ and $NRF2_{tot}$ increase but only to a limited extent, and the half-life of $NRF2_{tot}$ lengthens to 65 min (Fig. S2A). When $k_6$ is lowered to zero, mimicking complete shutoff of KEAP1-mediated NRF2 degradation, $NRF2_{tot}$ only increases to 858 nM, a 5.7-fold increase from the basal level and its half-life lengthens to 78 min, while $NRF2_{free}$ increases by 117-fold (Fig. S2C). The behavior when $k_6$=0 represents the maximal response Model 1 can be induced. We next examined the steady-state dose-response behavior of Model 1 by varying the $CLASS_{I\text{-}V}$ level. With $k'_6$ at the default value, $NRF2_{free}$ exhibits an ultrasensitive, sigmoidal dose-response with Hill coefficient ($n_H$) of 2.02 and maximal local response coefficient ($LRC_{max}$) of 1.92 (Figs. S1C in dual-log scale and S1D in dual-linear scale). $NRF2_{tot}$ is subsensitive with a much shallower dose-response curve. Setting $k'_6$ to lower values increases the ultrasensitivity of $NRF2_{free}$ slightly as its maximal steady-state level increases (Figs. S2B and S2D).

An interesting feature of Model 1 is that the open and closed KEAP1-NRF2 complexes ($KEAP1\_NRF2_{open}$ and $KEAP1\_NRF2_{closed}$) behave in an almost synchronized fashion in that their abundance ratio remains at 1:1 at all times in all conditions (Figs. S1A-S1C and S2), suggesting these two species are always at equilibrium to each other. Using FRET to track the open and closed states of KEAP1-NRF2 complex, Baird et al. observed that the two states diverge and do not follow an equilibrium mode of operation in a variety of chemically perturbed conditions (Baird et al. 2013). But rather, a "cyclic sequential attachment and regeneration" (abbreviated as "cycle") mode of operation was suggested. In this mode, because of the rapid degradation of NRF2 in the closed KEAP1-NRF2 complex, KEAP1 is quickly released (or regenerated) to join the free KEAP1 dimer pool and sequester newly synthesized NRF2 again, thus completing a global cycle for KEAP1. Under oxidative stress, this cycle is blocked as the NRF2 degradation-coupled release of KEAP1 from the closed KEAP1-NRF2 complex is inhibited, leading to accumulation of the closed state and depletion of free KEAP1 dimer. Therefore, in the next section, we evolved Model 1 into Model 2 such that the model behavior is aligned with the cycle mode of operation.

**Model 2 (Cycle Mode)**

Examining the basal steady-state behaviors of Model 1 revealed that the two fluxes of the reversible conversion between $KEAP1\_NRF2_{open}$ and $KEAP1\_NRF2_{closed}$ are comparable to each other ($flux_{k3}$ = 14.645 and $flux_{k4}$ = 14.495 nM/S) and overwhelmingly dominant over the connected turnover fluxes (> 96-fold of $flux_{k6}$ and $flux_{k9}$) (Table S10). The predominantly high $flux_{k3}$ and $flux_{k4}$ explain why the open and closed KEAP1-NRF2 complexes in Model 1 behave in an equilibrium mode of operation. To convert it to a cycle mode, we reduced the parameter values of $k_3$ and $k_4$. When $k_4$ is reduced to about 1.96E-4 $S^{-1}$ or lower (simultaneously reducing $k_3$ to keep the open:closed ratio at 1:1 at the basal condition), the behaviors of $KEAP1\_NRF2_{open}$ and $KEAP1\_NRF2_{closed}$ start to separate appreciably. As detailed in Table S1 footnote, Model 2 was finally configured with $k_4$=1.0E-4 as the default value and $k_0$, $k_3$ and $k_6$ adjusted accordingly to maintain the same basal $NRF2_{tot}$ level and half-life as Model 1. As a result, the basal $flux_{k3}$ = 0.135, $flux_{k4}$ = 7.35E-3, and $flux_{k6}$ = 0.128 nM/S approximately (Table S10), indicating that the majority of NRF2 moving from the open to closed state through the $k_3$ step is degraded within $KEAP1\_NRF2_{closed}$ through the $k_6$ step, and only a small fraction (5.5%) returns to the open state through the $k_4$ step.

Fig. S3A shows the behaviors of NRF2 species decaying from the basal steady state when setting $k_0$=0. While the half-life of $NRF2_{tot}$ is still 10 min, the levels of $KEAP1\_NRF2_{open}$ and $KEAP1\_NRF2_{closed}$ diverge quickly with the open state decaying much faster than the closed state. By 15 min the open:closed ratio is about 1:2.7, comparable to what was observed in HEK293 cells treated with cycloheximide (Baird et al. 2013). By setting $k_4$ to lower values than the default and in the extreme case $k_4$=0 such that the binding between KEAP1 and DLG becomes irreversible (and $k_0$, $k_3$ and $k_6$ were adjusted accordingly as above), the divergent behaviors of $KEAP1\_NRF2_{open}$ and $KEAP1\_NRF2_{closed}$ are similar to Fig. S3A (simulation results not shown).

To examine the basic behavior of the model when NRF2 in $KEAP1\_NRF2_{closed}$ is stabilized, $k_6$ was lowered to different values, while keeping $CLASS_{I\text{-}V}$=0. As $k_6$ decreases from the default 1.74E-3 $S^{-1}$ (equivalent $t_{1/2}$=5.7 min) to 1.454E-4 (which is the default value of $k'_6$, equivalent $t_{1/2}$=79 min), all NRF2 species reach steady states in about 400 min (Fig. S3B). $KEAP1\_NRF2_{open}$ and $KEAP1\_NRF2_{closed}$ quickly diverge with $KEAP1\_NRF2_{closed}$ increasing and reaching the steady state in about 100 min, while $KEAP1\_NRF2_{open}$ initially increases slightly but then decreases to a level slightly lower than the basal level. The open:closed ratio decreases and reaches about 1:4.6 at 1 h, concordant with what was observed experimentally in HEK293 cells treated with proteasomal inhibitor MG132 or chemical stressors such as sulforaphane and sulfoxythiocarbamate alkyne (Baird et al. 2013). $NRF2_{tot}$ increases from 150 to 750 nM, while $NRF2_{free}$ increases by a much greater fold, from 2 to 223 nM. At this activated state, by setting $k_0$=0, all NRF2 species degraded, with a half-life of 68.5 min for $NRF2_{tot}$, while $NRF2_{free}$ seems to disappear much more quickly approaching the zero level within 1 h (Fig. S3B). By setting $k_6$ to even lower values, the maximal levels of both $NRF2_{free}$ and $NRF2_{tot}$ increase but to a limited extent and the half-life of $NRF2_{tot}$ lengthens to 191 min in the extreme case when $k_6$=0 (Figs. S4A and S4C). Interestingly, the decay of $NRF2_{tot}$ starts to become biphasic. The first fast phase is due to rapid $NRF2_{free}$ drop, and the second slow phase follows the decay of $KEAP1\_NRF2_{closed}$.

We next examined the dynamical responses of Model 2 to a range of $CLASS_{I\text{-}V}$ levels. $NRF2_{tot}$ increases to higher steady-state levels with increasing $CLASS_{I\text{-}V}$ levels, and the time it takes to reach steady states also increases (Fig. S3C), which is concordant with the lengthening of the half-life as more NRF2 is diverted to the more stable, closed state complex. In comparison, there is a considerable delay in the response of $NRF2_{free}$, which does not rise tangibly above the basal level until after 60 min (Fig. S3D). After the initial delay, the rising time of $NRF2_{free}$ becomes shorter with higher $CLASS_{I\text{-}V}$ levels. The initial delay is caused by the sequestration of newly synthesized NRF2 by *free KEAP1 dimer* ($KEAP1_{free}$), the level of which decreases quickly as it forms complexes with NRF2 (Fig. S3E).

The steady-state $NRF2_{free}$ level exhibits an ultrasensitive response with respect to $CLASS_{I-V}$ levels, with $n_H$ of 2.62 and $LRC_{max}$ of 3.09 (Fig. S3F). Interestingly, unlike steady-state $KEAP1\_NRF2_{closed\_tot}$ ($KEAP1\_NRF2_{closed}$ + $KEAP1_o\_NRF2_{closed}$) which increases monotonically with $CLASS_{I-V}$ levels, steady-state $KEAP1\_NRF2_{open\_tot}$ ($KEAP1\_NRF2_{open}$ + $KEAP1_o\_NRF2_{open}$) exhibits a nonmonotonic dose-response profile (Fig. S3F, green line). The peak coincides with the juncture of KEAP1 saturation at which point $KEAP1_{free}$ is nearly depleted and $NRF2_{free}$ increases sharply. The decrease in $KEAP1\_NRF2_{open\_tot}$ at higher $CLASS_{I-V}$ levels is due to increasing $flux_{k5}$ associated with increasing $NRF2_{free}$, which takes away from the net flux toward the KEAP1-NRF2 complexes (Table S11). When $k'_6$ is set to lower values, the degree of $NRF2_{free}$ ultrasensitivity is enhanced slightly, due to higher maximal $NRF2_{tot}$ and $NRF2_{free}$ levels that can be achieved (Figs. S4B and S4D), and the opposite occurs when $k'_6$ is high (Fig. S4F).

To analyze the mechanism of ultrasensitivity, we conducted flux analysis by artificially clamping $NRF2_{free}$ to different levels. $flux_{k5}$ increases linearly with the clamped $NRF2_{free}$ level, while $flux_{k9}$ and $flux_{k6}$ also increase but become saturated eventually because of the depletion of free KEAP1 dimer (Figs. 3G and 3H). The *total degradation rate* ($flux_{k5}$ + $flux_{k6}$ + $flux_{k9}$) exhibits an S-shape containing 3 phases. The initial rising phase is dominated by $flux_{k6}$ because $k_6$ is the highest compared with $k_5$ and $k_9$ and the closed state is higher in concentration. The second phase is slowly rising and nearly flat because of saturation of $flux_{k6}$ and to a small extent of $flux_{k9}$. The flatness of this second phase represents zero-order degradation, i.e., the *total degradation rate* is insensitive to changes in NRF2 levels. In the third phase, the *total degradation rate* rises again, because $flux_{k5}$ now becomes dominant. The intersection point between the *total degradation rate* and NRF2 *synthesis rate* ($flux_{k0}$) represents the steady state of $NRF2_{free}$. When

$k_6$=5.22E-4 (30% of default value) to mimic a mild stress condition, the intersection point appears at the junction of the first and second phases and the corresponding $NRF2_{free}$ is about 4 nM (Fig. S3G). When $k_6$ is lowered further to 1.74E-4 (10% of default) to mimic a more severe stress condition, the $flux_{k6}$ curve shifts to lower levels and as a result the second, flat phase of *total degradation rate* shifts downward as well (Fig. S3H). The intersection point between *total degradation rate* and NRF2 *synthesis rate* swings dramatically to the right, at the junction of the second and third phase, resulting in a much higher steady-state level of $NRF2_{free}$ at 175 nM. Therefore, a remarkable signal amplification is evident here – as $k_6$ decreases by only 3-fold (from 30% to 10% of default value), $NRF2_{free}$ increases by 43-fold. In the meantime, the steady-state $NRF2_{tot}$ exhibits no ultrasensitivity, as it increases from 326 to 701 nM, a 2.2-fold change only.

#### Figures

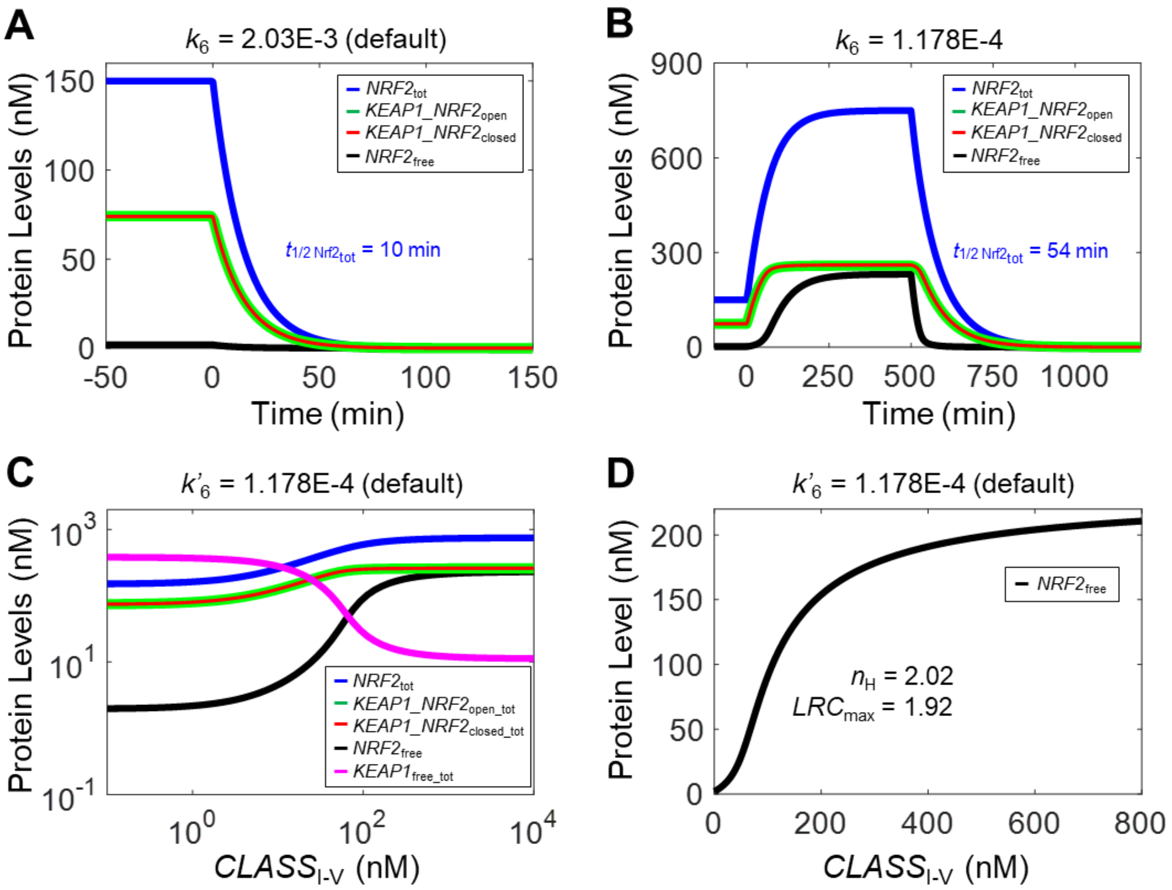

**Figure S1. Dynamical and steady-state behaviors of Model 1. (A)** Dynamical changes of basal $NRF2_{free}$, $KEAP1\_NRF2_{open}$, $KEAP1\_NRF2_{closed}$, and $NRF2_{tot}$ in response to termination of NRF2 synthesis (by setting $k_0$=0) starting at 0 min with $k_6$ at default value. **(B)** Dynamical changes of various NRF2 species in response to stabilization of NRF2 in $KEAP1\_NRF2_{closed}$ by setting $k_6$=1.178E-4 starting at 0 min and in response to termination of NRF2 synthesis (by setting $k_0$=0) starting at 500 min. For simulations in (A) and (B), $CLASS_{I\text{-}V}$ level is kept at zero. **(C)** Steady-state dose-response curves of various NRF2 species and $KEAP1_{free}$ for $CLASS_{I\text{-}V}$ on dual-log scale with $k'_6$ at default value. $n_H$ and $LRC_{max}$ for $NRF2_{total}$ are 1.17 and 0.40 respectively (not shown). **(D)** Steady-state dose-response curves of $NRF2_{free}$ in (C) on dual-linear scale illustrating sigmoidal shape, with $n_H$ and $LRC_{max}$ indicated.

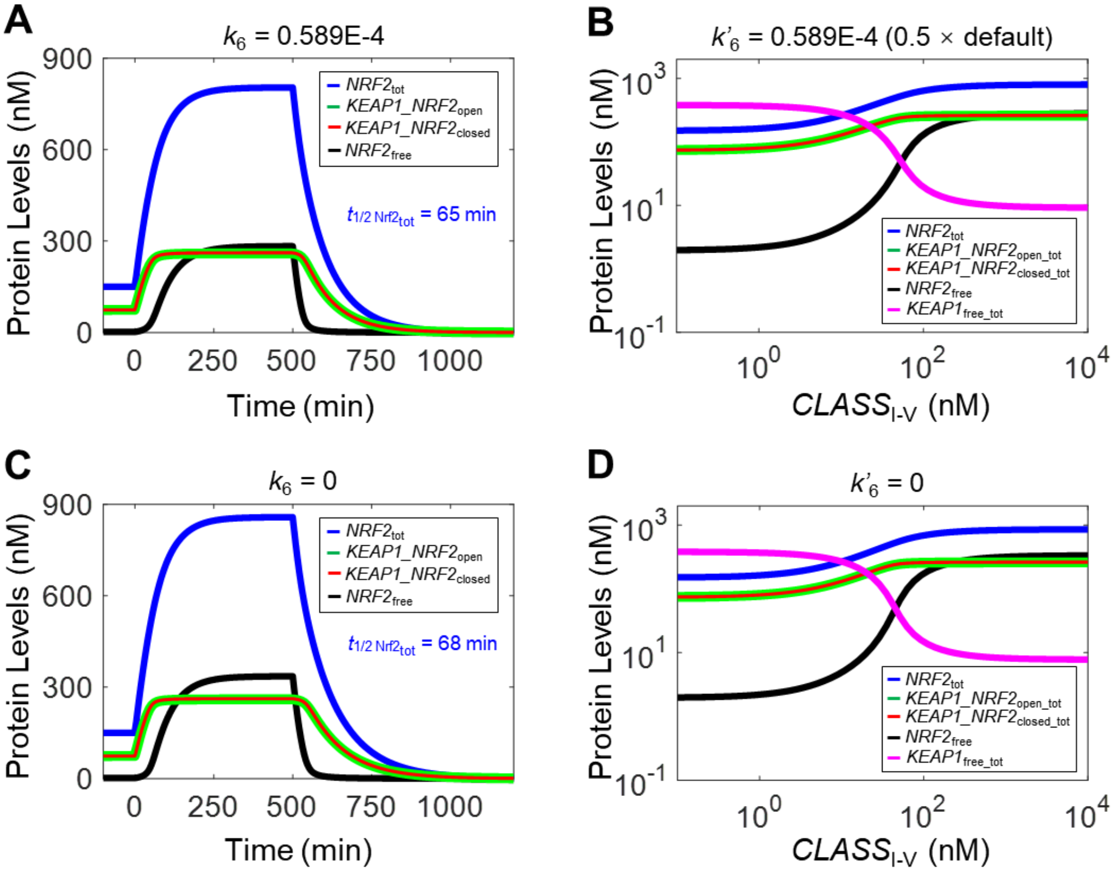

**Figure S2. Additional dynamical and steady-state behaviors of Model 1.** Dynamical changes of $NRF2_{free}$, $KEAP1\_NRF2_{open}$, $KEAP1\_NRF2_{closed}$, and $NRF2_{tot}$ in response to stabilization of NRF2 in $KEAP1\_NRF2_{closed}$ by setting **(A)** $k_6$=0.589E-4 and **(C)** $k_6$=0 starting at 0 min and in response to termination of NRF2 synthesis (by setting $k_0$=0) starting at 500 min. For simulations in (A) and (B), $CLASS_{I-V}$ level is kept at zero. Steady-state dose-response curves of various NRF2 species and $KEAP1_{free}$ with **(B)** $k'_6$=0.589E-4 and **(D)** $k'_6$=0.

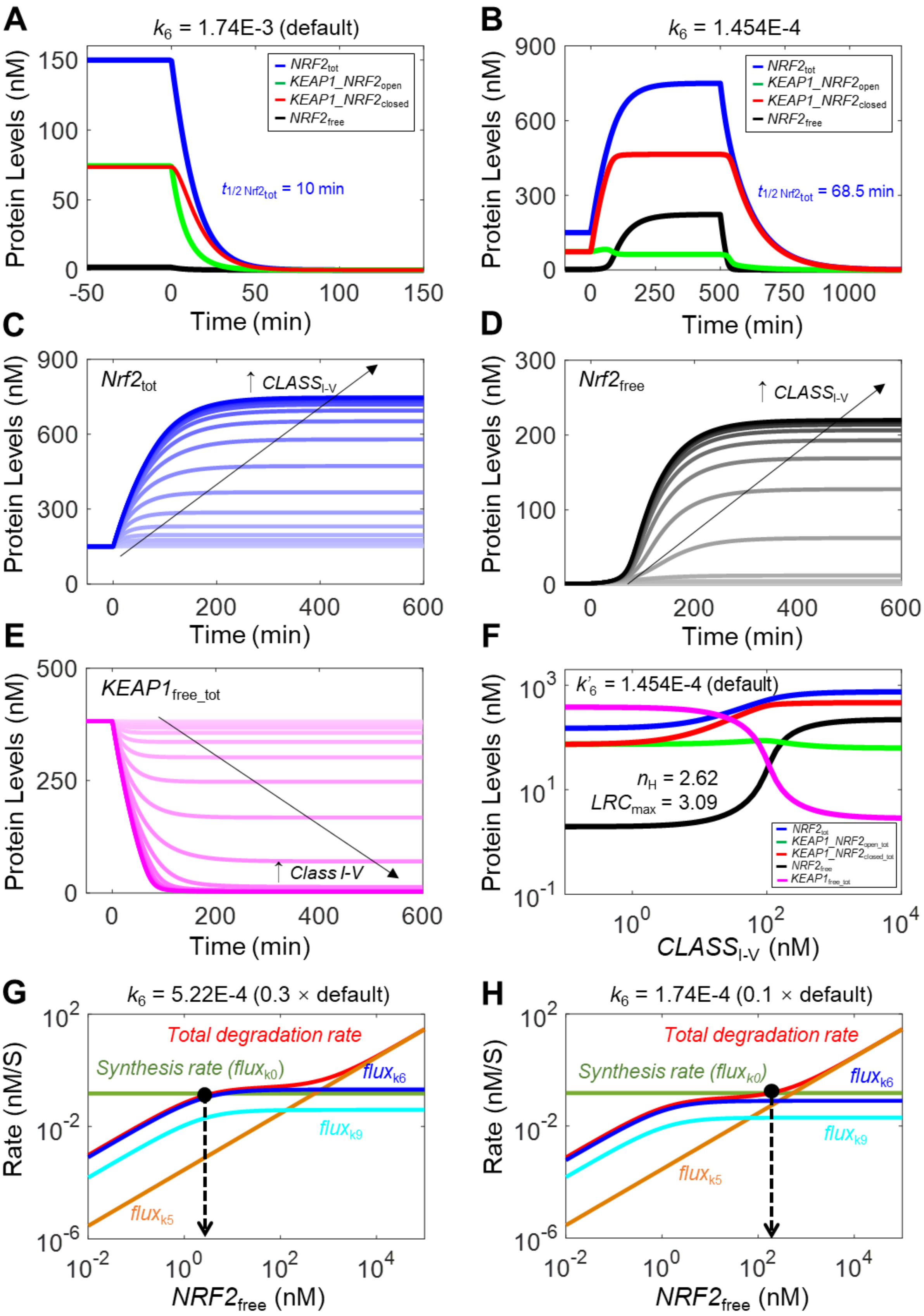
A
k6 = 1.74E-3 (default)
NRF2tot
KEAP1_NRF2open
KEAP1_NRF2closed
NRF2free
t1/2 Nrf2tot = 10 min
Protein Levels (nM)
Time (min)
B
k6 = 1.454E-4
t1/2 Nrf2tot = 68.5 min
C
Nrf2tot
↑ CLASSI-V
D
Nrf2free
E
KEAP1free_tot
↑ Class I-V
F
k'6 = 1.454E-4 (default)
nH = 2.62
LRCmax = 3.09
KEAP1_NRF2open_tot
KEAP1_NRF2closed_tot
KEAP1free_tot
CLASSI-V (nM)
G
k6 = 5.22E-4 (0.3 × default)
Total degradation rate
Synthesis rate (fluxk0)
fluxk6
fluxk9
fluxk5
Rate (nM/S)
NRF2free (nM)
H
k6 = 1.74E-4 (0.1 × default)

**Figure S3. Dynamical and steady-state behaviors of Model 2. (A)** Dynamical changes of basal *NRF2*$_{free}$, *KEAP1_NRF2*$_{open}$, *KEAP1_NRF2*$_{closed}$, and *NRF2*$_{tot}$ in response to termination of NRF2 synthesis (by setting $k_0$=0) starting at 0 min with $k_6$ at default value. **(B)** Dynamical changes of various NRF2 species in response to stabilization of NRF2 in *KEAP1_NRF2*$_{closed}$ by setting $k_6$=1.454E-4 starting at 0 min and in response to termination of NRF2 synthesis (by setting $k_0$=0) starting at 500 min. For simulations in (A) and (B), *CLASS*$_{I\text{-}V}$ level is kept at zero. **(C-E)** Dynamical changes of *NRF2*$_{tot}$ (C), *NRF2*$_{free}$ (D), and *KEAP1*$_{free\_tot}$ (*KEAP1*$_{free}$+*KEAP1o*$_{free}$) (E) in response to different levels of *CLASS*$_{I\text{-}V}$ (ranging from 0.1 to 1E4 nM) with $k'_6$ at default value. **(F)** Steady-state dose-response curves of various NRF2 species and *KEAP1*$_{free\_tot}$ on double-log scale with $k'_6$ at default value. Shown are $n_H$ and $LRC_{max}$ for *NRF2*$_{free}$; $n_H$ and $LRC_{max}$ for *NRF2*$_{total}$ are 1.27 and 0.42 respectively (not shown). **(G-H)** Flux analyses for conditions when NRF2 in *KEAP1_NRF2*$_{closed}$ is stabilized by setting $k_6$ to 30% (G) and 10% (H) of default value.

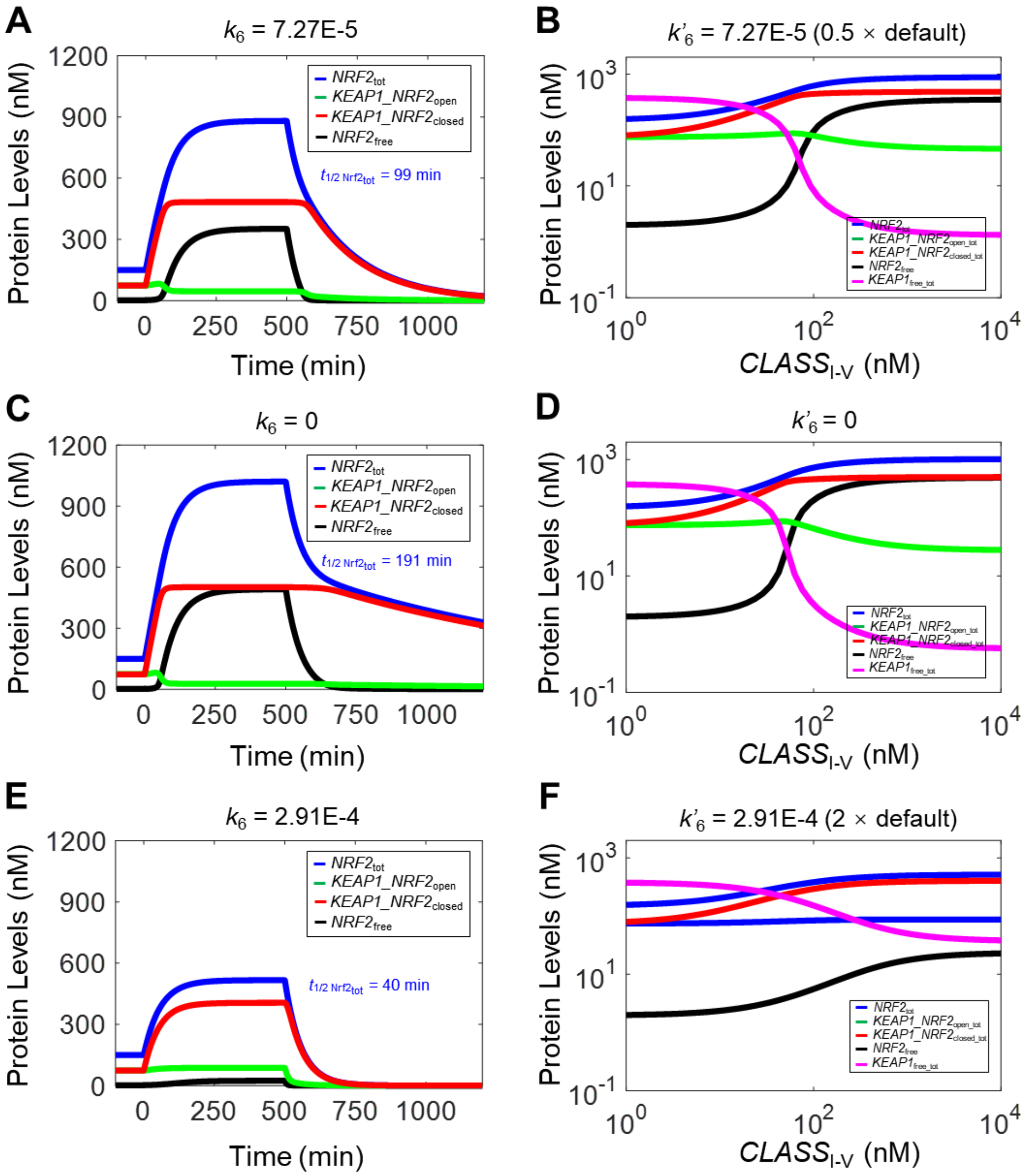

**Figure S4. Additional dynamical and steady-state behaviors of Model 2.** Dynamical changes of $NRF2_{free}$, $KEAP1\_NRF2_{open}$, $KEAP1\_NRF2_{closed}$, and $NRF2_{tot}$ in response to stabilization of NRF2 in $KEAP1\_NRF2_{closed}$ by setting **(A)** $k_6$=7.27E-5, **(C)** $k_6$=0, and **(E)** $k_6$=2.91E-4 starting at 0 min and in response to termination of NRF2 synthesis (by setting $k_0$=0) starting at 500 min. For simulations in (A, C, and E), $CLASS_{I-V}$ level is kept at zero. Steady-state dose-response curves of various NRF2 species and $KEAP1_{free}$ with **(B)** $k'_6$=7.27E-5, **(D)** $k'_6$=0, and **(F)** $k'_6$=2.91E-4.

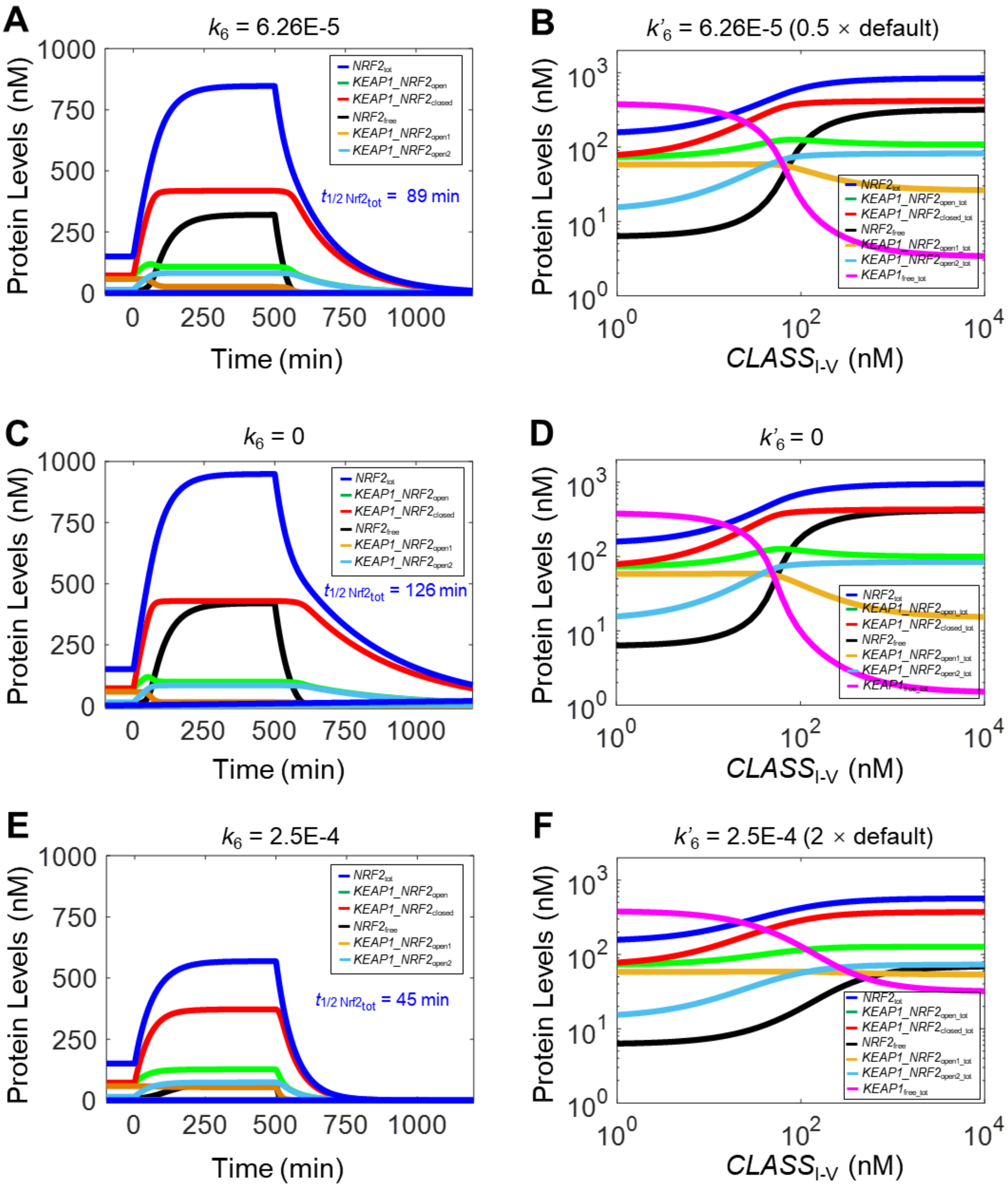

**Figure S5. Additional dynamical and steady-state behaviors of Model 3a.** Dynamical changes of $NRF2_{free}$, $KEAP1\_NRF2_{open1}$, $KEAP1\_NRF2_{open2}$, $KEAP1\_NRF2_{closed}$, and $NRF2_{tot}$ in response to stabilization of NRF2 in $KEAP1\_NRF2_{closed}$ by setting **(A)** $k_6$=6.25E-5, **(C)** $k_6$=0, and **(E)** $k_6$=2.5E-4 starting at 0 min and in response to termination of NRF2 synthesis (by setting $k_0$=0) starting at 500 min. For simulations in (A, C, and E), $CLASS_{I\text{-}V}$ level is kept at zero. Steady-state dose-response curves of various NRF2 species and $KEAP1_{free}$ with **(B)** $k'_6$=6.25E-5, **(D)** $k'_6$=0, and **(F)** $k'_6$=2.5E-4.

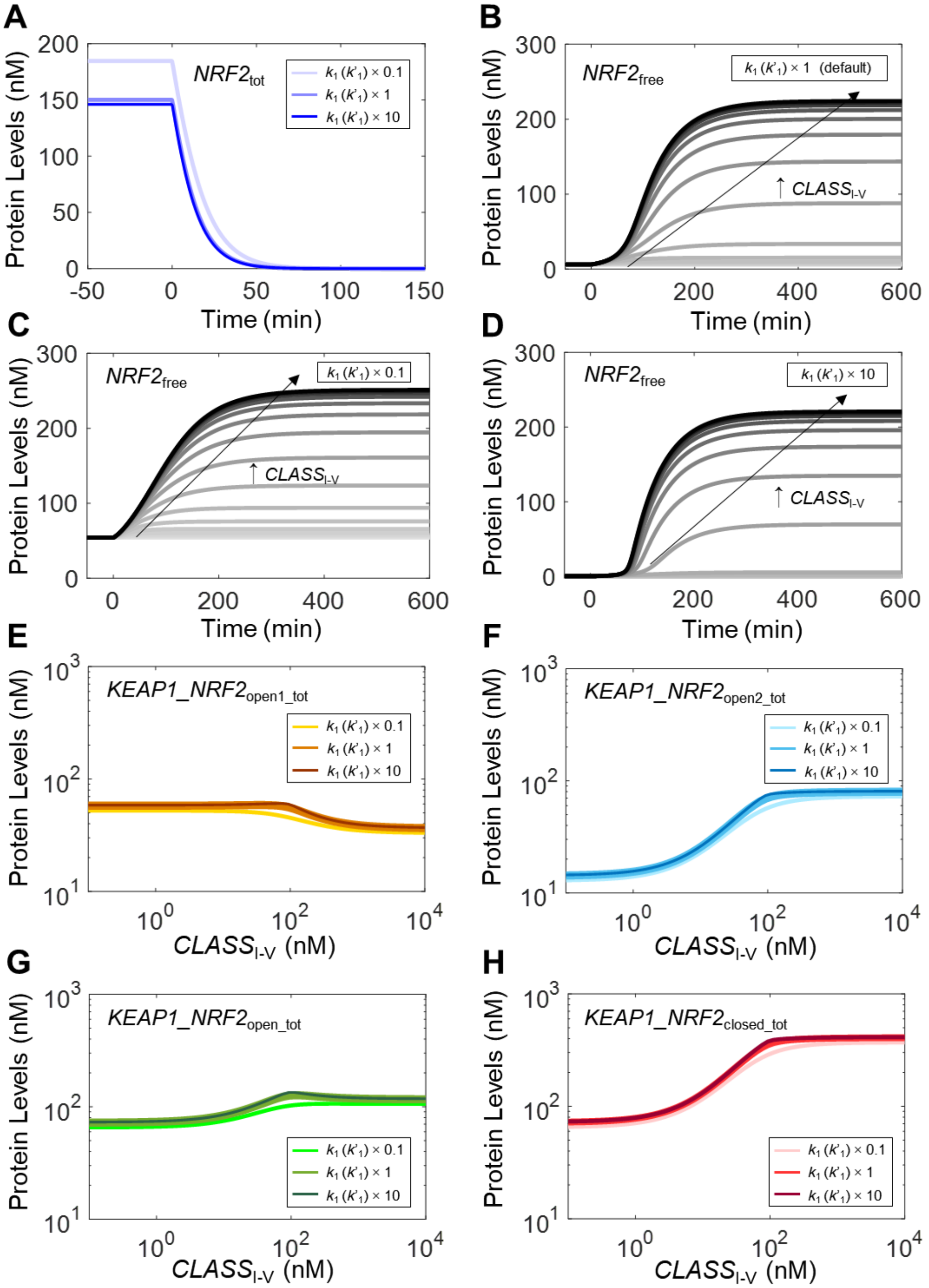

**Figure S6. Effects of parameters $k_1$ ($k'_1$) on NRF2 responses in Model 3a. (A)** Dynamical changes of basal $NRF2_{tot}$ in response to termination of NRF2 synthesis (by setting $k_0$=0) starting at 0 min with $k_1$=$k'_1$ set at different values from the default. **(B-D)** Dynamical changes of $NRF2_{free}$ in response to different $CLASS_{I\text{-}V}$ levels with $k_1$=$k'_1$ set at different indicated values from the default. **(E-H)** Steady-state dose-response curves of various NRF2 species with $k_1$=$k'_1$ set at different values from the default.

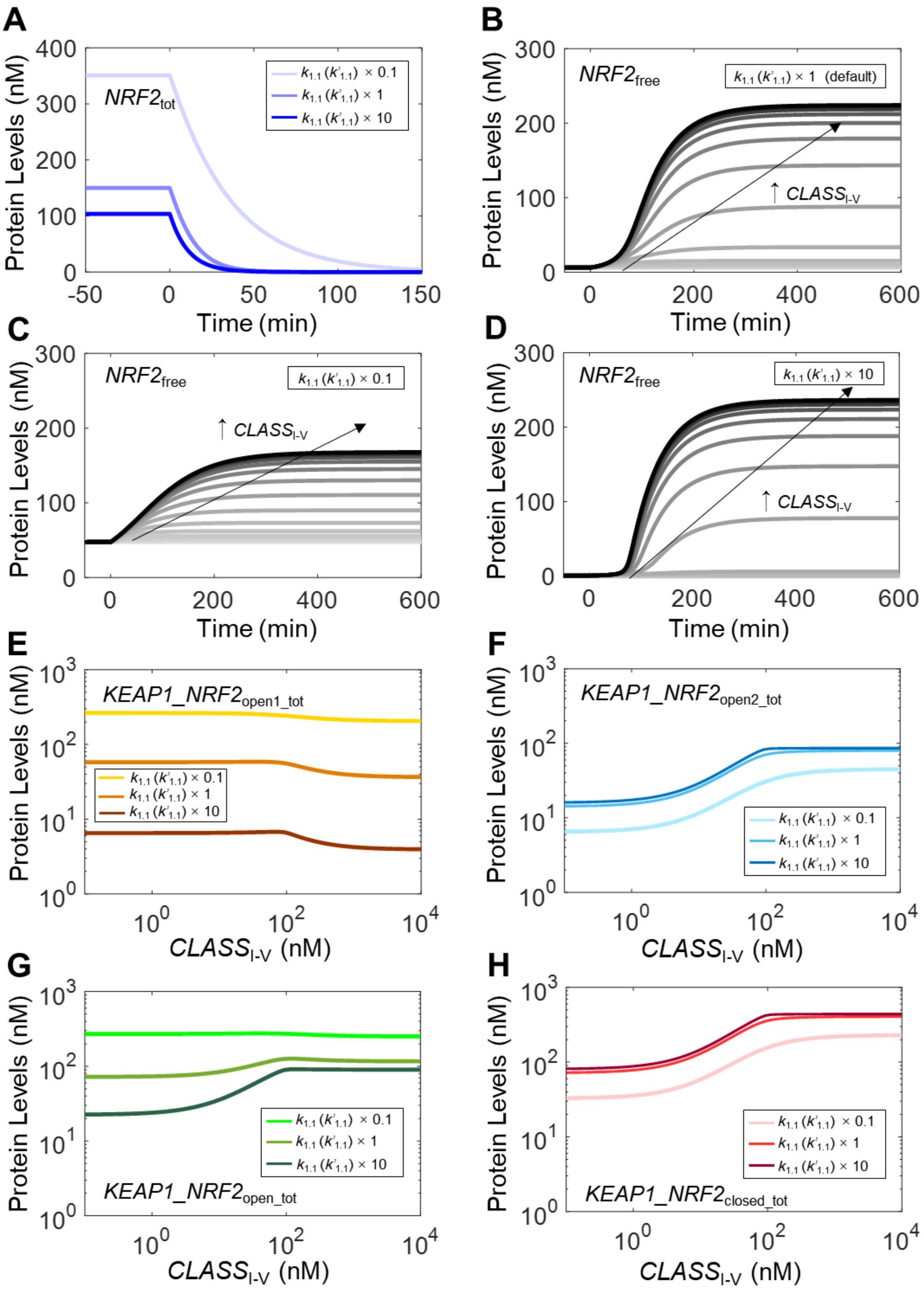
A
NRF2tot
k1.1 (k'1.1) × 0.1
k1.1 (k'1.1) × 1
k1.1 (k'1.1) × 10
Protein Levels (nM)
Time (min)
B
NRF2free
k1.1 (k'1.1) × 1 (default)
↑ CLASSI-V
Protein Levels (nM)
Time (min)
C
NRF2free
k1.1 (k'1.1) × 0.1
↑ CLASSI-V
Protein Levels (nM)
Time (min)
D
NRF2free
k1.1 (k'1.1) × 10
↑ CLASSI-V
Protein Levels (nM)
Time (min)
E
KEAP1_NRF2open1_tot
k1.1 (k'1.1) × 0.1
k1.1 (k'1.1) × 1
k1.1 (k'1.1) × 10
Protein Levels (nM)
CLASSI-V (nM)
F
KEAP1_NRF2open2_tot
k1.1 (k'1.1) × 0.1
k1.1 (k'1.1) × 1
k1.1 (k'1.1) × 10
Protein Levels (nM)
CLASSI-V (nM)
G
KEAP1_NRF2open_tot
k1.1 (k'1.1) × 0.1
k1.1 (k'1.1) × 1
k1.1 (k'1.1) × 10
Protein Levels (nM)
CLASSI-V (nM)
H
KEAP1_NRF2closed_tot
k1.1 (k'1.1) × 0.1
k1.1 (k'1.1) × 1
k1.1 (k'1.1) × 10
Protein Levels (nM)
CLASSI-V (nM)

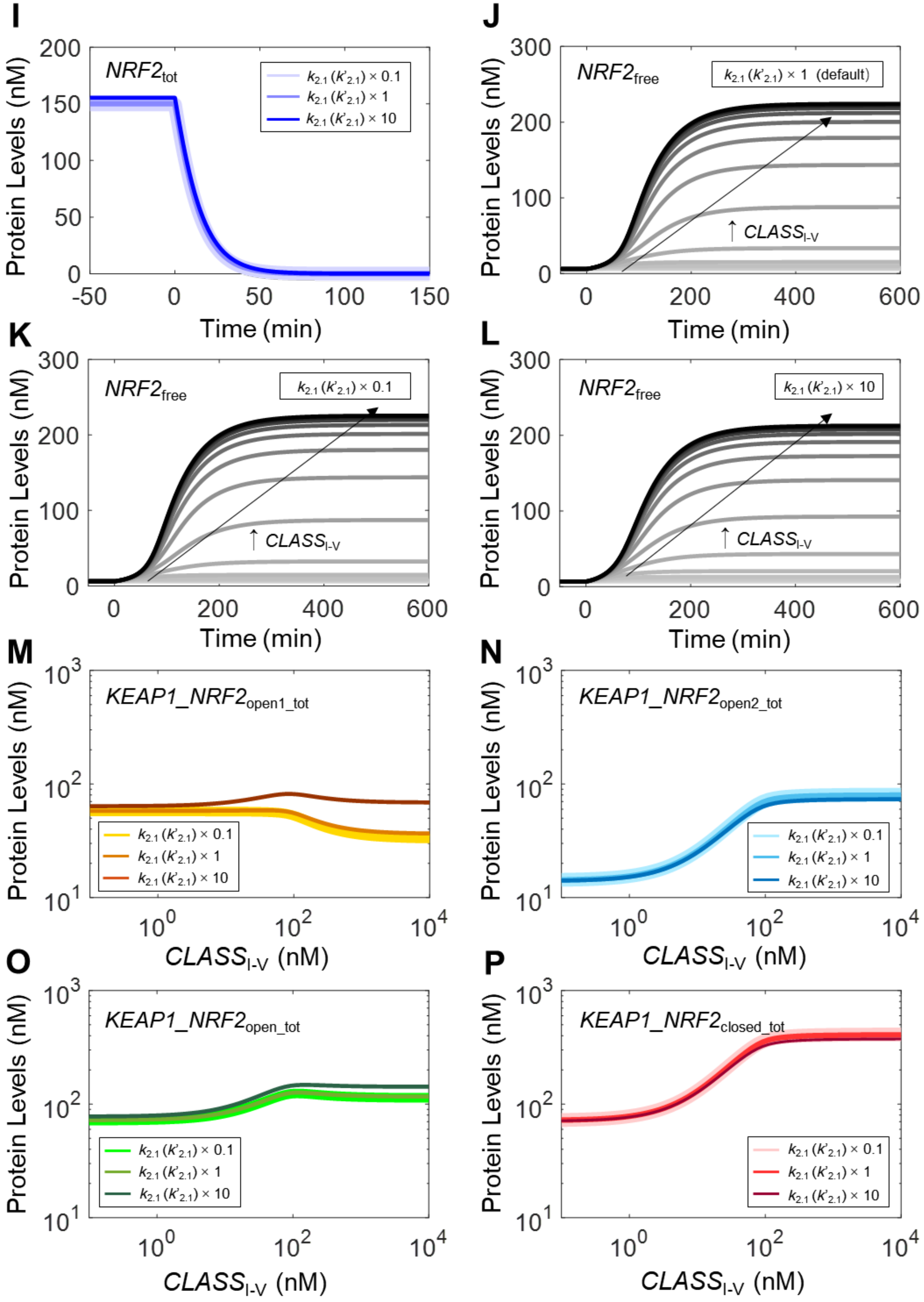
I
NRF2tot
k2.1 (k'2.1) × 0.1
k2.1 (k'2.1) × 1
k2.1 (k'2.1) × 10
Protein Levels (nM)
Time (min)
J
NRF2free
k2.1 (k'2.1) × 1 (default)
↑ CLASSI-V
Protein Levels (nM)
Time (min)
K
NRF2free
k2.1 (k'2.1) × 0.1
↑ CLASSI-V
Protein Levels (nM)
Time (min)
L
NRF2free
k2.1 (k'2.1) × 10
↑ CLASSI-V
Protein Levels (nM)
Time (min)
M
KEAP1_NRF2open1_tot
k2.1 (k'2.1) × 0.1
k2.1 (k'2.1) × 1
k2.1 (k'2.1) × 10
Protein Levels (nM)
CLASSI-V (nM)
N
KEAP1_NRF2open2_tot
k2.1 (k'2.1) × 0.1
k2.1 (k'2.1) × 1
k2.1 (k'2.1) × 10
Protein Levels (nM)
CLASSI-V (nM)
O
KEAP1_NRF2open_tot
k2.1 (k'2.1) × 0.1
k2.1 (k'2.1) × 1
k2.1 (k'2.1) × 10
Protein Levels (nM)
CLASSI-V (nM)
P
KEAP1_NRF2closed_tot
k2.1 (k'2.1) × 0.1
k2.1 (k'2.1) × 1
k2.1 (k'2.1) × 10
Protein Levels (nM)
CLASSI-V (nM)

**Figure S7. Effects of parameters $k_{1.1}$ ($k'_{1.1}$) and $k_{2.1}$ ($k'_{2.1}$) on NRF2 responses in Model 3a.** **(A)** Dynamical changes of basal $NRF2_{tot}$ in response to termination of NRF2 synthesis (by setting $k_0$=0) starting at 0 min with $k_{1.1}$=$k'_{1.1}$ set at different values from the default. **(B-D)** Dynamical changes of $NRF2_{free}$ in response to different $CLASS_{I\text{-}V}$ levels with $k_{1.1}$=$k'_{1.1}$ set at different indicated values from the default. **(E-H)** Steady-state dose-response curves of various NRF2 species with $k_{1.1}$=$k'_{1.1}$ set at different values from the default. **(I)** Dynamical changes of basal $NRF2_{tot}$ in response to termination of NRF2 synthesis (by setting $k_0$=0) starting at 0 min with $k_{2.1}$=$k'_{2.1}$ set at different values from the default. **(J-L)** Dynamical changes of $NRF2_{free}$ in response to different $CLASS_{I\text{-}V}$ levels with $k_{2.1}$=$k'_{2.1}$ set at different indicated values from the default. **(M-P)** Steady-state dose-response curves of various NRF2 species with $k_{2.1}$= $k'_{2.1}$ set at different values from the default.

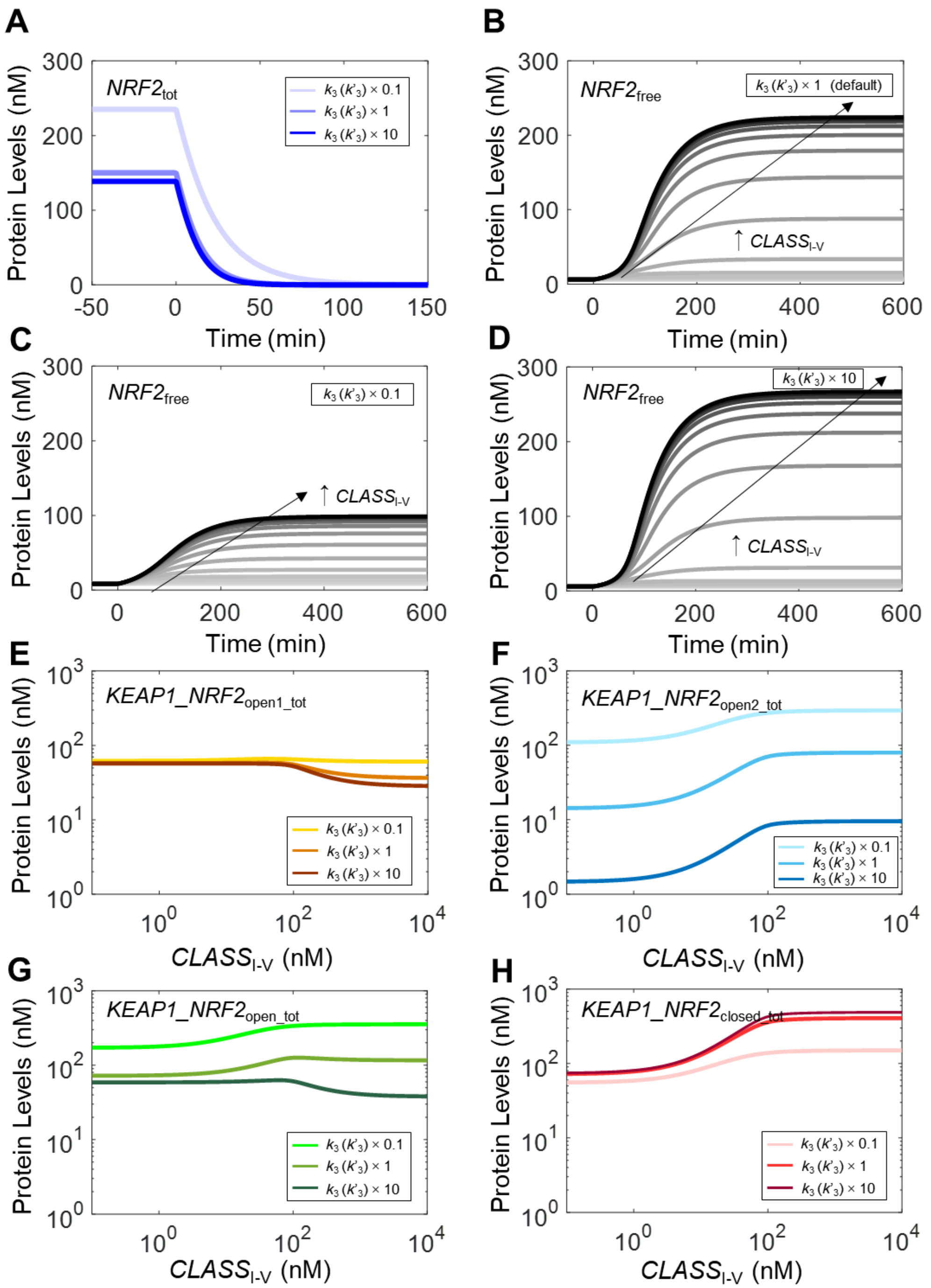

**Figure S8. Effects of parameters $k_3$ ($k'_3$) on NRF2 responses in Model 3a. (A)** Dynamical changes of basal $NRF2_{tot}$ in response to termination of NRF2 synthesis (by setting $k_0$=0) starting at 0 min with $k_3$=$k'_3$ set at different values from the default. **(B-D)** Dynamical changes of $NRF2_{free}$ in response to different $CLASS_{I\text{-}V}$ levels with $k_3$= ’$_3$ set at different indicated values from the default. **(E-H)** Steady-state dose-response curves of various NRF2 species with $k_3$=$k'_3$ set at different values from the default.

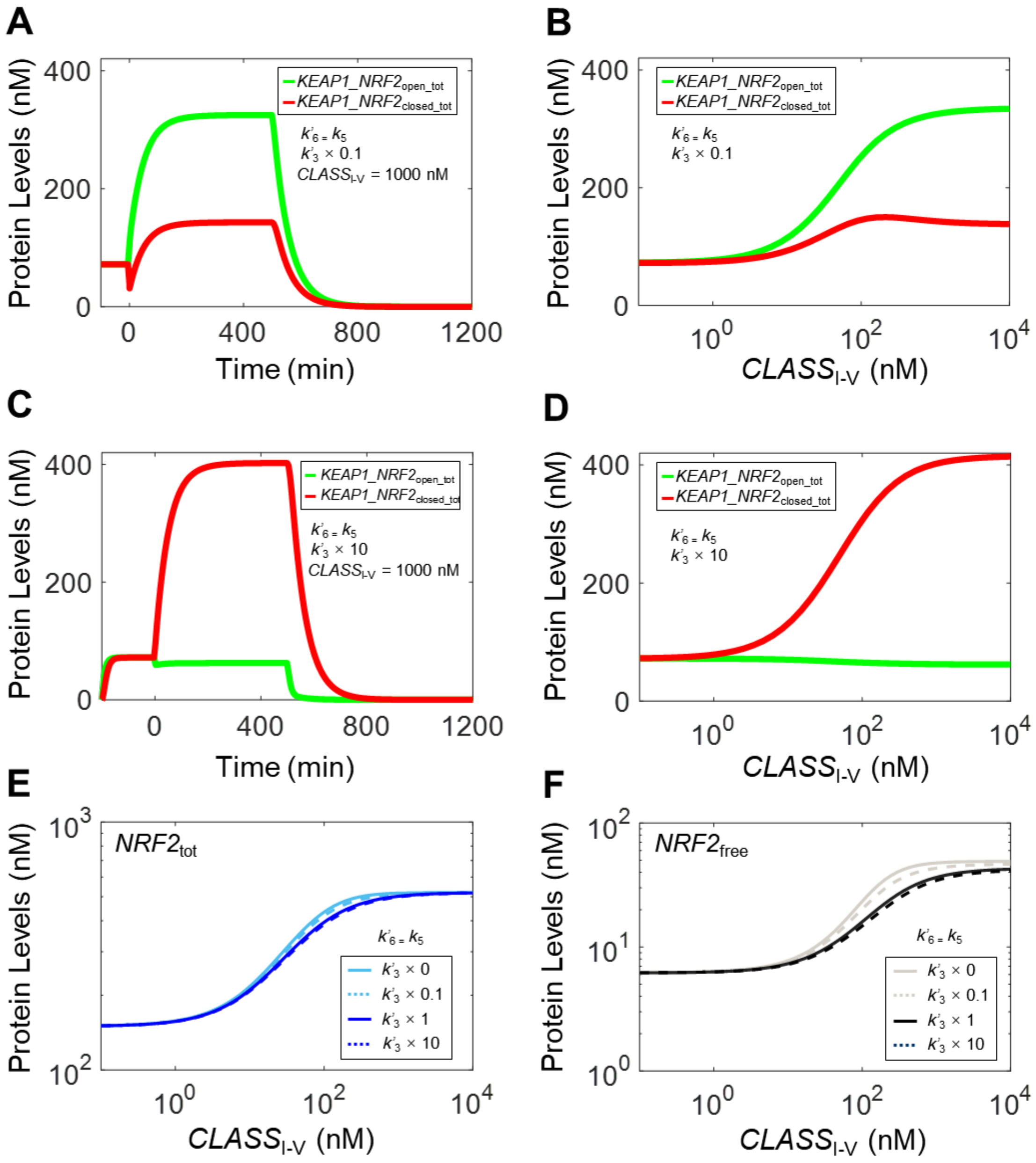

**Figure S9. Effects of varying parameter $k'_3$ alone on NRF2 responses in Model 3a to test hinge-latch hypothesis - with $k'_6$ set equal to $k_5$.** Dynamical changes of $KEAP1\_NRF2_{open\_tot}$ and $KEAP1\_NRF2_{closed\_tot}$ in response to a high level of $CLASS_{I\text{-}V}$ at 1000 nM starting at 0 min and in response to termination of NRF2 synthesis (by setting $k_0$=0) starting at 500 min, when $k'_3$ is **(A)** 0.1 and **(C)** 10 times of default value. Steady-state dose-response curves of $KEAP1\_NRF2_{open\_tot}$ and $KEAP1\_NRF2_{closed\_tot}$ when $k'_3$ is **(B)** 0.1 and **(D)** 10 times of default value. **(E-F)** Effects of varying $k'_3$ on steady-state dose-response curves of $NRF2_{tot}$ and $NRF2_{free}$ respectively.

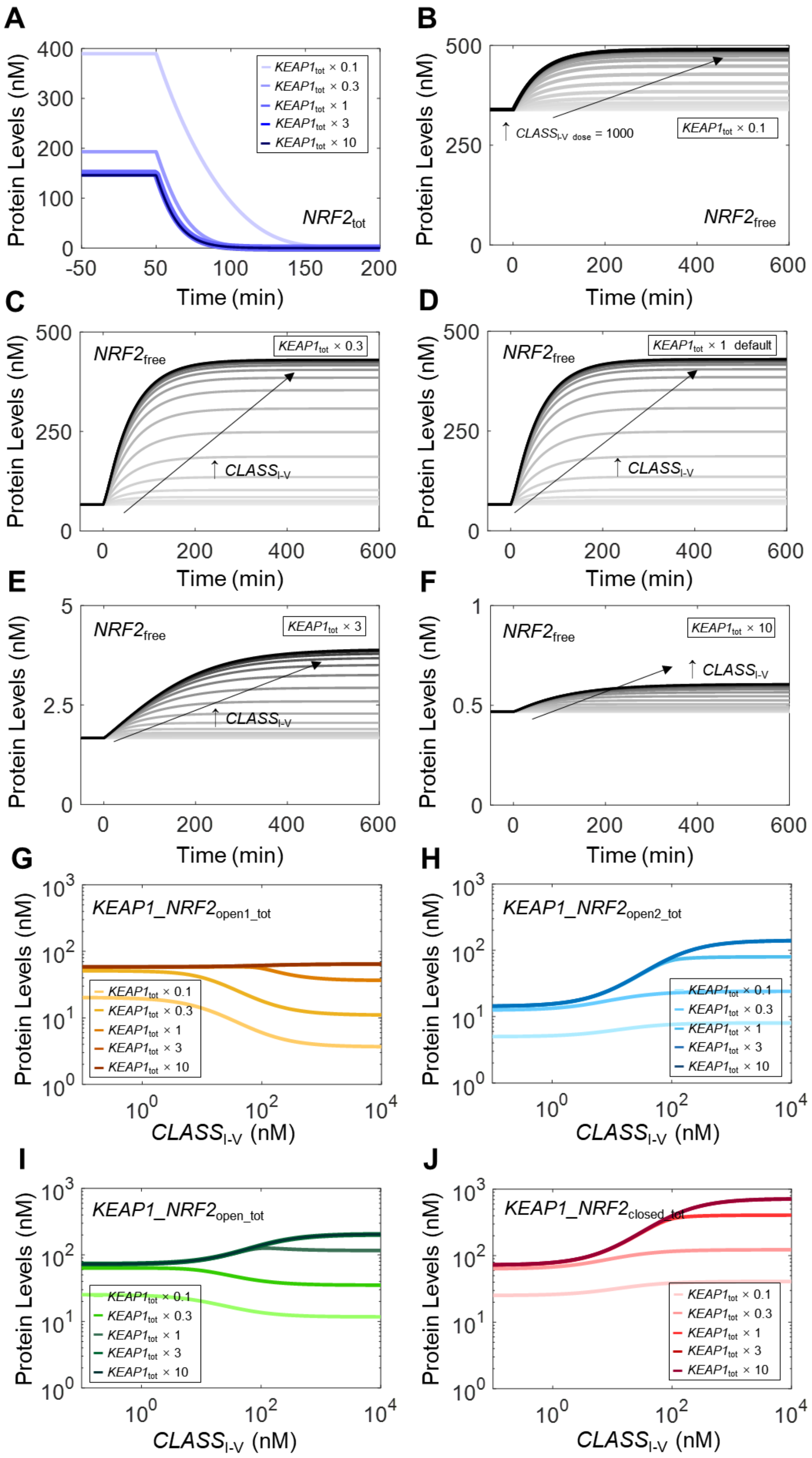
A
B
C
D
E
F
G
H
I
J
Protein Levels (nM)
Time (min)
$KEAP1_{tot}$ × 0.1
$KEAP1_{tot}$ × 0.3
$KEAP1_{tot}$ × 1
$KEAP1_{tot}$ × 3
$KEAP1_{tot}$ × 10
$NRF2_{tot}$
$NRF2_{free}$
↑ $CLASS_{I\text{-}V\ dose}$ = 1000
$KEAP1_{tot}$ × 1 default
↑ $CLASS_{I\text{-}V}$
$KEAP1\_NRF2_{open1\_tot}$
$KEAP1\_NRF2_{open2\_tot}$
$KEAP1\_NRF2_{open\_tot}$
$KEAP1\_NRF2_{closed\_tot}$
$CLASS_{I\text{-}V}$ (nM)

**Figure S10. Effects of total KEAP1 abundance on NRF2 responses in Model 3a. (A)** Dynamical changes of basal $NRF2_{tot}$ in response to termination of NRF2 synthesis (by setting $k_0$=0) starting at 0 min with $KEAP1_{tot}$ set at different values from the default. **(B-F)** Dynamical changes of $NRF2_{free}$ in response to different $CLASS_{I\text{-}V}$ levels with $KEAP1_{tot}$ set at different indicated values from the default. **(G-J)** Steady-state dose-response curves of various Keap1-NRF2 complex species with $KEAP1_{tot}$ set at different values from the default.

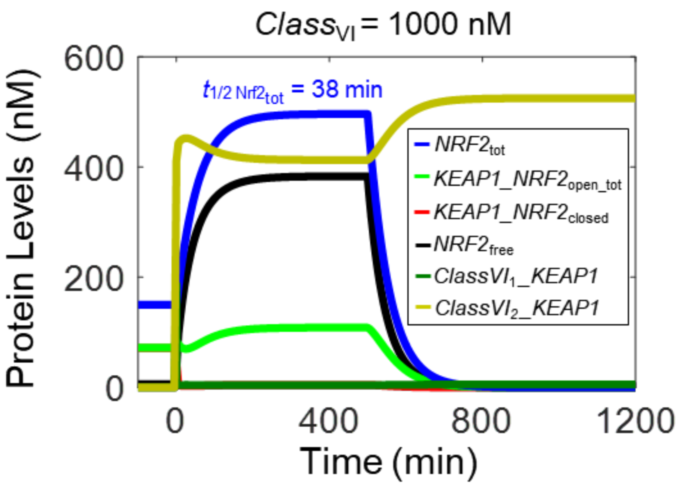

**Figure S11.** Dynamical changes of various NRF2 species in Model 3b in response to a high $CLASS_{VI}$ level (1000 nM) starting at 0 min and in response to termination of NRF2 synthesis (by setting $k_0$=0) starting at 500 min.

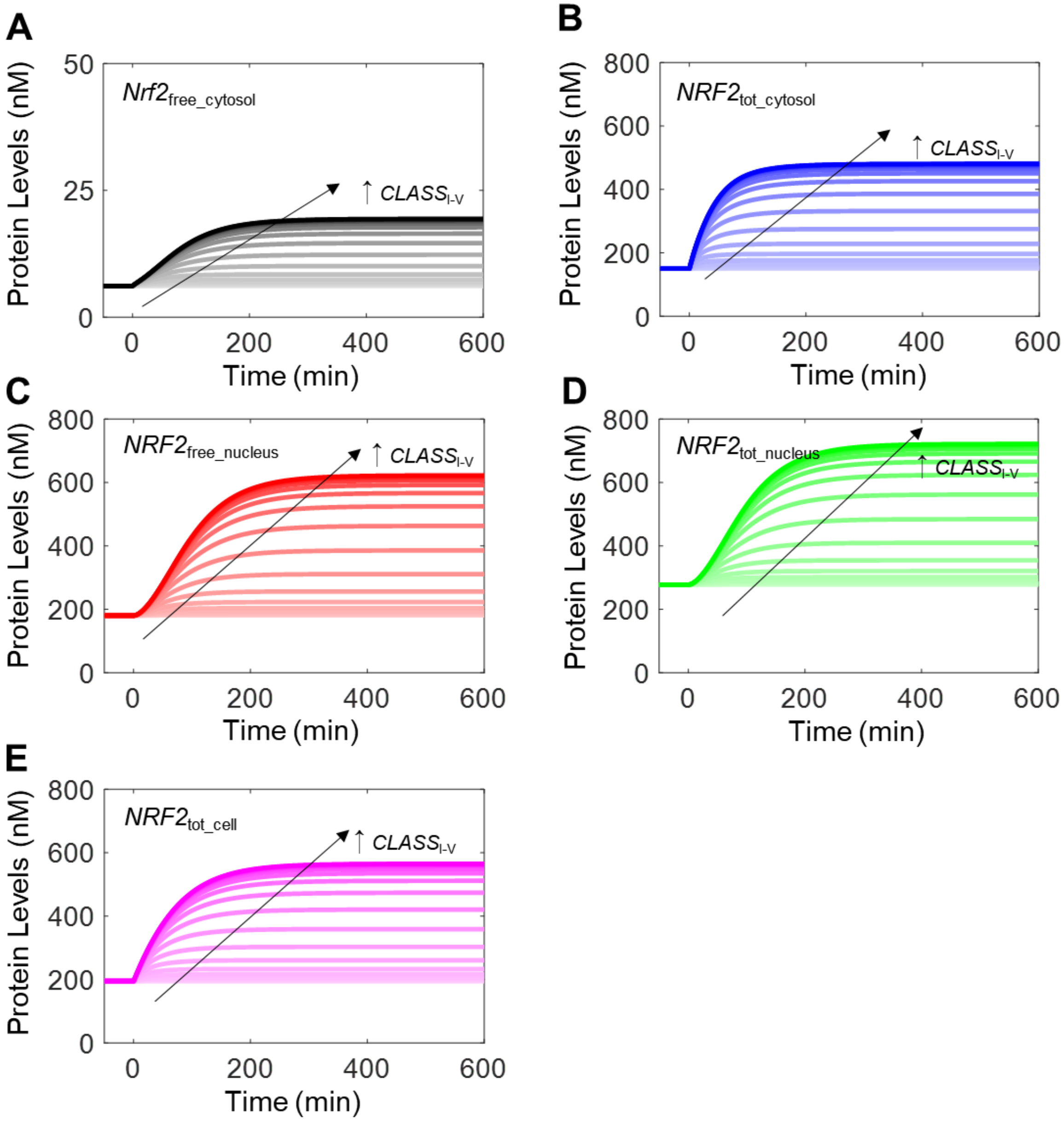

**Figure S12.** Dynamical changes of various NRF2 species in Model 4a in response to different $CLASS_{I\text{-}V}$ levels.

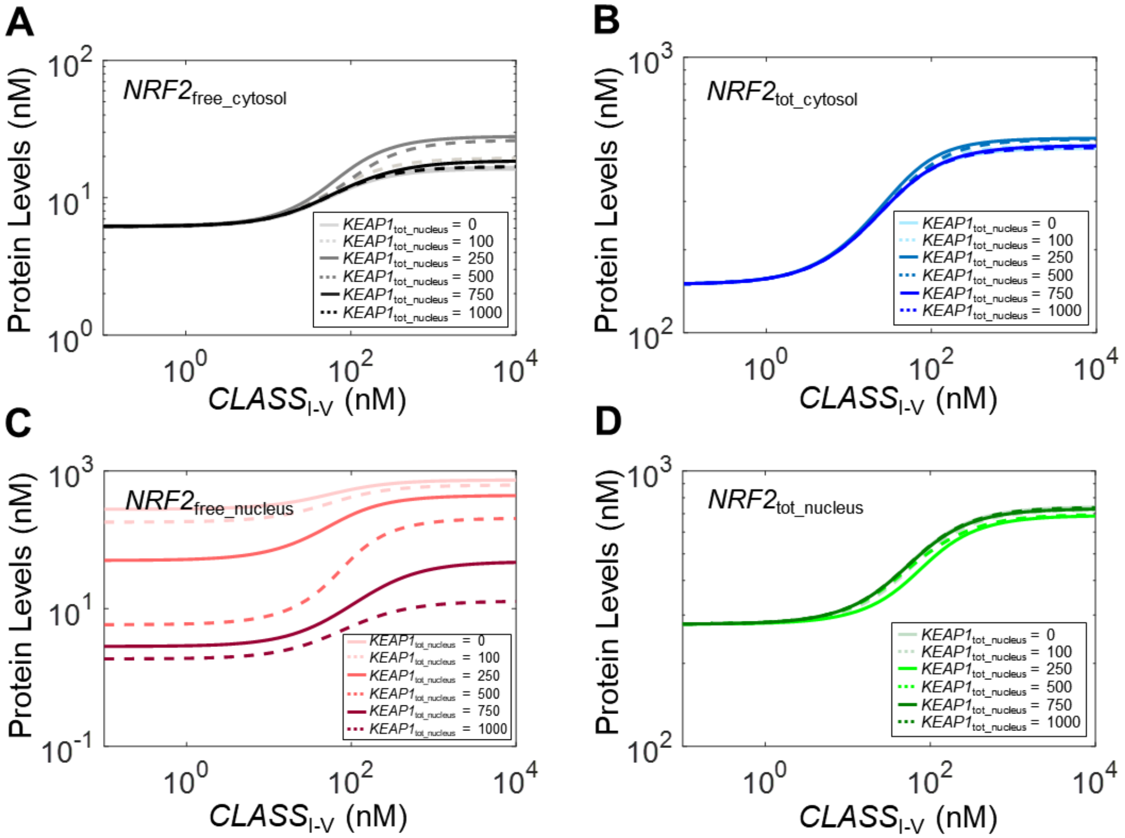

**Figure S13. Effects of total nuclear KEAP1 abundance on the steady-state dose-response curves of various NRF2 species in Model 4a.** For different total nuclear KEAP1 abundances, $k_{10}$ was adjusted accordingly to **(A)** 0.02571, **(B)** 0.01916，**(C)** 0.01039，**(D)** 7.405E-3, **(E)** 7.203E-3, and **(F)** 7.138E-3, such that the basal steady-state $NRF2_{tot\_cytosol}$ and $NRF2_{free\_cytosol}$ levels did not deviate from default values.

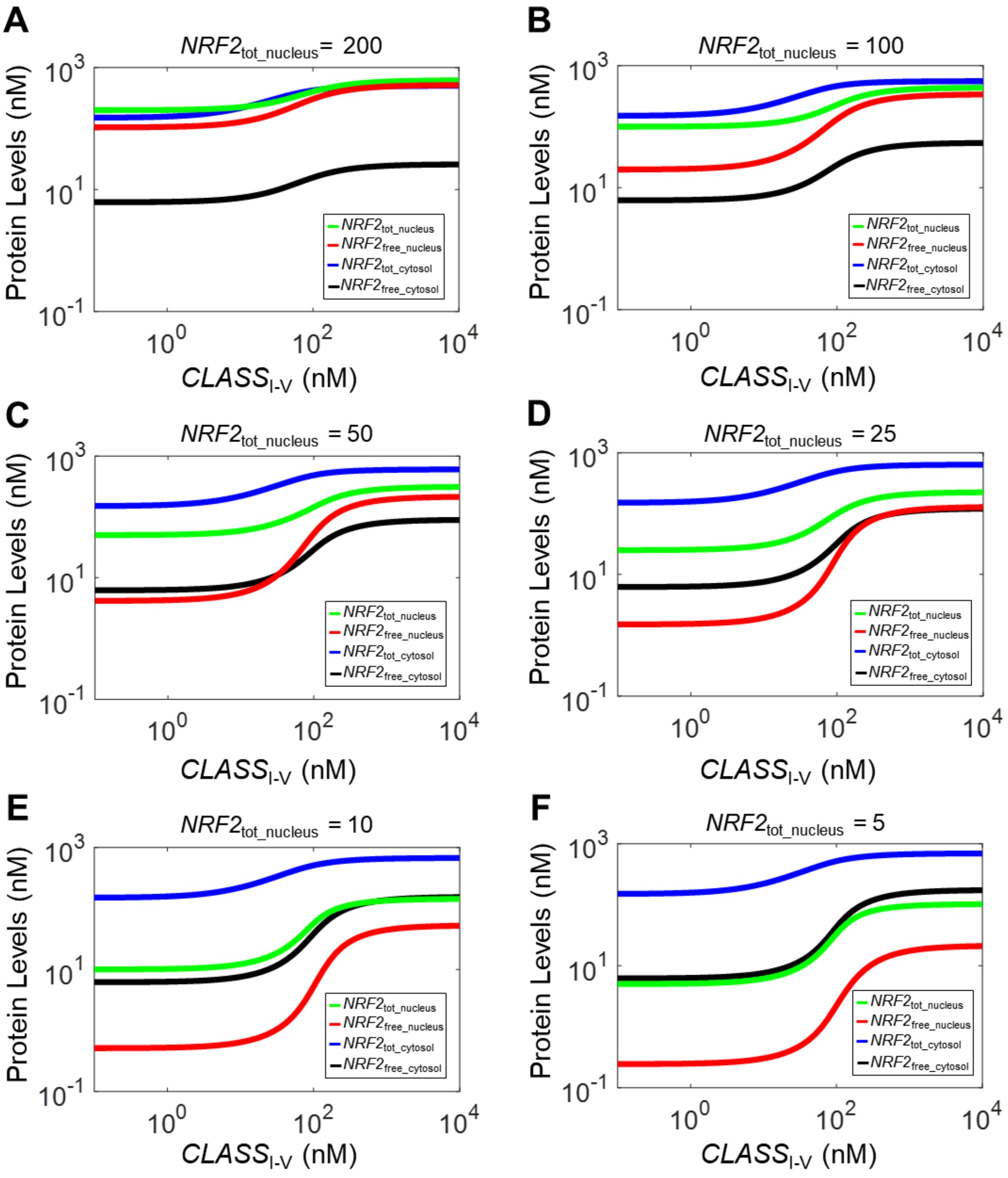

**Figure S14. Effects of basal total nuclear NRF2 abundance on steady-state dose-response of various NRF2 species in Model 4a.** Different total nuclear NRF2 abundances were achieved by setting **(A)** $k_0$=0.18125 and $k_{10}$ = 0.01206, **(B)** $k_0$=0.16556 and $k_{10}$ = 3.849E-3, **(C)** $k_0$=0.1579 and $k_{10}$ = 1.54E-3, **(D)** $k_0$=0.15398 and $k_{10}$ = 7.31E-4, **(E)** $k_0$=0.1516 and $k_{10}$ = 2.87E-4, and **(F)** $k_0$=0.1509 and $k_{10}$ = 1.425E-4. These pairs of values do not cause deviation of the basal steady-state $NRF2_{tot\_cytosol}$ and $NRF2_{free\_cytosol}$ levels from default values.

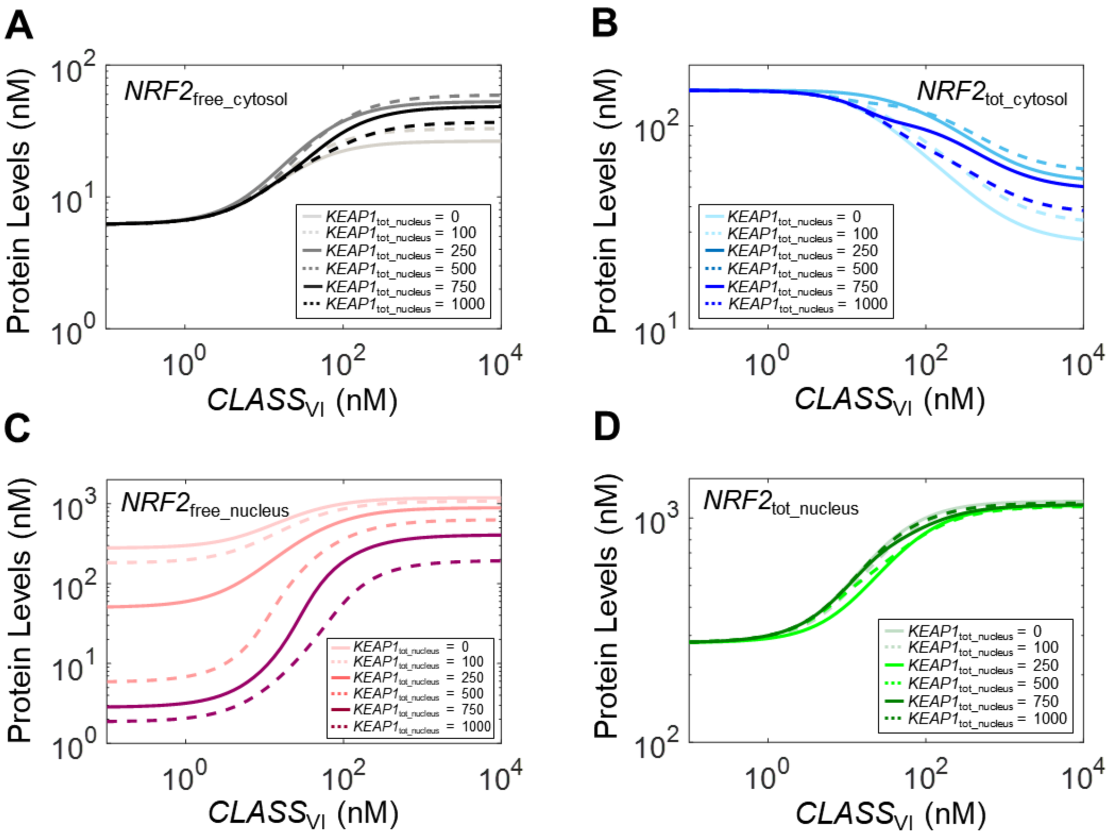

**Figure S15. Effects of total nuclear KEAP1 abundance on the steady-state dose-response curves of various NRF2 species in Model 4b.** For different total nuclear KEAP1 abundances, $k_{10}$ was adjusted accordingly to values as indicated in Figure S11 legend.

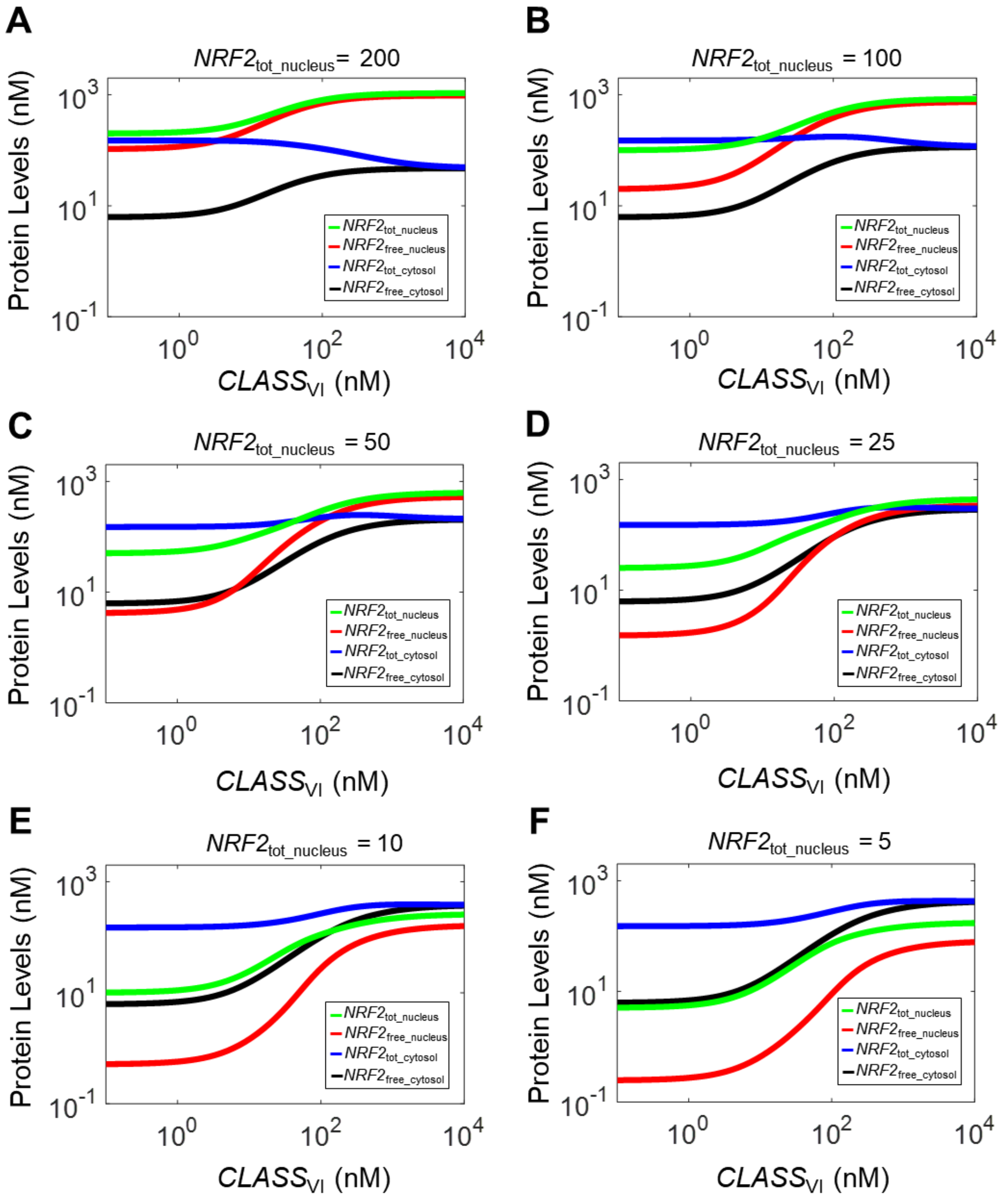

**Figure S16. Effects of basal total nuclear NRF2 abundance on steady-state dose-response of various NRF2 species in Model 4b.** Different total nuclear NRF2 abundances were achieved with the same $k_0$ and $k_{10}$ pair settings as in Figure S12 legend.

### Tables

**Table S1. Model Parameters, Default Values, Sources and Justifications**

| Parameters | Values | | | | | | Unit | Note |
|---|---|---|---|---|---|---|---|---|
| | Model 1 | Model 2 | Model 3a | Model 3b | Model 4a | Model 4b | | |
| $k_0$ | 1.721E-1 | 1.500E-1 | | | 1.933E-1 | | nM/S | i |
| $k_1$, $k'_1$ | 1.410E-2 | | 3.450E-3 | | | | 1/nM/S | ii, xvi |
| $k_2$, $k'_2$ | 2.820E-1 | | | | | | 1/S | iii, xvi |
| $k_{1.1}$, $k'_{1.1}$ | na | | 2.300E-3 | | | | 1/S | iv, xvi |
| $k_{2.1}$, $k'_{2.1}$ | na | | 1.220E-4 | | | | 1/S | v, xvi |
| $k_3$, $k'_3$ | 1.975E-1 | 1.82E-3 | 1.000 | na | 1.000 | na | 1/S | vi, xvi |
| $k_4$, $k'_4$ | 1.960E-1 | 1.000E-4 | 1.960E-1 | na | 1.960E-1 | na | 1/S | vii, xvi |
| $k_5$ | 2.888E-4 | | | | | | 1/S | viii |
| $k_6$ | 2.03E-3 | 1.74E-3 | 1.775E-3 | | | | 1/S | ix |
| $k'_6$ | 1.178E-4 | 1.454E-4 | 1.252E-4 | na | 1.252E-4 | na | 1/S | x |
| $k_7$ | 1.000E-2 | | | | | | 1/nM/S | xi |
| $k'_7$ | na | | | 1.000E-2 | na | 1.000E-2 | 1/nM/S | xi |
| $k_8$ | 1.000E-1 | | | | | | 1/S | xi |
| $k'_8$ | na | | | 1.000E-1 | na | 1.000E-1 | 1/S | xi |
| $k_9$, $k'_9$, | 2.888E-4 | | | | | | 1/S | xii, xvi |
| $k_{9.1}$, $k'_{9.1}$ | na | | 2.888E-4 | | | | 1/S | xiii, xvi |
| $k_{10}$ | na | | | | 1.916E-2 | | 1/S | xiv |
| $k_{11}$ | na | | | | 7.702E-4 | | 1/S | xv |
| $k_{n1}$ | na | | | | 3.450E-3 | | 1/nM/S | xvii |
| $k_{n2}$ | na | | | | 2.820E-1 | | 1/S | xvii |
| $k_{n1.1}$ | na | | | | 2.300E-3 | | 1/S | xvii |
| $k_{n2.1}$ | na | | | | 1.220E-4 | | 1/S | xvii |
| $k_{n3}$ | na | | | | 1.000 | | 1/S | xvii |
| $k_{n4}$ | na | | | | 1.960E-1 | | 1/S | xvii |
| $k_{n5}$, $k_{n6}$, $k_{n9}$, $k_{n9.1}$ | na | | | | 2.888E-4 | | 1/S | xvii |
| *Basal cytosolic total NRF2* | 150 | | | | | | nM | xviii |
| *Cytosolic total KEAP1 dimer* | 530 | | | | | | nM | xix |
| *Basal total nuclear NRF2* | na | | | 278 | na | 278 | nM | xx |
| *Total nuclear KEAP1 dimer* | na | | | 100 | na | 100 | nM | xxi |
| $V_n/V_c$ | na | | | 0.54 | na | 0.54 | - | xxii |

na: not applicable

#### Note:

(i) $\boldsymbol{k_0}$ – Zero-order rate constant for *de novo* NRF2 synthesis. As detailed in note *ix*, its value was adjusted iteratively with $k_6$ to ensure that the cytosolic total NRF2 is 150 nM (note *xviii*) at basal condition.

(ii) $\boldsymbol{k_1}$ – Second-order association rate constant for the binding between KEAP1 and the ETGE motif of NRF2. The binding affinities have been reported in several studies with various techniques. Using mouse KEAP1-DC domain fragment and NRF2-Neh2 domain fragment, the association constants ($K_a$) was reported to be 1.9E8 $M^{-1}$ (Tong et al. 2006). Also using mouse KEAP1-DC and NRF2-ETGE peptide, $K_a$ was reported to be 4.42E7 $M^{-1}$ (Ichimura et al. 2013) and 3.8E7 $M^{-1}$ (Fukutomi et al. 2014). The macroscopic dissociation constants ($K_d$) between KEAP1-DC and ETGE-containing Neh2ΔDLGex fragment was 7.31 nM (Fukutomi et al. 2014). The $K_d$ of the binding between a 16-mer peptide containing ETGE and the Kelch domain of KEAP1 was reported to be 20 nM (Lo et al. 2006) and 23.9 nM (Chen et al. 2011). In summary, these values are equivalent to $K_d$ of 5.26, 7.31, 20, 22.62, 23.9, and 26.32 nM. We chose to use $K_d$ = 20 nM as default for Models 1 and 2 which involve single-step binding between KEAP1 and ETGE. Therefore, $k_1$ was calculated as $k_1 = k_2/K_d$ = 2.82E-1/20 = 1.41E-2 $nM^{-1}S^{-1}$.

For Models 3a, 3b, 4a, and 4b, which involve 2-step binding between KEAP1 and ETGE, 3.450E-3 $nM^{-1}S^{-1}$ was used for $k_1$ as reported for the binding between KEAP1-DC and ETGE-containing Neh2ΔDLGex fragment (Fukutomi et al. 2014).

Given the symmetrical structural configurations of the two monomeric subunits of KEAP1 dimer (Ogura et al. 2010), it is expected that either of the two subunits can equally engage with ETGE. Therefore, the association rate governed by parameter $k_1$ is multiplied by 2, such that it is $2*k_1*NRF2_{free}*KEAP1_{free}$.

(iii) $\boldsymbol{k_2}$ – The dissociation rate constant between KEAP1 and ETGE of NRF2 was determined by using mouse KEAP1-DC fragment and KEAP1-DC and Neh2ΔDLGex for the first-step binding (Fukutomi et al. 2014).

(iv) $\boldsymbol{k_{1.1}}$ – The first-order association rate constant for the second-step binding between KEAP1 and ETGE of NRF2 was determined to be 1.2E-3 $S^{-1}$ by using mouse KEAP1-DC fragment and KEAP1-DC and Neh2ΔDLGex (Fukutomi et al. 2014). In our models, this parameter was adjusted to 2.3E-3, along with $k_3$ and $k_6$ to make sure that $KEAP1\_NRF2_{open}$ and $KEAP1\_NRF2_{closed}$ are equal at basal condition as observed in (Baird et al. 2013).

(v) $\boldsymbol{k_{2.1}}$ – The first-order dissociation rate constant for the second-step binding between KEAP1 and ETGE of NRF2 was determined by using mouse KEAP1-DC fragment and Neh2ΔDLGex (Fukutomi et al. 2014).

(vi) $\boldsymbol{k_3}$ – The first-order association rate constant between KEAP1 and DLG of NRF2. The binding affinities between KEAP1 and the DLG motif of NRF2 have been reported in a couple of studies. $K_a$ was reported to be 1.0E6 $M^{-1}$ between mouse KEAP1-DC domain fragment and NRF2-Neh2 domain fragment (Tong et al. 2006), and 1.9E6 $M^{-1}$ between KEAP1-DC and extended DLG motif peptide (DLGex) (Fukutomi et al. 2014). Using mouse KEAP1-DC domain fragment and DLG-containing Neh2(1-56) fragment of NRF2, $K_d$ was reported to be 3.2E3 nM (Fukutomi et al. 2014). In summary, these values are equivalent to $K_d$ of 526, 1000, and 3200 nM.

It is worth noting, however, that the binding between KEAP1-DC and DLG-containing Neh2 fragments in these *in vitro* studies above was measured as intermolecular event. In contrast, because of the much high binding affinity between KEAP1 and ETGE than between KEAP1 and DLG as indicated above, the DLG binding occurs almost always after

ETGE binding between full-length NRF2 and KEAP1 dimer *in vivo*. As a result, the DLG binding is predominantly an intramolecular event. Therefore, $k_3$ here is actually a first-order association rate constant as opposed to second-order association rate constant measured *in vitro*. For Models 1 and 2, $k_3$ was determined by manually adjusting its value such that at basal condition, the abundance of open and closed states of KEAP1-NRF2 complex are equal. For Models 3a-4b, $k_3$ was adjusted, along with $k_{1.1}$, such that basal open:closed ratio is close to 1:1, and when setting $k_0$=0 to observe NRF2 decay, the ratio at 15 min is close to 1:2.5 ratio as in (Baird et al. 2013).

(vii) ***k*₄** – The dissociation rate constant between KEAP1 and DLG of NRF2 was determined to be 1.960E-1 $S^{-1}$ by using mouse KEAP1-DC fragment and extended DLG motif peptide (DLGex) (Fukutomi et al. 2014). This value was used in all models except Model 2, in which the value was lowered to 1.0E-4 $S^{-1}$ to explore the cycle mode of operation for the open and closed KEAP1-NRF2 complexes.

(viii) **$k_5$** – The first-order rate constant for degradation of free cytosolic NRF2. It was reported that mouse NRF2 is degraded in a KEAP1-independent (redox-independent) manner via the Neh6 domain, with a half-life of about 40 min in COS-1 cells (McMahon et al. 2004) and HEK293T cells (Rada et al. 2011). Therefore, $k_5$ was determined as $k_5$ = ln(2)/(40*60) = 2.888E-4 $S^{-1}$.

(ix) **$k_6$** – First-order rate constant for NRF2 degradation mediated by KEAP1 in the closed state of the KEAP1-NRF2 complex in cytosol at basal condition. NRF2 half-life in whole cell at basal condition has been reported to range mostly between 6-20 min (Kwak et al. 2002, Alam et al. 2003, Itoh et al. 2003, Stewart et al. 2003, Kobayashi et al. 2004, He et al. 2006, Khalil et al. 2015, Crinelli et al. 2021). Since nuclear NRF2 is relatively more stable than cytosolic NRF2 (Itoh et al. 2003, Burroughs et al. 2018), likely mediated by Neh6 involving the GSK-3, *β*-TrCP, and Cul1 system (Chowdhry et al. 2013), as well as PML-NB (Burroughs et al. 2018), it is reasonable to expect that cytosolic NRF2 is actually degraded at a faster rate than measured in whole cell. We decided to use 10 min as the default cytosolic total NRF2 half-life at basal condition for all models, and $k_6$ was estimated as follows.

a. Since at basal condition, cytosolic KEAP1 dimer is 530 nM and cytosolic total NRF2 is 150 nM (note xviii and xix), given the high binding affinity between KEAP1 and NRF2 mediated by ETGE ($K_d$ = 20 nM, note ii), the majority of NRF2 is in a KEAP1-NRF2 complex. Therefore, the overall NRF2 half-life at basal condition for Models 1 and 2 can be initially approximated by the rate constants $k_9$ and $k_6$, which govern the degradation of NRF2 in the open and closed states of KEAP1-NRF2 complex, respectively.

b. Since the open and closed states of KEAP1-NRF2 complex are in equal abundance (Baird et al. 2013), for an overall NRF2 half-life of 10 min, ln(2)/(10*60) = 0.5*$k_9$+0.5*$k_6$, which produced an initial value $k_6$=0.002 (about 5.7-min half-life).

c. $k_3$ in Models 1 and 2 was then adjusted to make sure that open:closed ratio of KEAP1-NRF2 complex = 1:1.

d. $k_0$ was next adjusted to make total NRF2 equal to 150 nM (note xviii) at basal condition.

e. $k_0$, $k_3$ and $k_6$ were fine-tuned iteratively following steps (b)–(d) to make sure that the above 3 conditions (basal total NRF2 half-life = 10 min (by setting $k_0$=0 to observe NRF2 decay), basal total NRF2 = 150 nM, and open:closed ratio = 1:1) are met simultaneously.

f. As a result of the above adjustments, for Model 1, $k_6$=2.03E-3 $S^{-1}$ (equivalent half-life = 5.7 min); for Model 2, $k_6$=1.74E-3 $S^{-1}$ (equivalent half-life = 6.6 min).

g. For Model 3a-4b, $k_6$ was adjusted in a similar spirit, along with additional parameter $k_{1.1}$. The final $k_6$=1.775E-3 $S^{-1}$ (equivalent half-life = 6.5 min).

(x) $\boldsymbol{k'_6}$ – First-order rate constant for NRF2 degradation mediated by class I-V compound-oxidized (or conjugated) KEAP1 in the closed state of the KEAP1-NRF2 complex. This is the step where NRF2 stabilization occurs under oxidative stress. $k'_6$ was estimated to make sure that under full stress cytosolic total NRF2 can increase by 5-fold, which is a reasonable fold-increase (Iso et al. 2016).

(xi) $\boldsymbol{k_7}$ ***($\boldsymbol{k'_7}$) and*** $\boldsymbol{k_8}$ ***($\boldsymbol{k'_8}$)*** – For Models 1, 2, 3a and 4a, $k_7$ is the second-order rate constant for oxidation or conjugation of KEAP1 by class I-V compounds, and $k_8$ is the first-order rate constant for reduction or deconjugation of KEAP1.

For Models 3b and 4b, $k_7$ is the second-order association rate constant for the binding between a monomeric subunit of free KEAP1 dimer and class VI compounds, and $k_8$ is the corresponding first-order dissociation rate constant. Like the $k_1$ binding step, either of the two KEAP1 subunits in free KEAP1 dimer can equally engage with a class VI compound, thus the association rate governed by parameter $k_7$ is multiplied by 2, such that it is 2* $k_7$**ClassVI*$KEAP1_{\text{free}}$.

For Models 3b and 4b, $k'_7$ is the second-order association rate constant for the binding between the monomeric subunit of KEAP1 dimer already bound by a single class VI compound, and $k'_8$ is the corresponding first-order dissociation rate constant. We assumed that $k'_7=k_7$ and $k'_8=k_8$.

The values of all these parameters are set so that the oxidation/conjugation or binding events occur fast in the order of seconds. The model behaviors are insensitive to these parameters except that varying their values will shift the dose-response curves horizontally.

(xii) $\boldsymbol{k_9}$ – First-order rate constant for NRF2 degradation in the open state of KEAP1-NRF2 complex ($KEAP1\_NRF2_{\text{open}}$) for Models 1-2 or the intermediate open state of KEAP1-NRF2 complex ($KEAP1\_NRF2_{\text{open1}}$) for Models 3a-4b in cytosol. Since only in the closed state can KEAP1 mediates the degradation of NRF2, we assume that the degradation of NRF2 in these open states is not affected by ETGE-bound KEAP1 and is equal to the degradation rate constant of free NRF2, such that $k_9=k_5$.

(xiii) $\boldsymbol{k_{9.1}}$ – First-order rate constant for NRF2 degradation in the final open state of KEAP1-NRF2 complex ($KEAP1\_NRF2_{\text{open2}}$) for Models 3a-4b in cytosol. Since only in the closed state can KEAP1 mediates the degradation of NRF2, we assume that the degradation of NRF2 in this open state is not affected by ETGE-bound KEAP1, such that $k_{9.1}=k_5$.

(xiv) $\boldsymbol{k_{10}}$ – First-order rate constant for nuclear importation of NRF2 in Models 4a-4b. The value was adjusted along with $k_0$ such that at basal condition nuclear total NRF2 is 278 nM and cytosolic total NRF2 is 150 nM (note xviii and xx).

(xv) $\boldsymbol{k_{11}}$ – First-order rate constant for nuclear exportation of free NRF2. The value was set such that it is not rate-limiting and half of nuclear NRF2 can translocate from the nucleus to cytosol in 15 min (equivalent to average NRF2 nuclear residence time 21.6 min).

(xvi) ***k'***$_i$ where i $\in \{1, 1.1, 2, 2.1, 3, 4, 9, 9.1\}$ – Parameters for interactions between NRF2 and *KEAP1*$_o$ (i.e., KEAP1 that is oxidized/conjugated by class I-V compounds or bound by class VI compounds), and parameters for degradation of NRF2 in the open-state complex involving *KEAP1*$_o$. Since oxidative stress by class I-V compounds do not alter the binding affinity between KEAP1 and NRF2 (Eggler et al. 2005, He et al. 2006, Kobayashi et al. 2006, Horie et al. 2021), the $k'_i$ parameters for these binding events were set the same as the corresponding $k_i$ parameters at basal condition where applicable. The degradation rate constants of NRF2 in the open-state complex in these situations are also assumed the same as at basal condition.

(xvii) $\boldsymbol{k_{ni}}$ where i $\in \{1, 1.1, 2, 2.1, 3, 4, 5, 9, 9.1\}$ – Nuclear parameters corresponding to identically-numbered cytosolic parameters ($k_i$). Their values were assumed the same as the corresponding cytosolic parameters.

(xviii) $\boldsymbol{k_{n6}}$ **–** First-order rate constant for NRF2 degradation in the closed state of the KEAP1-NRF2 complex in nucleus at basal condition. No evidence exists supporting that KEAP1-mediated NRF2 ubiquitination and rapid degradation also occurs in the nucleus, therefore we assumed $k_{n6} = k_{n9}$.

(xix) ***Basal cytosolic total NRF2*** – Total cellular NRF2 abundance has been reported to vary greatly among different cell types (Khalil et al. 2015, Iso et al. 2016), however in most of these studies the absolute cytosolic and nuclear NRF2 concentrations were not measured or reported separately. We chose 150 nM as the default value in our model which is the cytosolic total NRF2 concentration reported for RAW 264.7 cells at basal condition (Iso et al. 2016).

(xx) ***Cytosolic total KEAP1 dimer*** – Like the case of NRF2, total cellular KEAP1 abundance has been reported to vary greatly among different cell types (Khalil et al. 2015, Iso et al. 2016), however information on the absolute cytosolic and nuclear KEAP1 concentrations is generally lacking. We chose 530 nM as the default value in our model which is the cytosolic total KEAP1 dimer concentration reported for RAW 264.7 cells (Iso et al. 2016). In the Results, we also varied this parameter to explore the effect of KEAP1/NRF2 abundance ratio, which varies greatly among different cell types (Khalil et al. 2015, Iso et al. 2016).

(xxi) ***Basal nuclear total NRF2*** – 278 nM was used as the default value in Models 4a and 4b, which is the nuclear total NRF2 concentration reported for RAW 264.7 cells at basal condition (Iso et al. 2016). In the Results, we also varied basal nuclear total NRF2 concentration to explore its effect of on nuclear NRF2 accumulation.

(xxii) ***Nuclear total KEAP1 dimer*** – 100 nM was used as the default value in Models 4a and 4b, which is the nuclear total KEAP1 dimer concentration reported for RAW 264.7 cells (Iso et al. 2016). In the Results, we also varied this parameter to explore the effect of nuclear total KEAP1 dimer concentration on nuclear NRF2 accumulation.

(xxiii) $\boldsymbol{V_n/V_c}$ – Ratio of nuclear volume to cytosolic volume. 0.54 was used in Models 4a and 4b, which is the average ratio for mononuclear cells during mouse embryonic development (Sasaki and Matsumura 1989).

**Table S2. ODEs of Models 1 and 2**

| | | | | |
|---|---|---|---|---|
| *1* | Free NRF2 | $\frac{dNRF2_{free}}{dt}$ | = | $k_0 - k_5 \times NRF2_{free} - 2 \times k_1 \times KEAP1_{free} \times NRF2_{free} + k_2 \times KEAP1\_NRF2_{open} - 2 \times k'_1 \times KEAP1_{o\_free} \times NRF2_{free} + k'_2 \times KEAP1_o\_NRF2_{open}$ |
| *2* | Free KEAP1 dimer | $\frac{dKEAP1_{free}}{dt}$ | = | $- 2 \times k_1 \times KEAP1_{free} \times NRF2_{free} + k_2 \times KEAP1\_NRF2_{open} - k_7 \times KEAP1_{free} \times CLASS_{I-V} + k_8 \times KEAP1_{o\_free} + k_6 \times KEAP1\_NRF2_{closed} + k_9 \times KEAP1\_NRF2_{open}$ |
| *3* | Free oxidized KEAP1 dimer | $\frac{dKEAP1_{o\_free}}{dt}$ | = | $- 2 \times k'_1 \times KEAP1_{o\_free} \times NRF2_{free} + k'_2 \times KEAP1_o\_NRF2_{open} + k_7 \times KEAP1_{free} \times CLASS_{I-V} - k_8 \times KEAP1_{o\_free} + k'_9 \times KEAP1_o\_NRF2_{open} + k'_6 \times KEAP1_o\_NRF2_{closed}$ |
| *4* | Open-state KEAP1-NRF2 complex | $\frac{dKEAP1\_NRF2_{open}}{dt}$ | = | $2 \times k_1 \times KEAP1_{free} \times NRF2_{free} - k_2 \times KEAP1\_NRF2_{open} - k_3 \times KEAP1\_NRF2_{open} + k_4 \times KEAP1\_NRF2_{closed} - k_7 \times CLASS_{I-V} \times KEAP1\_NRF2_{open} + k_8 \times KEAP1_o\_NRF2_{open} - k_9 \times KEAP1\_NRF2_{open}$ |
| *5* | Open-state KEAP1$_o$-NRF2 complex | $\frac{dKEAP1_o\_NRF2_{open}}{dt}$ | = | $2 \times k'_1 \times KEAP1_{o\_free} \times NRF2_{free} - k'_2 \times KEAP1_o\_NRF2_{open} - k'_3 \times KEAP1_o\_NRF2_{open} + k'_4 \times KEAP1_o\_NRF2_{closed} + k_7 \times CLASS_{I-V} \times KEAP1\_NRF2_{open} - k_8 \times KEAP1_o\_NRF2_{open} - k'_9 \times KEAP1_o\_NRF2_{open}$ |
| *6* | Closed-state KEAP1-NRF2 complex | $\frac{dKEAP1\_NRF2_{closed}}{dt}$ | = | $k_3 \times KEAP1\_NRF2_{open} - k_4 \times KEAP1\_NRF2_{closed} - k_6 \times KEAP1\_NRF2_{closed} - k_7 \times CLASS_{I-V} \times KEAP1\_NRF2_{closed} + k_8 \times KEAP1_o\_NRF2_{closed}$ |
| *7* | Closed-state KEAP1$_o$-NRF2 complex | $\frac{dKEAP1_o\_NRF2_{closed}}{dt}$ | = | $k'_3 \times KEAP1_o\_NRF2_{open} - k'_4 \times KEAP1_o\_NRF2_{closed} + k_7 \times CLASS_{I-V} \times KEAP1\_NRF2_{closed} - k_8 \times KEAP1_o\_NRF2_{closed} - k'_6 \times KEAP1_o\_NRF2_{closed}$ |

**Table S3. ODEs of Model 3a**

| | | | | |
|---|---|---|---|---|
| *1* | Free NRF2 | $\frac{dNRF2_{free}}{dt}$ | = | $k_0 - k_5 \times NRF2_{free} - 2 \times k_1 \times KEAP1_{free} \times NRF2_{free} + k_2 \times KEAP1\_NRF2_{open1} - 2 \times k'_1 \times KEAP1_{o\_free} \times NRF2_{free} + k'_2 \times KEAP1_o\_NRF2_{open1}$ |
| *2* | Free KEAP1 dimer | $\frac{dKEAP1_{free}}{dt}$ | = | $- 2 \times k_1 \times KEAP1_{free} \times NRF2_{free} + k_2 \times KEAP1\_NRF2_{open1} - k_7 \times KEAP1_{free} \times CLASS_{I\text{-}V} + k_8 \times KEAP1_{o\_free} + k_6 \times KEAP1\_NRF2_{closed} + k_9 \times KEAP1\_NRF2_{open1} + k_{9.1} \times KEAP1\_NRF2_{open2}$ |
| *3* | Free oxidized KEAP1 dimer | $\frac{dKEAP1_{o\_free\_cytosol}}{dt}$ | = | $- 2 \times k'_1 \times KEAP1_{o\_free} \times NRF2_{free} + k'_2 \times KEAP1_o\_NRF2_{open1} + k_7 \times KEAP1_{free} \times CLASS_{I\text{-}V} - k_8 \times KEAP1_{o\_free} + k'_6 \times KEAP1_o\_NRF2_{closed} + k'_9 \times KEAP1_o\_NRF2_{open1} + k'_{9.1} \times KEAP1_o\_NRF2_{open2}$ |
| *4* | Intermediate open-state KEAP1-NRF2 complex | $\frac{dKEAP1\_NRF2_{open1}}{dt}$ | = | $2 \times k_1 \times KEAP1_{free} \times NRF2_{free} - k_2 \times KEAP1\_NRF2_{open1} - k_{1.1} \times KEAP1\_NRF2_{open1} + k_{2.1} \times KEAP1\_NRF2_{open2} - k_7 \times CLASS_{I\text{-}V} \times KEAP1\_NRF2_{open1} + k_8 \times KEAP1_o\_NRF2_{open1} - k_9 \times KEAP1\_NRF2_{open1}$ |
| *5* | Intermediate open-state KEAP1$_o$-NRF2 complex | $\frac{dKEAP1_o\_NRF2_{open1}}{dt}$ | = | $2 \times k'_1 \times KEAP1_{o\_free} \times NRF2_{free} - k'_2 \times KEAP1_o\_NRF2_{open1} - k'_{1.1} \times KEAP1_o\_NRF2_{open1} + k'_{2.1} \times KEAP1_o\_NRF2_{open2} + k_7 \times CLASS_{I\text{-}V} \times KEAP1\_NRF2_{open1} - k_8 \times KEAP1_o\_NRF2_{open1} - k'_9 \times KEAP1_o\_NRF2_{open1}$ |
| *6* | Final open-state KEAP1-NRF2 complex | $\frac{dKEAP1\_NRF2_{open2}}{dt}$ | = | $k_{1.1} \times KEAP1\_NRF2_{open1} - k_{2.1} \times KEAP1\_NRF2_{open2} - k_3 \times KEAP1\_NRF2_{open2} + k_4 \times KEAP1\_NRF2_{closed} - k_7 \times CLASS_{I\text{-}V} \times KEAP1\_NRF2_{open2} + k_8 \times KEAP1_o\_NRF2_{open2} - k_{9.1} \times KEAP1\_NRF2_{open2}$ |
| *7* | Final open-state KEAP1$_o$-NRF2 complex | $\frac{dKEAP1_o\_NRF2_{open2}}{dt}$ | = | $k'_{1.1} \times KEAP1_o\_NRF2_{open1} - k'_{2.1} \times KEAP1_o\_NRF2_{open2} - k'_3 \times KEAP1_o\_NRF2_{open2} + k'_4 \times KEAP1_o\_NRF2_{closed} + k_7 \times CLASS_{I\text{-}V} \times KEAP1\_NRF2_{open2} - k_8 \times KEAP1_o\_NRF2_{open2} - k'_{9.1} \times KEAP1_o\_NRF2_{open2}$ |
| *8* | Closed-state KEAP1-NRF2 complex | $\frac{dKEAP1\_NRF2_{closed}}{dt}$ | = | $k_3 \times KEAP1\_NRF2_{open2} - k_4 \times KEAP1\_NRF2_{closed} - k_6 \times KEAP1\_NRF2_{closed} - k_7 \times CLASS_{I\text{-}V} \times KEAP1\_NRF2_{closed} + k_8 \times KEAP1_o\_NRF2_{closed}$ |
| *9* | Closed-state KEAP1$_o$-NRF2 complex | $\frac{dKEAP1_o\_NRF2_{closed}}{dt}$ | = | $k'_3 \times KEAP1_o\_NRF2_{open2} - k'_4 \times KEAP1_o\_NRF2_{closed} + k_7 \times CLASS_{I\text{-}V} \times KEAP1\_NRF2_{closed} - k_8 \times KEAP1_o\_NRF2_{closed} - k'_6 \times KEAP1_o\_NRF2_{closed}$ |

**Table S4. ODEs of Model 3b**

| | | | | |
|---|---|---|---|---|
| *1* | Free NRF2 | $\frac{dNRF2_{free}}{dt}$ | = | $k_0 - k5 \times NRF2_{free} - 2 \times k_1 \times KEAP1_{free} \times NRF2_{free} + k_2 \times KEAP1\_NRF2_{open1} - k'_1 \times CLASS_{VI}\_KEAP1_{free} \times NRF2_{free} + k'_2 \times CLASS_{VI}\_KEAP1\_NRF2_{open1}$ |
| *2* | Free KEAP1 dimer | $\frac{dKEAP1_{free}}{dt}$ | = | $- 2 \times k_1 \times KEAP1_{free} \times NRF2_{free} + k_2 \times KEAP1\_NRF2_{open1} - 2 \times k_7 \times KEAP1_{free} \times CLASS_{VI} + k_8 \times CLASS_{VI}\_KEAP1 + k_6 \times KEAP1\_NRF2_{closed} + k_9 \times KEAP1\_NRF2_{open1} + k_{9.1} \times KEAP1\_NRF2_{open2}$ |
| *3* | Single ClassVI-bound KEAP1 dimer | $\frac{dCLASS_{VI}\_KEAP1}{dt}$ | = | $- k'_1 \times CLASS_{VI}\_KEAP1 \times NRF2_{free} + k'_2 \times ClassVI\_KEAP1\_NRF2_{open1} + 2 \times k_7 \times KEAP1_{free} \times CLASS_{VI} - k_8 \times CLASS_{VI}\_KEAP1 - k'_7 \times CLASS_{VI} \times CLASS_{VI}\_KEAP1 + k'_8 \times CLASS_{VI2}\_KEAP1 + k'_9 \times CLASS_{VI}\_KEAP1\_NRF2_{open1} + k'_{9.1} \times CLASS_{VI}\_KEAP1\_NRF2_{open2}$ |
| *4* | Dual ClassVI-bound KEAP1 dimer | $\frac{dCLASS_{VI2}\_KEAP1}{dt}$ | = | $k'_7 \times CLASS_{VI} \times CLASS_{VI}\_KEAP1 - k'_8 \times CLASS_{VI2}\_KEAP1$ |
| *5* | Intermediate open-state KEAP1-NRF2 complex | $\frac{dKEAP1\_NRF2_{open1}}{dt}$ | = | $2 \times k_1 \times KEAP1_{free} \times NRF2_{free} - k_2 \times KEAP1\_NRF2_{open1} - k_{1.1} \times KEAP1\_NRF2_{open1} + k_{2.1} \times KEAP1\_NRF2_{open2} - k_7 \times CLASS_{VI} \times KEAP1\_NRF2_{open1} + k_8 \times CLASS_{VI}\_KEAP1\_NRF2_{open1} - k_9 \times KEAP1\_NRF2_{open1}$ |
| *6* | Final open-state KEAP1-NRF2 complex | $\frac{dKEAP1\_NRF2_{open2}}{dt}$ | = | $k_{1.1} \times KEAP1\_NRF2_{open1} - k_{2.1} \times KEAP1\_NRF2_{open2} - k_3 \times KEAP1\_NRF2_{open2} + k_4 \times KEAP1\_NRF2_{closed} - k_7 \times CLASS_{VI} \times KEAP1\_NRF2_{open2} + k_8 \times CLASS_{VI}\_KEAP1\_NRF2_{open2} - k_{9.1} \times KEAP1\_NRF2_{open2}$ |
| *7* | Closed-state KEAP1-NRF2 complex | $\frac{dKEAP1\_NRF2_{closed}}{dt}$ | = | $k_3 \times KEAP1\_NRF2_{open2} - k_4 \times KEAP1\_NRF2_{closed} - k_6 \times KEAP1\_NRF2_{closed}$ |
| *8* | Intermediate open-state ClassVI-KEAP1-NRF2 complex | $\frac{dCLASS_{VI}\_KEAP1\_NRF2_{open1}}{dt}$ | = | $k'_1 \times CLASS_{VI}\_KEAP1 \times NRF2_{free} - k'_2 \times CLASS_{VI}\_KEAP1\_NRF2_{open1} - k'_{1.1} \times CLASS_{VI}\_KEAP1\_NRF2_{open1} + k'_{2.1} \times CLASS_{VI}\_KEAP1\_NRF2_{open2} + k_7 \times CLASS_{VI} \times KEAP1\_NRF2_{open1} - k_8 \times CLASS_{VI}\_KEAP1\_NRF2_{open1} - k'_9 \times CLASS_{VI}\_KEAP1\_NRF2_{open1}$ |
| *9* | Final open-state ClassVI-KEAP1-NRF2 complex | $\frac{dCLASS_{VI}\_KEAP1\_NRF2_{open2}}{dt}$ | = | $k'_{1.1} \times CLASS_{VI}\_KEAP1\_NRF2_{open1} - k'_{2.1} \times CLASS_{VI}\_KEAP1\_NRF2_{open2} + k_7 \times CLASS_{VI} \times KEAP1\_NRF2_{open2} - k_8 \times CLASS_{VI}\_KEAP1\_NRF2_{open2} - k'_{9.1} \times CLASS_{VI}\_KEAP1\_NRF2_{open2}$ |

**Table S5. ODEs of Model 4a**

| | | | | |
|---|---|---|---|---|
| *1* | Cytosolic free NRF2 | $\frac{dNRF2_{free\_cytosol}}{dt}$ | = | $k_0 - k_5 \times NRF2_{free\_cytosol} - 2 \times k_1 \times KEAP1_{free\_cytosol} \times NRF2_{free\_cytosol} + k_2 \times KEAP1\_NRF2_{open1\_cytosol} - 2 \times k'_1 \times KEAP1_{o\_free\_cytosol} \times NRF2_{free\_cytosol} + k'_2 \times KEAP1_o\_NRF2_{open1\_cytosol} - k_{10} \times NRF2_{free\_cytosol} + k_{11} \times NRF2_{free\_nucleus} \times \frac{V_n}{V_c}$ |
| *2* | Cytosolic free KEAP1 dimer | $\frac{dKEAP1_{free\_cytosol}}{dt}$ | = | $- 2 \times k_1 \times KEAP1_{free\_cytosol} \times NRF2_{free\_cytosol} + k_2 \times KEAP1\_NRF2_{open1\_cytosol} - k_7 \times KEAP1_{free\_cytosol} \times CLASS_{I\text{-}V} + k_8 \times KEAP1_{o\_free\_cytosol} + k_6 \times KEAP1\_NRF2_{closed\_cytosol} + k_9 \times KEAP1\_NRF2_{open1\_cytosol} + k_{9.1} \times KEAP1\_NRF2_{open2\_cytosol}$ |
| *3* | Cytosolic free oxidized KEAP1 dimer | $\frac{dKEAP1_{o\_free\_cytosol}}{dt}$ | = | $- 2 \times k'_1 \times KEAP1_{o\_free\_cytosol} \times NRF2_{free\_cytosol} + k'_2 \times KEAP1_o\_NRF2_{open1\_cytosol} + k_7 \times KEAP1_{free\_cytosol} \times CLASS_{I\text{-}V} - k_8 \times KEAP1_{o\_free\_cytosol} + k'_6 \times KEAP1_o\_NRF2_{closed\_cytosol} + k'_9 \times KEAP1_o\_NRF2_{open1\_cytosol} + k'_{9.1} \times KEAP1_o\_NRF2_{open2\_cytosol}$ |
| *4* | Cytosolic intermediate open-state KEAP1-NRF2 complex | $\frac{dKEAP1\_NRF2_{open1\_cytosol}}{dt}$ | = | $2 \times k_1 \times KEAP1_{free\_cytosol} \times NRF2_{free\_cytosol} - k_2 \times KEAP1\_NRF2_{open1\_cytosol} - k_{1.1} \times KEAP1\_NRF2_{open1\_cytosol} + k_{2.1} \times KEAP1\_NRF2_{open2\_cytosol} - k_7 \times CLASS_{I\text{-}V} \times KEAP1\_NRF2_{open1\_cytosol} + k_8 \times KEAP1_o\_NRF2_{open1\_cytosol} - k_9 \times KEAP1\_NRF2_{open1\_cytosol}$ |
| *5* | Cytosolic intermediate open-state KEAP1$_o$-NRF2 complex | $\frac{dKEAP1_o\_NRF2_{open1\_cytosol}}{dt}$ | = | $2 \times k'_1 \times KEAP1_{o\_free\_cytosol} \times NRF2_{free\_cytosol} - k'_2 \times KEAP1_o\_NRF2_{open1\_cytosol} - k'_{1.1} \times KEAP1_o\_NRF2_{open1\_cytosol} + k'_{2.1} \times KEAP1_o\_NRF2_{open2\_cytosol} + k_7 \times CLASS_{I\text{-}V} \times KEAP1\_NRF2_{open1\_cytosol} - k_8 \times KEAP1_o\_NRF2_{open1\_cytosol} - k'_9 \times KEAP1_o\_NRF2_{open1\_cytosol}$ |
| *6* | Cytosolic final open-state KEAP1-NRF2 complex | $\frac{dKEAP1\_NRF2_{open2\_cytosol}}{dt}$ | = | $k_{1.1} \times KEAP1\_NRF2_{open1\_cytosol} - k_{2.1} \times KEAP1\_NRF2_{open2\_cytosol} - k_3 \times KEAP1\_NRF2_{open2\_cytosol} + k_4 \times KEAP1\_NRF2_{closed\_cytosol} - k_7 \times CLASS_{I\text{-}V} \times KEAP1\_NRF2_{open2\_cytosol} + k_8 \times KEAP1_o\_NRF2_{open2\_cytosol} - k_{9.1} \times KEAP1\_NRF2_{open2\_cytosol}$ |
| *7* | Cytosolic final open-state KEAP1$_o$-NRF2 complex | $\frac{dKEAP1_o\_NRF2_{open2\_cytosol}}{dt}$ | = | $k'_{1.1} \times KEAP1_o\_NRF2_{open1\_cytosol} - k'_{2.1} \times KEAP1_o\_NRF2_{open2\_cytosol} - k'_3 \times KEAP1_o\_NRF2_{open2\_cytosol} + k'_4 \times KEAP1_o\_NRF2_{closed\_cytosol} + k_7 \times CLASS_{I\text{-}V} \times KEAP1\_NRF2_{open2\_cytosol} - k_8 \times KEAP1_o\_NRF2_{open2\_cytosol} - k'_{9.1} \times KEAP1_o\_NRF2_{open2\_cytosol}$ |
| *8* | Cytosolic closed-state KEAP1-NRF2 complex | $\frac{dKEAP1\_NRF2_{closed\_cytosol}}{dt}$ | = | $k_3 \times KEAP1\_NRF2_{open2\_cytosol} - k_4 \times KEAP1\_NRF2_{closed\_cytosol} - k_6 \times KEAP1\_NRF2_{closed\_cytosol} - k_7 \times CLASS_{I\text{-}V} \times KEAP1\_NRF2_{closed\_cytosol} + k_8 \times KEAP1_o\_NRF2_{closed\_cytosol}$ |
| *9* | Cytosolic closed-state KEAP1$_o$-NRF2 complex | $\frac{dKEAP1_o\_NRF2_{closed\_cytosol}}{dt}$ | = | $k'_3 \times KEAP1_o\_NRF2_{open2\_cytosol} - k'_4 \times KEAP1_o\_NRF2_{closed\_cytosol} + k_7 \times CLASS_{I\text{-}V} \times KEAP1\_NRF2_{closed\_cytosol} - k_8 \times KEAP1_o\_NRF2_{closed\_cytosol} - k'_6 \times KEAP1_o\_NRF2_{closed\_cytosol}$ |
| *10* | Nuclear free NRF2 | $\frac{dNRF2_{free\_nucleus}}{dt}$ | = | $k_{10} \times NRF2_{free\_cytosol} \times \frac{V_c}{V_n} - k_{11} \times NRF2_{free\_nucleus} - k_{n5} \times NRF2_{free\_nucleus} - 2 \times k_{n1} \times NRF2_{free\_nucleus} \times KEAP1_{free\_nucleus} + k_{n2} \times KEAP1\_NRF2_{open1\_nucleus}$ |
| *11* | Nuclear free KEAP1 | $\frac{dKEAP1_{free\_nucleus}}{dt}$ | = | $- 2 \times k_{n1} \times NRF2_{free\_nucleus} \times KEAP1_{free\_nucleus} + k_{n2} \times KEAP1\_NRF2_{open1\_nucleus} + k_{n6} \times KEAP1\_NRF2_{closed\_nucleus} + k_{n9} \times KEAP1\_NRF2_{open1\_nucleus} + k_{n9.1} \times KEAP1\_NRF2_{open2\_nucleus}$ |

| | | | | |
|---|---|---|---|---|
| *12* | Nuclear intermediate open-state KEAP1-NRF2 complex | $\frac{dKEAP1\_NRF2_{open1\_nucleus}}{dt}$ | = | $2 \times k_{n1} \times NRF2_{free\_nucleus} \times KEAP1_{free\_nucleus} - k_{n2} \times KEAP1\_NRF2_{open1\_nucleus} - k_{n1.1} \times KEAP1\_NRF2_{open1\_nucleus} + k_{n2.1} \times KEAP1\_NRF2_{open2\_nucleus} - k_{n9} \times KEAP1\_NRF2_{open1\_nucleus}$ |
| *13* | Nuclear final open-state KEAP1-NRF2 complex | $\frac{dKEAP1\_NRF2_{open2\_nucleus}}{dt}$ | = | $k_{n1.1} \times KEAP1\_NRF2_{open1\_nucleus} - k_{n2.1} \times KEAP1\_NRF2_{open2\_nucleus} - k_{n3} \times KEAP1\_NRF2_{open2\_nucleus} + k_{n4} \times KEAP1\_NRF2_{closed\_nucleus} - k_{n9.1} \times KEAP1\_NRF2_{open2\_nucleus}$ |
| *14* | Nuclear closed-state KEAP1-NRF2 complex | $\frac{dKEAP1\_NRF2_{closed\_nucleus}}{dt}$ | = | $k_{n3} \times KEAP1\_NRF2_{open2\_nucleus} - k_{n4} \times KEAP1\_NRF2_{closed\_nucleus} - k_{n6} \times KEAP1\_NRF2_{closed\_nucleus}$ |

**Table S6. ODEs of Model 4b**

| | | |
|---|---|---|
| *1* | Cytosolic free NRF2 | $\frac{dNRF2_{free\_cytosol}}{dt} = k_0 - k5 \times NRF2_{free\_cytosol} - 2 \times k_1 \times KEAP1_{free\_cytosol} \times NRF2_{free\_cytosol} + k_2 \times KEAP1\_NRF2_{open1\_cytosol} - k'_1 \times CLASS_{VI}\_KEAP1_{free\_cytosol} \times NRF2_{free\_cytosol} + k'_2 \times ClassVI\_KEAP1\_NRF2_{open1\_cytosol} - k_{10} \times NRF2_{free\_cytosol} + k_{11} \times NRF2_{free\_nucleus} \times \frac{V_n}{V_c}$ |
| *2* | Cytosolic free KEAP1 dimer | $\frac{dKEAP1_{free\_cytosol}}{dt} = - 2 \times k_1 \times KEAP1_{free\_cytosol} \times NRF2_{free\_cytosol} + k_2 \times KEAP1\_NRF2_{open1\_cytosol} - 2 \times k_7 \times KEAP1_{free\_cytosol} \times CLASS_{VI} + k_8 \times CLASS_{VI}\_KEAP1_{\_cytosol} + k_6 \times KEAP1\_NRF2_{closed\_cytosol} + k_9 \times KEAP1\_NRF2_{open1\_cytosol} + k_{9.1} \times KEAP1\_NRF2_{open2\_cytosol}$ |
| *3* | Cytosolic single ClassVI-bound KEAP1 dimer | $\frac{dCLASS_{VI}\_KEAP1_{cytosol}}{dt} = - k'_1 \times CLASS_{VI}\_KEAP1_{\_cytosol} \times NRF2_{free\_cytosol} + k'_2 \times CLASS_{VI}\_KEAP1\_NRF2_{open1\_cytosol} + 2 \times k_7 \times KEAP1_{free\_cytosol} \times CLASS_{VI} - k_8 \times CLASS_{VI}\_KEAP1_{\_cytosol} - k'_7 \times CLASS_{VI} \times CLASS_{VI}\_KEAP1_{\_cytosol} + k'_8 \times CLASS_{VI2}\_KEAP1_{\_cytosol} + k'_9 \times CLASS_{VI}\_KEAP1\_NRF2_{open1\_cytosol} + k'_{9.1} \times CLASS_{VI}\_KEAP1\_NRF2_{open2\_cytosol}$ |
| *4* | Cytosolic dual ClassVI-bound KEAP1 dimer | $\frac{dCLASS_{VI2}\_KEAP1_{cytosol}}{dt} = k'_7 \times CLASS_{VI} \times CLASS_{VI}\_KEAP1_{\_cytosol} - k'_8 \times CLASS_{VI2}\_KEAP1_{\_cytosol}$ |
| *5* | Cytosolic intermediate open-state KEAP1-NRF2 complex | $\frac{dKEAP1\_NRF2_{open1\_cytosol}}{dt} = 2 \times k_1 \times KEAP1_{free\_cytosol} \times NRF2_{free\_cytosol} - k_2 \times KEAP1\_NRF2_{open1\_cytosol} - k_{1.1} \times KEAP1\_NRF2_{open1\_cytosol} + k_{2.1} \times KEAP1\_NRF2_{open2\_cytosol} - k_7 \times CLASS_{VI} \times KEAP1\_NRF2_{open1\_cytosol} + k_8 \times CLASS_{VI}\_KEAP1\_NRF2_{open1\_cytosol} - k_9 \times KEAP1\_NRF2_{open1\_cytosol}$ |
| *6* | Cytosolic final open-state KEAP1-NRF2 complex | $\frac{dKEAP1\_NRF2_{open2\_cytosol}}{dt} = k_{1.1} \times KEAP1\_NRF2_{open1} - k_{2.1} \times KEAP1\_NRF2_{open2\_cytosol} - k_3 \times KEAP1\_NRF2_{open2\_cytosol} + k_4 \times KEAP1\_NRF2_{closed\_cytosol} - k_7 \times CLASS_{VI} \times KEAP1\_NRF2_{open2\_cytosol} + k_8 \times CLASS_{VI}\_KEAP1\_NRF2_{open2\_cytosol} - k_{9.1} \times KEAP1\_NRF2_{open2\_cytosol}$ |
| *7* | Cytosolic closed-state KEAP1-NRF2 complex | $\frac{dKEAP1\_NRF2_{closed\_cytosol}}{dt} = k_3 \times KEAP1\_NRF2_{open2\_cytosol} - k_4 \times KEAP1\_NRF2_{closed\_cytosol} - k_6 \times KEAP1\_NRF2_{closed\_cytosol}$ |
| *8* | Cytosolic intermediate open-state ClassVI-KEAP1-NRF2 complex | $\frac{dCLASS_{VI}\_KEAP1\_NRF2_{open1\_cytosol}}{dt} = k'_1 \times CLASS_{VI}\_KEAP1_{\_cytosol} \times NRF2_{free\_cytosol} - k'_2 \times CLASS_{VI}\_KEAP1\_NRF2_{open1\_cytosol} - k'_{1.1} \times CLASS_{VI}\_KEAP1\_NRF2_{open1\_cytosol} + k'_{2.1} \times CLASS_{VI}\_KEAP1\_NRF2_{open2\_cytosol} + k_7 \times CLASS_{VI} \times KEAP1\_NRF2_{open1\_cytosol} - k_8 \times CLASS_{VI}\_KEAP1\_NRF2_{open1\_cytosol} - k'_9 \times CLASS_{VI}\_KEAP1\_NRF2_{open1\_cytosol}$ |
| *9* | Cytosolic intermediate open-state ClassVI-KEAP1-NRF2 complex | $\frac{dCLASS_{VI}\_KEAP1\_NRF2_{open2\_cytosol}}{dt} = k'_{1.1} \times CLASS_{VI}\_KEAP1\_NRF2_{open1\_cytosol} - k'_{2.1} \times CLASS_{VI}\_KEAP1\_NRF2_{open2\_cytosol} + k_7 \times CLASS_{VI} \times KEAP1\_NRF2_{open2\_cytosol} - k_8 \times CLASS_{VI}\_KEAP1\_NRF2_{open2\_cytosol} - k'_{9.1} \times CLASS_{VI}\_KEAP1\_NRF2_{open2\_cytosol}$ |
| *10* | Nuclear free NRF2 | $\frac{dNRF2_{free\_nucleus}}{dt} = k_{10} \times NRF2_{free\_cytosol} \times \frac{V_c}{V_n} - k_{11} \times NRF2_{free\_nucleus} - k_{n5} \times NRF2_{free\_nucleus} - 2 \times k_{n1} \times NRF2_{free\_nucleus} \times KEAP1_{free\_nucleus} + k_{n2} \times KEAP1\_NRF2_{open1\_nucleus}$ |
| *11* | Nuclear free KEAP1 dimer | $\frac{dKEAP1_{free\_nucleus}}{dt} = - 2 \times k_{n1} \times NRF2_{free\_nucleus} \times KEAP1_{free\_nucleus} + k_{n2} \times KEAP1\_NRF2_{open1\_nucleus} + k_{n6} \times KEAP1\_NRF2_{closed\_nucleus} + k_{n9} \times KEAP1\_NRF2_{open1\_nucleus} + k_{n9.1} \times KEAP1\_NRF2_{open2\_nucleus}$ |

| | | | | |
|---|---|---|---|---|
| *12* | Nuclear intermediate open-state KEAP1-NRF2 complex | $\frac{dKEAP1\_NRF2_{open1\_nucleus}}{dt}$ | = | $2 \times k_{n1} \times NRF2_{free\_nucleus} \times KEAP1_{free\_nucleus} - k_{n2} \times KEAP1\_NRF2_{open1\_nucleus} - k_{n1.1} \times KEAP1\_NRF2_{open1\_nucleus} + k_{n2.1} \times KEAP1\_NRF2_{open2\_nucleus} - k_{n9} \times KEAP1\_NRF2_{open1\_nucleus}$ |
| *13* | Nuclear final open-state KEAP1-NRF2 complex | $\frac{dKEAP1\_NRF2_{open2\_nucleus}}{dt}$ | = | $k_{n1.1} \times KEAP1\_NRF2_{open1\_nucleus} - k_{n2.1} \times KEAP1\_NRF2_{open2\_nucleus} - k_{n3} \times KEAP1\_NRF2_{open2\_nucleus} + k_{n4} \times KEAP1\_NRF2_{closed\_nucleus} - k_{n9.1} \times KEAP1\_NRF2_{open2\_nucleus}$ |
| *14* | Nuclear closed-state KEAP1-NRF2 complex | $\frac{dKEAP1\_NRF2_{closed\_nucleus}}{dt}$ | = | $k_{n3} \times KEAP1\_NRF2_{open2\_nucleus} - k_{n4} \times KEAP1\_NRF2_{closed\_nucleus} - k_{n6} \times KEAP1\_NRF2_{closed\_nucleus}$ |

**Table S7. Algebraic equations**

| | | | |
|---|---|---|---|
| **Cytosolic total NRF2** | | | |
| Models 1-2: | $NRF2_{tot\_cytosol}$ | = | $NRF2_{free\_cytosol} + KEAP1\_NRF2_{open\_cytosol} + KEAP1_{o}\_NRF2_{open\_cytosol} + KEAP1\_NRF2_{closed\_cytosol} + KEAP1_{o}\_NRF2_{closed\_cytosol}$ |
| Models 3a-4b: | $NRF2_{tot\_cytosol}$ | = | $NRF2_{free\_cytosol} + KEAP1\_NRF2_{open1\_cytosol} + KEAP1_{o}\_NRF2_{open1\_cytosol} + KEAP1\_NRF2_{open2\_cytosol} + KEAP1_{o}\_NRF2_{open2\_cytosol} + KEAP1\_NRF2_{closed\_cytosol} + KEAP1_{o}\_NRF2_{closed\_cytosol}$ |
| **Nuclear total NRF2** | | | |
| Models 4a-4b: | $NRF2_{tot\_\_nucleus}$ | = | $NRF2_{free\_\_nucleus} + KEAP1\_NRF2_{open1\_\_nucleus} + KEAP1\_NRF2_{open2\_\_nucleus} + KEAP1\_NRF2_{closed\_\_nucleus}$ |
| **Cellular total NRF2** | | | |
| Models 4a-4b: | $NRF2_{tot\_cell}$ | = | $\dfrac{NRF2_{tot\_cytosol} \times V_c + NRF2_{tot\_\_nucleus} \times V_n}{V_c + V_n}$ |
| **Cytosolic free KEAP1 dimer** | | | |
| Models 1, 2, 3a, 4a: | $KEAP1_{free\_cytosol}$ | = | $KEAP1_{free\_cytosol} + KEAP1_{o\_free\_cytosol}$ |
| **Cytosolic total open-state KEAP1-NRF2 complex** | | | |
| Models 1-2: | $KEAP1\_NRF2_{open\_tot\_cytosol}$ | = | $KEAP1\_NRF2_{open\_cytosol} + KEAP1_{o}\_NRF2_{open\_cytosol}$ |
| Models 3a and 4a: | $KEAP1\_NRF2_{open\_tot\_cytosol}$ | = | $KEAP1\_NRF2_{open1\_cytosol} + KEAP1_{o}\_NRF2_{open1\_cytosol} + KEAP1\_NRF2_{open2\_cytosol} + KEAP1_{o}\_NRF2_{open2\_cytosol}$ |
| Models 3b and 4b: | $KEAP1\_NRF2_{open\_tot\_cytosol}$ | = | $KEAP1\_NRF2_{open1\_cytosol} + CLASS_{VI}\_KEAP1\_NRF2_{open1\_cytosol} + KEAP1\_NRF2_{open2\_cytosol} + CLASS_{VI}\_KEAP1\_NRF2_{open2\_cytos}$ |
| **Cytosolic total closed-state KEAP1-NRF2 complex** | | | |
| Models 1, 2, 3a, 4a: | $KEAP1\_NRF2_{closed\_tot\_cytosol}$ | = | $KEAP1\_NRF2_{closed\_cytosol} + KEAP1_{o}\_NRF2_{closed\_cytosol}$ |
| Models 3b and 4b: | $KEAP1\_NRF2_{closed\_tot\_cytosol}$ | = | $KEAP1\_NRF2_{closed\_cytosol}$ |

**Table S8. Basal steady-state concentrations of NRF2 and KEAP1 species (nM)**

| Model # | $KEAP1_{free}$ | $NRF2_{free}$ | $KEAP1\_NRF2_{open1\_tot}$ | $KEAP1\_NRF2_{open2\_tot}$ | $KEAP1\_NRF2_{open\_tot}$ | $KEAP1\_NRF2_{closed\_tot}$ | $KEAP1_{tot}$ | $NRF2_{tot}$ |
|---|---|---|---|---|---|---|---|---|
| | | | | **Cytosol** | | | | |
| **1-2** | 382 | 2 | na | na | 74 | 74 | 530 | 150 |
| **3a-4b** | 386 | 6.2 | 58 | 14 | 72 | 72 | 530 | 150 |
| | | | | **Nucleus** | | | | |
| **4a-4b** | 2.6 | 180 | 11.5 | 14 | 25.5 | 72 | 100 | 278 |

Note: na: not applicable. Due to rounding, values may not add up exactly.

**Table S9. Maximally induced steady-state concentrations of NRF2 and KEAP1 species (nM)**

| Model # | $KEAP1_{free}$ | $NRF2_{free}$ | $KEAP1\_NRF2_{open1\_tot}$ | $KEAP1\_NRF2_{open2\_tot}$ | $KEAP1\_NRF2_{open\_tot}$ | $KEAP1\_NRF2_{closed\_tot}$ | $KEAP1_{tot}$ | $NRF2_{tot}$ |
|---|---|---|---|---|---|---|---|---|
| | | | | **Cytosol** | | | | |
| **1** | 11.2 | 231 | na | na | 258 | 260 | 530 | 750 |
| **2** | 2.82 | 223 | na | na | 63 | 465 | 530 | 750 |
| **3a** | 6.62 | 227 | 36 | 80 | 116 | 407 | 530 | 750 |
| **3b** | 2.65E-8 | 519 | 0.033 | 0.187 | 0.22 | 9.46E-6 | 530 | 519 |
| **4a** | 68 | 19 | 32 | 70 | 102 | 359 | 530 | 481 |
| **4b** | 2.65E-8 | 33 | 2.13E-3 | 0.012 | 0.014 | 6.02E-7 | 530 | 33.03 |
| | | | | **Nucleus** | | | | |
| **4a** | 0.77 | 626 | 12 | 14 | 26 | 73 | 100 | 725 |
| **4b** | 0.45 | 1079 | 12 | 14 | 26 | 73 | 100 | 1178 |

Note: Maximal induction is achieved by setting $CLASS_{I\text{-}V}$ or $CLASS_{VI}$ to 10E6. na: not applicable. Due to rounding, values may not add up exactly.

**Table S10. Basal turnover fluxes of KEAP1-NRF2 models (nM/S)**

| Model # | $flux_{k0}$ | $flux_{k1}$ | $flux_{k2}$ | $flux_{k1.1}$ | $flux_{k2.1}$ | $flux_{k3}$ | $flux_{k4}$ | $flux_{k5}$ | $flux_{k6}$ | $flux_{k9}$ | $flux_{k9.1}$ | $flux_{k10}$ | $flux_{k11}$* |
|---|---|---|---|---|---|---|---|---|---|---|---|---|---|
| **1** | 0.172 | 21.078 | 20.906 | na | na | 14.645 | 14.495 | 5.65E-4 | 0.1501 | 0.0214 | na | na | na |
| **2** | 0.15 | 21.116 | 20.966 | na | na | 0.1353 | 7.35E-3 | 5.66E-4 | 0.128 | 0.0215 | na | na | na |
| **3a-3b** | 0.15 | 16.482 | 16.334 | 0.1332 | 1.73E-3 | 14.194 | 14.066 | 1.78E-3 | 0.1274 | 1.67E-3 | 4.1E-3 | na | na |
| **4a-4b** | 0.1933 | Same as Models 3a-3b | | | | | | | | | | 0.1185 | 0.0751 |

Note: *: normalized to cytosolic volume; na: not applicable. Due to rounding, values may not add up exactly.

**Table S11. Maximally-induced turnover fluxes of KEAP1-NRF2 models (nM/S)**

| Model # | $flux_{k0}$ | $flux_{k1}$ + $flux_{k'1}$ | $flux_{k2}$ + $flux_{k'2}$ | $flux_{k1.1}$ + $flux_{k'1.1}$ | $flux_{k2.1}$ + $flux_{k'2.1}$ | $flux_{k3}$ + $flux_{k'3}$# | $flux_{k4}$ + $flux_{k'4}$# | $flux_{k5}$ | $flux_{k6}$ | $flux_{k'6}$ | $flux_{k9}$ + $flux_{k'9}$ | $flux_{k9.1}$ + $flux_{k'9.1}$ | $flux_{k10}$ | $flux_{k11}$* |
|---|---|---|---|---|---|---|---|---|---|---|---|---|---|---|
| **1** | 0.172 | 72.99 | 72.89 | na | na | 51.06 | 51.03 | 0.067 | 5.3E-6 | 0.031 | 0.075 | na | na | na |
| **2** | 0.15 | 17.75 | 17.66 | na | na | 0.114 | 0.046 | 0.064 | 8.1E-6 | 0.068 | 0.018 | na | na | na |
| **3a** | 0.15 | 10.36 | 10.27 | 0.084 | 9.7E-3 | 79.84 | 79.79 | 0.065 | 7.2E-6 | 0.051 | 0.011 | 0.023 | na | na |
| **3b** | 0.15 | 9.5E-3 | 9.4E-3 | 7.7E-5 | 2.3E-5 | 1.87E-6 | 1.85E-6 | 0.15 | 1.7E-8 | na | 9.7E-6 | 5.4E-5 | na | na |
| **4a** | 0.1933 | 9.141 | 9.067 | 0.074 | 8.6E-3 | 70.48 | 70.43 | 5.6E-3 | 6.4E-6 | 0.045 | 9.3E-3 | 0.02 | 0.373 | 0.26 |
| **4b** | 0.1933 | 6.0E-4 | 6.0E-4 | 4.9E-6 | 1.5E-6 | 1.19E-7 | 1.18E-7 | 9.5E-3 | 1.0E-9 | na | 6.1E-7 | 3.4E-6 | 0.632 | 0.449 |

Note: Maximal induction is achieved by setting $CLASS_{I\text{-}V}$ or $CLASS_{VI}$ to 10E6. #: $flux_{k'3}$ and $flux_{k'3}$ do not exist for Models 3b and 4b; *: normalized to cytosolic volume; na: not applicable. Due to rounding, values may not add up exactly.
