## Supplementary material for "Mathematical Modeling Reveals Quantitative Properties of KEAP1-NRF2 Signaling": Model Code in Berkeley Madonna and R: Model 3a.R

```
### {Model 3a: Two-step ETGE binding KEAP1-NRF2 model for Class I-V activators}
#Unit: time = second, concentration = nM

# Load Package
library(deSolve)

## ---------------------------------
## Parameter values
## --------------------------------- 
parameters <- c(
  ClassI_V 		= 0,
  k0 		= 0.15,
  k1		= 3.45e-3,
  kprime1		= 3.45e-3,
  k2		= 0.282,
  kprime2		= 0.282,
  k1dot1		= 0.0023,
  kprime1dot1	= 0.0023,
  k2dot1		= 1.22e-4,
  kprime2dot1	= 1.22e-4,
  k3		= 1.0,
  kprime3		= 1.0,
  k4		= 0.196,
  kprime4		= 0.196,
  k5		= log(2)/(40*60),
  k6		= 1.775e-3,
  kprime6		= 1.252e-4,
  k7		= 0.01,
  k8		= 0.1,
  k9		= log(2)/(40*60),
  kprime9		= log(2)/(40*60),
  k9dot1		= log(2)/(40*60),
  kprime9dot1	= log(2)/(40*60)
)

## ----------------------------------
## Initial conditions
## ----------------------------------
y0 <- c(
  NRF2free 			= 0,
  KEAP1free			= 530,
  KEAP1o_free		= 0,
  KEAP1_NRF2open1 		= 0,
  KEAP1o_NRF2open1 	= 0,
  KEAP1_NRF2open2 		= 0,
  KEAP1o_NRF2open2 	= 0,
  KEAP1_NRF2closed 	= 0,
  KEAP1o_NRF2closed 	= 0
)

## ----------------------------------
## Differential equations to solve
## ----------------------------------
Model_3a <- function(times, y, parms)
{
  with(as.list(c(y, parms)),
       {
         #ODEs for the state variables
         dNRF2freedt		= k0 - k5 * NRF2free - 2 * k1 * KEAP1free * NRF2free + k2 * KEAP1_NRF2open1 - 2 * kprime1 * KEAP1o_free * NRF2free + kprime2 * KEAP1o_NRF2open1
         dKEAP1freedt		= - 2 * k1 * KEAP1free * NRF2free + k2 * KEAP1_NRF2open1 - k7 * KEAP1free * ClassI_V + k8 * KEAP1o_free + k6 * KEAP1_NRF2closed + k9 * KEAP1_NRF2open1 + k9dot1 * KEAP1_NRF2open2
         dKEAP1o_freedt		= - 2 * kprime1 * KEAP1o_free * NRF2free + kprime2 * KEAP1o_NRF2open1 + k7 * KEAP1free * ClassI_V - k8 * KEAP1o_free + kprime6 * KEAP1o_NRF2closed + kprime9 * KEAP1o_NRF2open1 + kprime9dot1 * KEAP1o_NRF2open2
         dKEAP1_NRF2open1dt	= 2 * k1 * KEAP1free * NRF2free - k2 * KEAP1_NRF2open1 - k1dot1 * KEAP1_NRF2open1 + k2dot1 * KEAP1_NRF2open2 - k7 * ClassI_V * KEAP1_NRF2open1 + k8 * KEAP1o_NRF2open1 - k9 * KEAP1_NRF2open1
         dKEAP1o_NRF2open1dt	= 2 * kprime1 * KEAP1o_free * NRF2free - kprime2 * KEAP1o_NRF2open1 - kprime1dot1 * KEAP1o_NRF2open1 + kprime2dot1 * KEAP1o_NRF2open2 + k7 * ClassI_V * KEAP1_NRF2open1 - k8 * KEAP1o_NRF2open1 - kprime9 * KEAP1o_NRF2open1
         dKEAP1_NRF2open2dt	=  k1dot1 * KEAP1_NRF2open1 - k2dot1 * KEAP1_NRF2open2 - k3 * KEAP1_NRF2open2 + k4 * KEAP1_NRF2closed - k7 * ClassI_V * KEAP1_NRF2open2 + k8 * KEAP1o_NRF2open2 - k9dot1 * KEAP1_NRF2open2 
         dKEAP1o_NRF2open2dt	= kprime1dot1 * KEAP1o_NRF2open1 - kprime2dot1 * KEAP1o_NRF2open2 - kprime3 * KEAP1o_NRF2open2 + kprime4 * KEAP1o_NRF2closed + k7 * ClassI_V * KEAP1_NRF2open2 - k8 * KEAP1o_NRF2open2 - kprime9dot1 * KEAP1o_NRF2open2
         dKEAP1_NRF2closeddt	= k3 * KEAP1_NRF2open2 - k4 * KEAP1_NRF2closed - k6 * KEAP1_NRF2closed - k7 * ClassI_V * KEAP1_NRF2closed + k8 * KEAP1o_NRF2closed 
         dKEAP1o_NRF2closeddt	= kprime3 * KEAP1o_NRF2open2 - kprime4 * KEAP1o_NRF2closed + k7 * ClassI_V * KEAP1_NRF2closed - k8 * KEAP1o_NRF2closed - kprime6 * KEAP1o_NRF2closed
         #Return simulated value of Y at each time step
         list(c(dNRF2freedt, dKEAP1freedt, dKEAP1o_freedt, dKEAP1_NRF2open1dt, dKEAP1o_NRF2open1dt, dKEAP1_NRF2open2dt, dKEAP1o_NRF2open2dt, dKEAP1_NRF2closeddt, dKEAP1o_NRF2closeddt)); #They need to follow the same order as the ODEs
       })
}

## ----------------------------------
## Run simulation
## ----------------------------------
#Time span of simulation
tspan <- seq(0, 108000, by = 10)

#Call the lsoda function to numerically solve the ODEs
output <- lsoda(y = y0, times = tspan, func = Model_3a, parms = parameters)

#Optional: show the result of the first 6 time steps
head(output)

## ----------------------------------
## Plotting simulation results
## ----------------------------------
par(mar = c(5, 5, 1, 1), mgp = c(2, 0.5, 0)) # mar(figure margins):bottom, left, top, right; mgp(label position): distance from label, scale, tick marks to plot 
plot(output, xlab = "Time (S)", ylab = "conc (nM)", las=1, cex.lab=1.4, cex.axis=1.3, col="blue")
```
