## Supplementary material for "Mathematical Modeling Reveals Quantitative Properties of KEAP1-NRF2 Signaling": Model Code in Berkeley Madonna and R: Model 4a.R

```
### {Model 4a: Two-step ETGE binding KEAP1-NRF2 model with nucleus for Class I-V activators}
#Unit: time = second, concentration = nM

# Load Package
library(deSolve)

## ---------------------------------
## Parameter values
## --------------------------------- 
parameters <- c(
  ClassI_V 	= 0,
  k0		= 0.1933,
  k1		= 3.45e-3,
  kprime1		= 3.45e-3,
  k2		= 0.282,
  kprime2		= 0.282,
  k1dot1		= 0.0023,
  kprime1dot1	= 0.0023,
  k2dot1		= 1.22e-4,
  kprime2dot1	= 1.22e-4,
  k3		= 1.0,
  kprime3		= 1.0,
  k4		= 0.196,
  kprime4		= 0.196,
  k5		= log(2)/(40*60),
  k6		= 1.775e-3,
  kprime6		= 1.252e-4,
  k7		= 0.01,
  k8		= 0.1,
  k9		= log(2)/(40*60),
  kprime9		= log(2)/(40*60),
  k9dot1		= log(2)/(40*60),
  kprime9dot1	= log(2)/(40*60),
  Vc		= 1,
  Vn		= 0.54,
  k10 		= 0.01916,
  k11 		= log(2)/(15*60),
  kn1		= 3.450E-3,
  kn2		= 2.820E-1,
  kn1dot1		= 0.0023,
  kn2dot1		= 1.220E-4,
  kn3		= 1.0,
  kn4		= 1.960E-1,
  kn5		= log(2)/(40*60),
  kn6		= log(2)/(40*60),
  kn9		= log(2)/(40*60),
  kn9dot1		= log(2)/(40*60)
)

## ----------------------------------
## Initial conditions
## ----------------------------------
y0 <- c(
  NRF2free_cytosol 			= 0,
  KEAP1free_cytosol			= 530,
  KEAP1o_free_cytosol		= 0,
  KEAP1_NRF2open1_cytosol 	= 0,
  KEAP1o_NRF2open1_cytosol  	= 0,
  KEAP1_NRF2open2_cytosol 	= 0,
  KEAP1o_NRF2open2_cytosol 	= 0,
  KEAP1_NRF2closed_cytosol  	= 0,
  KEAP1o_NRF2closed_cytosol  	= 0,
  NRF2free_nucleus 			= 0,
  KEAP1free_nucleus 		= 100,
  KEAP1_NRF2open1_nucleus 	= 0,
  KEAP1_NRF2open2_nucleus 	= 0,
  KEAP1_NRF2closed_nucleus 	= 0
)

## ----------------------------------
## Differential equations to solve
## ----------------------------------
Model_4a <- function(times, y, parms)
{
  with(as.list(c(y, parms)),
       {
         #ODEs for the state variables
         dNRF2free_cytosoldt 			= k0 - k5 * NRF2free_cytosol - 2 * k1 * KEAP1free_cytosol * NRF2free_cytosol + k2 * KEAP1_NRF2open1_cytosol - 2 * kprime1 * KEAP1o_free_cytosol * NRF2free_cytosol + kprime2 * KEAP1o_NRF2open1_cytosol - k10 * NRF2free_cytosol + k11 * NRF2free_nucleus * Vn/Vc
         dKEAP1free_cytosoldt 			= - 2 * k1 * KEAP1free_cytosol * NRF2free_cytosol + k2 * KEAP1_NRF2open1_cytosol - k7 * KEAP1free_cytosol * ClassI_V + k8 * KEAP1o_free_cytosol + k6 * KEAP1_NRF2closed_cytosol  + k9 * KEAP1_NRF2open1_cytosol + k9dot1 * KEAP1_NRF2open2_cytosol 
         dKEAP1o_free_cytosoldt 		= - 2 * kprime1 * KEAP1o_free_cytosol * NRF2free_cytosol + kprime2 * KEAP1o_NRF2open1_cytosol  + k7 * KEAP1free_cytosol * ClassI_V - k8 * KEAP1o_free_cytosol + kprime6 * KEAP1o_NRF2closed_cytosol  + kprime9 * KEAP1o_NRF2open1_cytosol  + kprime9dot1 * KEAP1o_NRF2open2_cytosol 
         dKEAP1_NRF2open1_cytosoldt  	= 2 * k1 * KEAP1free_cytosol * NRF2free_cytosol - k2 * KEAP1_NRF2open1_cytosol - k1dot1 * KEAP1_NRF2open1_cytosol + k2dot1 * KEAP1_NRF2open2_cytosol  - k7 * ClassI_V * KEAP1_NRF2open1_cytosol + k8 * KEAP1o_NRF2open1_cytosol  - k9 * KEAP1_NRF2open1_cytosol
         dKEAP1o_NRF2open1_cytosoldt   	= 2 * kprime1 * KEAP1o_free_cytosol * NRF2free_cytosol - kprime2 * KEAP1o_NRF2open1_cytosol  - kprime1dot1 * KEAP1o_NRF2open1_cytosol  + kprime2dot1 * KEAP1o_NRF2open2_cytosol  + k7 * ClassI_V * KEAP1_NRF2open1_cytosol - k8 * KEAP1o_NRF2open1_cytosol  - kprime9 * KEAP1o_NRF2open1_cytosol 
         dKEAP1_NRF2open2_cytosoldt  	=  k1dot1 * KEAP1_NRF2open1_cytosol - k2dot1 * KEAP1_NRF2open2_cytosol  - k3 * KEAP1_NRF2open2_cytosol  + k4 * KEAP1_NRF2closed_cytosol  - k7 * ClassI_V * KEAP1_NRF2open2_cytosol  + k8 * KEAP1o_NRF2open2_cytosol  - k9dot1 * KEAP1_NRF2open2_cytosol 
         dKEAP1o_NRF2open2_cytosoldt  	= kprime1dot1 * KEAP1o_NRF2open1_cytosol  - kprime2dot1 * KEAP1o_NRF2open2_cytosol  - kprime3 * KEAP1o_NRF2open2_cytosol  + kprime4 * KEAP1o_NRF2closed_cytosol  + k7 * ClassI_V * KEAP1_NRF2open2_cytosol  - k8 * KEAP1o_NRF2open2_cytosol  - kprime9dot1 * KEAP1o_NRF2open2_cytosol 
         dKEAP1_NRF2closed_cytosoldt   	= k3 * KEAP1_NRF2open2_cytosol  - k4 * KEAP1_NRF2closed_cytosol  - k6 * KEAP1_NRF2closed_cytosol  - k7 * ClassI_V * KEAP1_NRF2closed_cytosol  + k8 * KEAP1o_NRF2closed_cytosol
         dKEAP1o_NRF2closed_cytosoldt   	=  kprime3 * KEAP1o_NRF2open2_cytosol  - kprime4 * KEAP1o_NRF2closed_cytosol  + k7 * ClassI_V * KEAP1_NRF2closed_cytosol  - k8 * KEAP1o_NRF2closed_cytosol  - kprime6 * KEAP1o_NRF2closed_cytosol 
         dNRF2free_nucleusdt  			= k10 * NRF2free_cytosol * Vc/Vn - k11 * NRF2free_nucleus - kn5 * NRF2free_nucleus - 2 * kn1 *  NRF2free_nucleus * KEAP1free_nucleus + kn2 * KEAP1_NRF2open1_nucleus
         dKEAP1free_nucleusdt  		= - 2 * kn1 * NRF2free_nucleus * KEAP1free_nucleus + kn2 * KEAP1_NRF2open1_nucleus + kn6 * KEAP1_NRF2closed_nucleus + kn9 * KEAP1_NRF2open1_nucleus + kn9dot1 * KEAP1_NRF2open2_nucleus
         dKEAP1_NRF2open1_nucleusdt  	= 2 * kn1 * NRF2free_nucleus * KEAP1free_nucleus - kn2 * KEAP1_NRF2open1_nucleus - kn1dot1 * KEAP1_NRF2open1_nucleus + kn2dot1 * KEAP1_NRF2open2_nucleus - kn9 * KEAP1_NRF2open1_nucleus
         dKEAP1_NRF2open2_nucleusdt  	=  kn1dot1 * KEAP1_NRF2open1_nucleus - kn2dot1 * KEAP1_NRF2open2_nucleus - kn3 * KEAP1_NRF2open2_nucleus + kn4 * KEAP1_NRF2closed_nucleus - kn9dot1 * KEAP1_NRF2open2_nucleus
         dKEAP1_NRF2closed_nucleusdt  	= kn3 * KEAP1_NRF2open2_nucleus - kn4 * KEAP1_NRF2closed_nucleus - kn6 * KEAP1_NRF2closed_nucleus
         #Return simulated value of Y at each time step
         list(c(dNRF2free_cytosoldt,dKEAP1free_cytosoldt,dKEAP1o_free_cytosoldt,dKEAP1_NRF2open1_cytosoldt,dKEAP1o_NRF2open1_cytosoldt,dKEAP1_NRF2open2_cytosoldt,dKEAP1o_NRF2open2_cytosoldt,dKEAP1_NRF2closed_cytosoldt,dKEAP1o_NRF2closed_cytosoldt,dNRF2free_nucleusdt,dKEAP1free_nucleusdt,dKEAP1_NRF2open1_nucleusdt,dKEAP1_NRF2open2_nucleusdt,dKEAP1_NRF2closed_nucleusdt)); #They need to follow the same order as the ODEs
       })
}

## ----------------------------------
## Run simulation
## ----------------------------------
#Time span of simulation
tspan <- seq(0, 108000, by = 10)

#Call the lsoda function to numerically solve the ODEs
output <- lsoda(y = y0, times = tspan, func = Model_4a, parms = parameters)

#Optional: show the result of the first 6 time steps
head(output)

## ----------------------------------
## Plotting simulation results
## ----------------------------------
par(mar = c(5, 5, 1, 1), mgp = c(2, 0.5, 0)) # mar(figure margins):bottom, left, top, right; mgp(label position): distance from label, scale, tick marks to plot 
plot(output, xlab = "Time (S)", ylab = "conc (nM)", las=1, cex.lab=1.4, cex.axis=1.3, col="blue")
```
