## Supplementary material for "Mathematical Modeling Reveals Quantitative Properties of KEAP1-NRF2 Signaling": Model Code in Berkeley Madonna and R: Reduced Model.R

```
### {Reduced KEAP1-NRF2 model}
#Unit: time = second, concentration = nM

# Load Package
library(deSolve)

## ---------------------------------
## Parameter values
## --------------------------------- 
parameters <- c(
  ClassI_V = 0,
  ClassVI = 0,
  k0 	= 0.15,
  kf 	= 0.01,
  kb 	= 0.0001,
  k1 	= 2.888E-4,
  k2 	= 1.775E-3,
  k3 	= 0.14,
  k4 	= 7.702E-4,
  k5 	= 2.888E-4,
  Kd1 	= 7,
  Kd2 	= 2,
  KEAP1_tot = 200
)

## ----------------------------------
## Initial conditions
## ----------------------------------
y0 <- c(
  NRF2 	= 0,
  KEAP1 	= 200,
  KEAP1_NRF2 = 0,
  NRF2n 	= 0
)

## ----------------------------------
## Differential equations to solve
## ----------------------------------
Model_Reduced_KEAP1_NRF2 <- function(times, y, parms)
{
  with(as.list(c(y, parms)),
       {
         #ODEs for the state variables
         dNRF2dt 		= k0 - Kd2/(Kd2+ClassVI)*kf*KEAP1*NRF2 + kb*KEAP1_NRF2 - k1*NRF2 - k3*NRF2 + k4*NRF2n
         dKEAP1dt  	= - Kd2/(Kd2+ClassVI)*kf*KEAP1*NRF2 + kb*KEAP1_NRF2 + Kd1/(Kd1+ClassI_V)*k2*KEAP1_NRF2
         dKEAP1_NRF2dt  	= Kd2/(Kd2+ClassVI)*kf*KEAP1*NRF2 - kb*KEAP1_NRF2 - Kd1/(Kd1+ClassI_V)*k2*KEAP1_NRF2
         dNRF2ndt  		= k3*NRF2 - k4*NRF2n - k5*NRF2n
         #Return simulated value of Y at each time step
         list(c(dNRF2dt,dKEAP1dt,dKEAP1_NRF2dt,dNRF2ndt)); #They need to follow the same order as the ODEs
       })
}

## ----------------------------------
## Run simulation
## ----------------------------------
#Time span of simulation
tspan <- seq(0, 108000, by = 10)

#Call the lsoda function to numerically solve the ODEs
output <- lsoda(y = y0, times = tspan, func = Model_Reduced_KEAP1_NRF2, parms = parameters)

#Optional: show the result of the first 6 time steps
head(output)

## ----------------------------------
## Plotting simulation results
## ----------------------------------
par(mar = c(5, 5, 1, 1), mgp = c(2, 0.5, 0)) # mar(figure margins):bottom, left, top, right; mgp(label position): distance from label, scale, tick marks to plot 
plot(output, xlab = "Time (S)", ylab = "conc (nM)", las=1, cex.lab=1.4, cex.axis=1.3, col="blue")
```
